# Reconstructing and Analysing The Genome of The Last Eukaryote Common Ancestor to Better Understand the Transition from FECA to LECA

**DOI:** 10.1101/538264

**Authors:** David Newman, Fiona J. Whelan, Matthew Moore, Martin Rusilowicz, James O. McInerney

## Abstract

It is still a matter of debate whether the First Eukaryote Common Ancestor (FECA) arose from the merger of an archaeal host with an alphaproteobacterium, or was a proto-eukaryote with significant eukaryotic characteristics way before endosymbiosis occurred. The Last Eukaryote Common Ancestor (LECA) as its descendant is thought to be an entity that possessed functional and cellular complexity comparable to modern organisms. The precise nature and physiology of both of these organisms has been a long-standing, unanswered question in evolutionary and cell biology. Recently, a much broader diversity of eukaryotic genomes has become available and this means we can reconstruct early eukaryote evolution with a greater deal of precision. Here, we reconstruct a hypothetical genome for LECA from modern eukaryote genomes. The constituent genes were mapped onto 454 pathways from the KEGG database covering cellular, genetic, and metabolic processes across six model species to provide functional insights into it’s capabilities. We reconstruct a LECA that was a facultatively anaerobic, single-celled organism, similar to a modern Protist possessing complex predatory and sexual behaviour. We go on to examine how much of these capabilities arose along the FECA-to-LECA transition period. We see a at least 1,554 genes gained by FECA during this evolutionary period with extensive remodelling of pathways relating to lipid metabolism, cellular processes, genetic information processing, protein processing, and signalling. We extracted the BRITE classifications for the genes from the KEGG database, which arose during the transition from FECA-to-LECA and examine the types of genes that saw the most gains and what novel classifications were introduced. Two-thirds of our reconstructed LECA genome appears to be prokaryote in origin and the remaining third consists of genes with functional classifications that originate from prokaryote homologs in our LECA genome. Signal transduction and Post Translational Modification elements stand out as the primary novel classes of genes developed during this period. These results suggest that largely the eukaryote common ancestors achieved the defining characteristics of modern eukaryotes by primarily expanding on prokaryote biology and gene families.

## Introduction

Ernst Mayr proposed that the differences between prokaryotes and eukaryotes are the biggest phenotypic split in all of cellular life ^1^. The formation of the eukaryotic cell constitutes a major transition in life’s history, and as such is a major focus for study in evolutionary biology ^2-4^. Recently the Archaean Supergroup, the Asgard Archaea, have challenged much of our understanding about how distinct eukaryotes and prokaryotes are, consistently showing a robust phylogenetic affiliation with eukaryotes ^5-7^. Many proteins that until recently were thought to be eukaryote-specific now seem to have homologs in these Asgard genomes. Specifically, components of the ESCRT system, the TRAPP membrane-trafficking systems, the ubiquitin modifier system, the actin cytoskeleton, and an expanded range of GTPases have been found in the metagenomic assembles that identified the Asgardarchaeota species ^6-8^. Increasingly characteristic structures and biology previously used to characterize eukaryotes appear to have its origins from within this archaeal group.

While the Asgard Archaea challenge the distinctiveness of many of the characteristic elements of the eukaryote lineage, the eukaryotic cell still possesses many notable unique features. For example, the eukaryotic cell, on average, is 1,000-fold larger by volume than bacterial and archaeal cells requiring it to be governed by different physical principles; prokaryotes can rely on free diffusion for intracellular transport but eukaryotic cells possess elaborate cytoskeleton and endomembrane systems ^9-12^. Eukaryotes have also compartmentalized their nuclear genetic material, their mitochondria and mitochondrial related organelles (MROs), and their cytoplasm ^13-18^. The organizational and structural complexity that eukaryote cells exhibit is accompanied by a multitude of sophisticated signaling networks, including the kinase-phosphatase and ubiquitin systems ^19-26^. Eukaryote gene expression is distinct from Prokaryote mechanisms with the transcriptional regulation of individual genes occurring separately from their translation - regulated by microRNAs - and epigenetic gene regulation via chromatin remodeling ^13,27-32^.

In her seminal paper on endosymbiotic theory, Margulis proposed that the Eukaryotic lineage was established from the merger of an archaeal host with a α-proteobacterium, giving rise to the First Eukaryote Common Ancestor (FECA) ^33^. This bacterial symbiont would give rise to the eukaryotic mitochondria and MROs, a theory supported by evidence that these organelles are monophyletic ^34^. Modern hypotheses largely accept the central role of mitochondrial endosymbiosis, though disagreement on whether this occurred early or late during eukaryogenesis persists ^34^. Mito-early hypotheses argue that the mitochondria emerged early (or first) in the process of eukaryogenesis, while mito-late hypotheses posit mitochondria were incorporated after some or much of eukaryote complexity was acquired. While these hypotheses may agree on the nature of the mitochondrial ancestor, the nature of the host that engulfed it is hotly contested. Typically, mito-early hypotheses assume an archaeal host, whereas mito-late hypotheses tend to posit a host with some eukaryotic features, sometimes referred to as a “proto-eukaryote” ^35,36^. The level of complexity of this purported proto-eukaryote varies widely among different hypotheses. The archezoa hypothesis argued amitochondriate eukaryotes were primitive and eukaryotic complexity was acquired before endosymbiosis, with early molecular phylogenies placing amitochondriate eukaryotes as early branching lineages in the tree of eukaryotes supporting this proposition ^37^. This evidence was later found to be a phylogenetic artefact, and mitochrondrially derived genes are present in the nuclear genomes of the supposed amitochondriate archezoa species ^38^. But with the increasing body of literature on the surprising eukaryote-like properties of the Asgard Archaea ^5-7,8^, it increasingly appears that prokaryote complexity has been consistently underestimated. For example, while members of the Planctomycetes, Verrucomicrobiae, and Chlamydiae (PVC) bacterial Superphylum don’t have true compartmentalized nor nucleated cells they demonstrate a striking level of intracellular structural complexity and there are other examples of true membrane-bound vesicles in prokaryotes ^39,40^.

With recent advances in DNA sequencing technology, an expanded repertoire of eukaryotic genomes has become available. Consequently, we can trace ancestral states in the eukaryotic lineage more accurately than ever before. In this paper, we have used this expanded diversity of genomes, in combination with homolog identification across all of life to create a parsimonious reconstruction of ancestral character states in order to reconstruct the LECA genome. We have used this reconstructed genome to uncover crucial aspects of the biology of eukaryotes during the period where FECA became LECA. By reconstructing this genome of the LECA followed by tracing the elements that arose during the transition from First to Last Common Ancestor we attempt to better understand what molecular functions and biological processes were developed and expanded to make eukaryotes so unique.

## Results

### Reconstruction of LECA Pathways

We identified a total of 4,462 gene clusters that included (a) a broad taxonomic distribution across eukaryotes, and (b) that formed a monophyletic group on a phylogeny inferred from the gene sequences (herein referred to as monophyletic clusters). If we include gene clusters with a broad distribution of homologs among eukaryotes but whose genes were not monophyletic, we recover a total of 1,476 additional homologous families (herein referred to as clusters with complex history) making a sum total of 5,938 clusters. Both datasets were mapped onto 454 pathways from the KEGG database covering cellular processes, genetic information processing, environmental information processing, and metabolic pathways across six model species (*Homo sapiens, Saccharomyces cerevisiae, Arabidopsis thaliana, Dictyostelium discoideum, Methanobrevibacter smithii*, and *Escherichia coli*) (SI Figures S1-S5). These gene clusters along with their FASTA gene annotations, and their matches to KEGG components and corresponding KEGG pathways are available to the interested reader as a neo4j network (SI Data 1). We have created a series of heatmaps of pathway completeness for each model organism used in the analyses that summarises the breadth of the data contained in the network (Figure 1).

**Figure 1.**
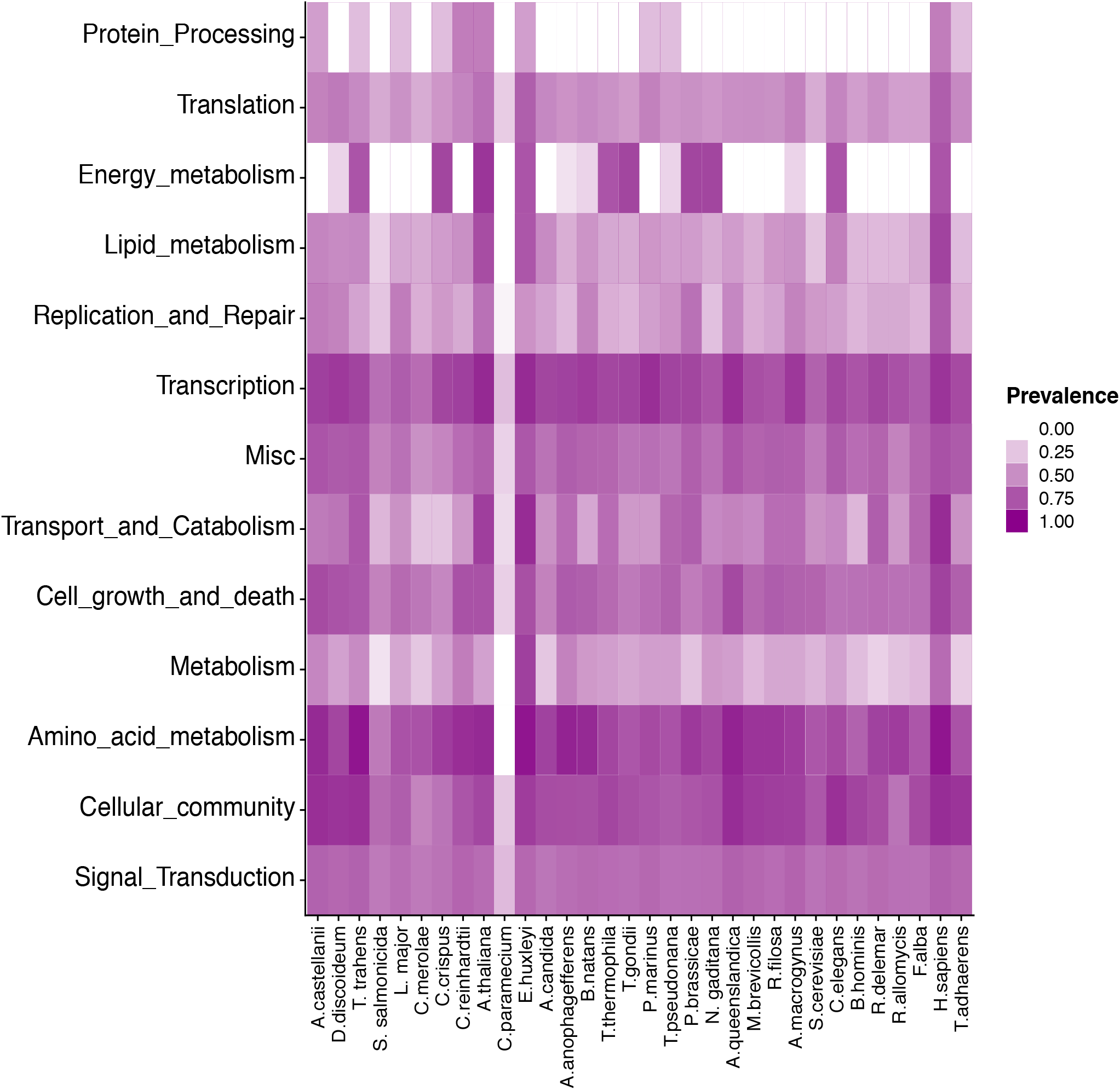
Heatmap of pathway completeness for human pathways across each of the constituent genomes of the reconstructed LECA genome, additional heatmaps detailing pathway completeness for the other model species can be found in the supplementary information (Figure S6). KEGG pathways categorised into groups of similar or linked function. *C. paramecium* represents a minimalist nucleomorph genome that lacks it’s own metabolic genes.

We analysed a variety of pathways of interest including central metabolism, as well as many pathways involved in membrane biology, cell division, the spliceosome, endocytosis, and the phagosome (Figure 1, SI Figures S1-S5). These analyses confirmed what many previous studies have shown, proposed, and inferred about the biology and complexity of LECA ^41^. Our reconstruction indicates that LECA possessed largely the same metabolic capacity to process glucose into pyruvate, oxidize acetate into CO_2_ and water, and then to produce ATP from NADH through electron transport as the majority of modern eukaryotes (SI Figures S1-S5). Our reconstruction also suggests that mitosis, recombination, and sexual reproduction were all present in LECA in a form not dissimilar to modern mechanisms (SI Figures S1-S5). Our LECA reconstruction has a largely intact set of membrane biosynthesis pathways, as well as the capacity to manipulate these membranes for the purposes of phagocytosis and endocytosis (SI Figures S1-S5). Overall our reconstruction strongly supports the idea, and the growing body of evidence that LECA was a complex organism that would have more or less resembled a modern protist ^41^.

### Notable Expansion of Genes during The FECA-to-LECA transition

Becoming a recognisable eukaryote took some time and in this span between the first and last common ancestor of eukaryotes, many biological functions were invented ^42,43^. By assessing our reconstruction to identify genes present in LECA that lack prokaryote homologues, we can identify the biological functions that arose during the FECA-to-LECA transition period. This subset of our data, independently crossed-referenced against the taxonomic gene tree data in the KEGG database, reveals 1,554 genes involved in 212 KEGG pathways across four species, comprising 95 unique pathways that were biological innovations of the FECA-to-LECA transition period. These pathways have been organised into eight categories based on their KEGG classifications: Energy Metabolism; Lipid Metabolism; Miscellaneous Metabolism; Cellular Processes; Genetic Information Processing; Protein Processing; Signalling; and Amino Acid and Nucleotide Metabolism (Figure 2). The categories showing the largest number of elements introduced during this time period are Cellular Processes, Genetic Information Processing, Protein Processing, Lipid Metabolism, and Signalling (SI Table 2). As one may predict, these results indicate that the majority of the changes in biology relate to membrane biology, and how it was used to create distinct internal environments and structures within the cell, as well as systems to regulate the interaction between these structures and complex behaviours.

**Figure 2.**
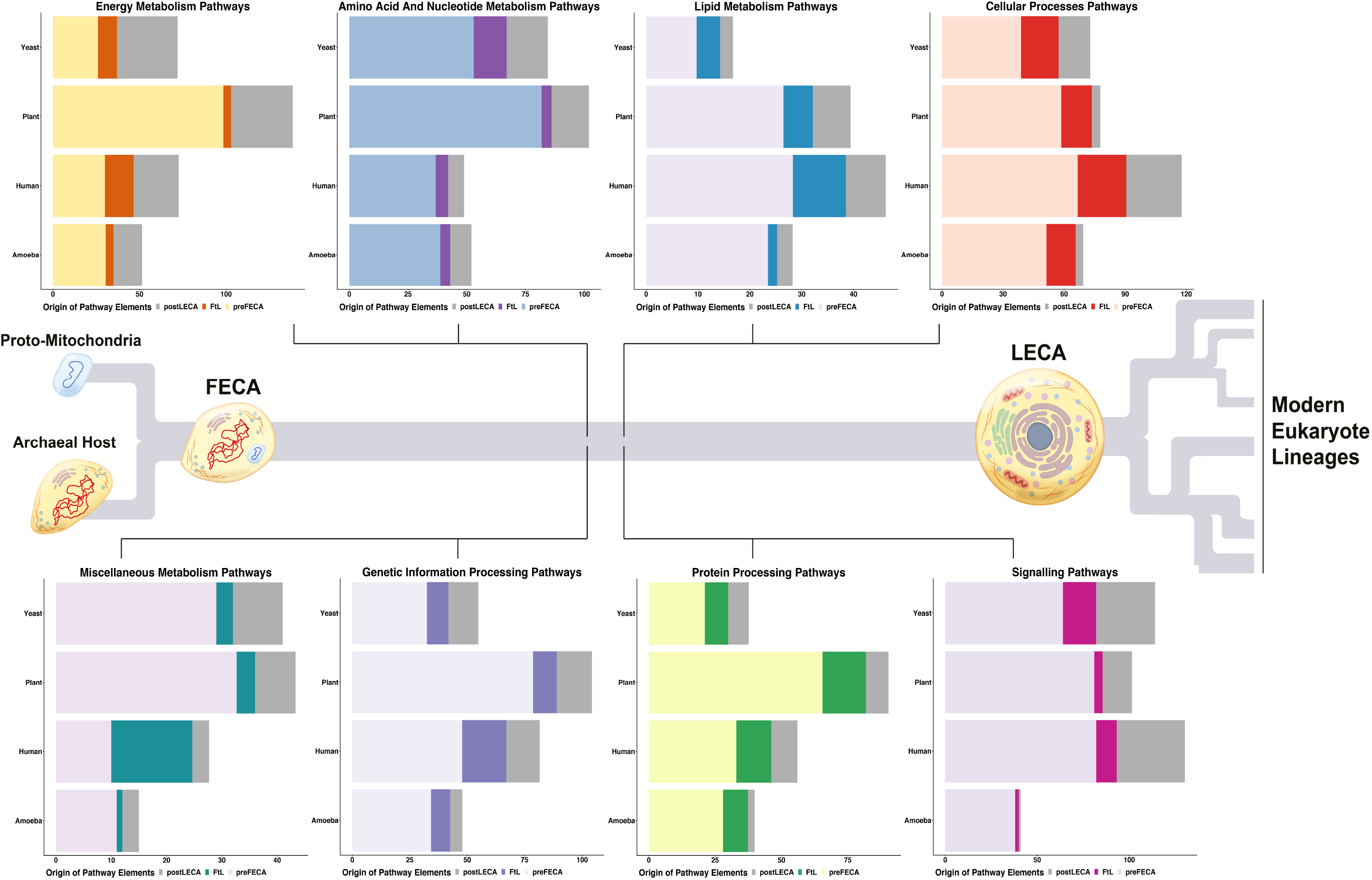
Diagrammatic schemata of the development from FECA to LECA and the gene acquisition amongst genetic, metabolic and cellular process related pathways. This Figure shows the mean gene gain per pathway across the eight categories of pathways that see gain gains during the evolutionary time period between the first and last common ancestor, for the pathways of four eukaryote model species, amoeba, human, plant, and yeast. FtL = FECA-to-LECA transition.

### Substantial Increases in the number of genes do not show a corresponding substantial increase in functional novelty

The KEGG database employs a hierarchical system of functional annotations for genes and other biological entities, called BRITE ^89^. We extracted these BRITE classifications for the genes, which arose between FECA and LECA. While the most common was the generic designation “Enzymes”, the other major groups of genes belonged to the following classifications: Membrane trafficking, Ubiquitin system, Exosome, Protein kinases, Messenger RNA biogenesis, Chromosome and associated proteins, and DNA repair and recombination proteins (Figure 3). Membrane trafficking as a major area of expansion is a predictable result, as organelles and the endomembrane system developed substantially between our nearest archaeal ancestor and modern eukaryotes, and transport between these subcompartments became necessary ^44^. The Ubiquitin system co-ordinates a large number of processes through targeting proteins for degradation, and is of primary interest in this context as it has roles in the control of signal transduction pathways, transcriptional regulation, and endocytosis ^19,21^. Protein Kinases are well-studied signalling components related to a large number of relevant biological processes of early eukaryotes; particularly intracellular signaling, nuclear transcription, translocation of transcription factors, and nuclear receptors ^25,45^. Alongside the membrane trafficking and signal transduction elements, the remaining four functional classes relate to genetic information processing tasks. We see increases in genes involved in Messenger RNA biogenesis, as mRNA became required to be moved from discrete locations within the cell, from the nucleus to the cytoplasm, and systems arose to facilitate this ^46^. RNA quality control mechanisms were also further expanded upon within the lineage, including the exosome, a highly conserved complex responsible for RNA processing ^47^. DNA repair and recombination proteins increased in number; we see both an expansion of repair machinery and mechanisms ^48-50^, but also that early eukaryotes were required to interact with chromatin, accounting for the increase in chromosome and associated proteins that is also observed, in order to accomplish these tasks ^51^. Dealing with the consequences of subcompartmentalising biology and its increasing cellular size appears to have been the driving cause of many of these genetic additions to the early eukaryote lineage.

**Figure 3.**
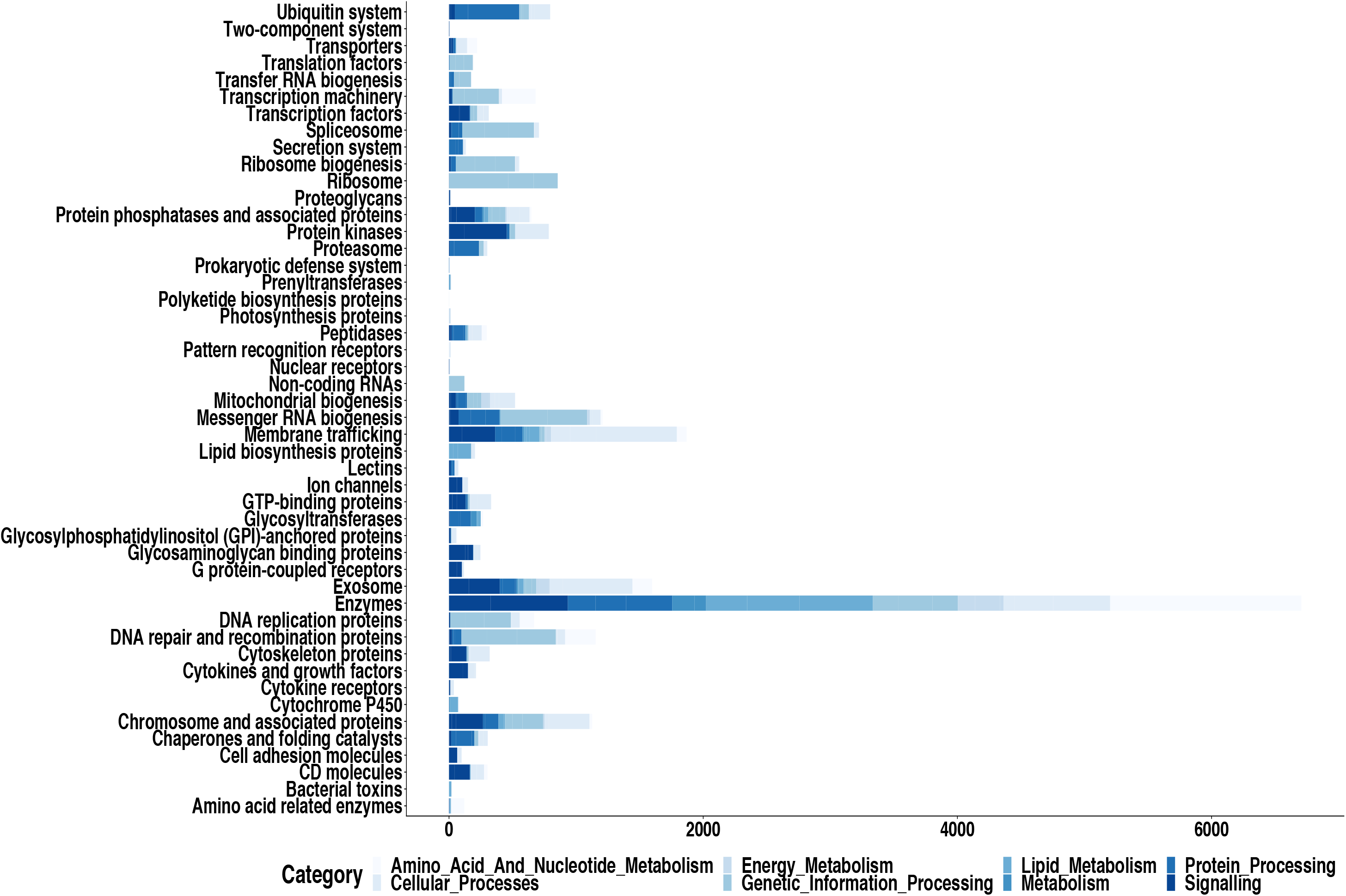
BRITE classifications of genes acquired during the transition between FECA to LECA, across the major categories of pathways that show gene gain during this period.

### Novel Functional Classes

Both in the cases of those classes that see the largest increases and in general overall, there is little demarcation of functional classes that belong to prokaryote genes or to genes developed at the early origins of eukaryotes (Figure 4-8). However we undertook to examine the genes that display a novel classification. During the transition to becoming LECA FECA expanded the repertoire of Prenyltransferases in the Lipid Metabolism pathways examined (Figure 4). Isoprenoid Biosynthesis isn’t unique to eukaryotes and Prenyl quinines are employed as electron carriers required for mitochondrial metabolism ^52^. Protein prenylation is an important post-translational modification that plays an important role in the membrane association of signal transduction regulatory elements and isoprenoids, in particular Dolichol - are involved in the production of glycoproteins ^52^. While glycosylation is found in all domains of life, it is much more restricted in prokaryotes ^53^, and this expansion of the process appears to have been important at the beginning of the eukaryotic lineage.

**Figure 4.**
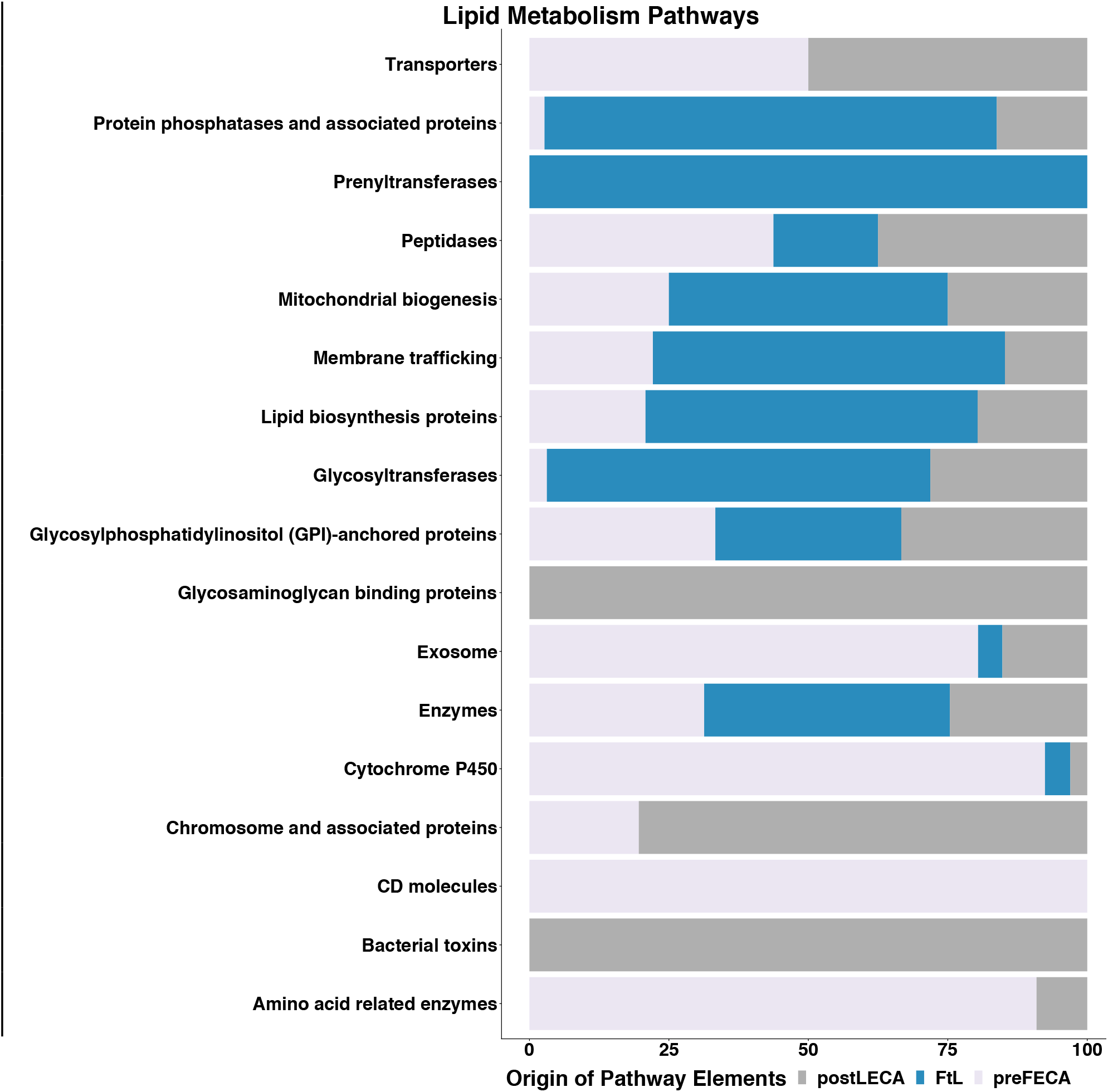
Proportion of BRITE functional classifications for genes, which originate pre-FECA, during the FECA to LECA transition, or post-LECA, from lipid metabolism pathways that saw gene gain during the FECA to LECA transition. FtL = FECA-to-LECA transition.

**Figure 5.**
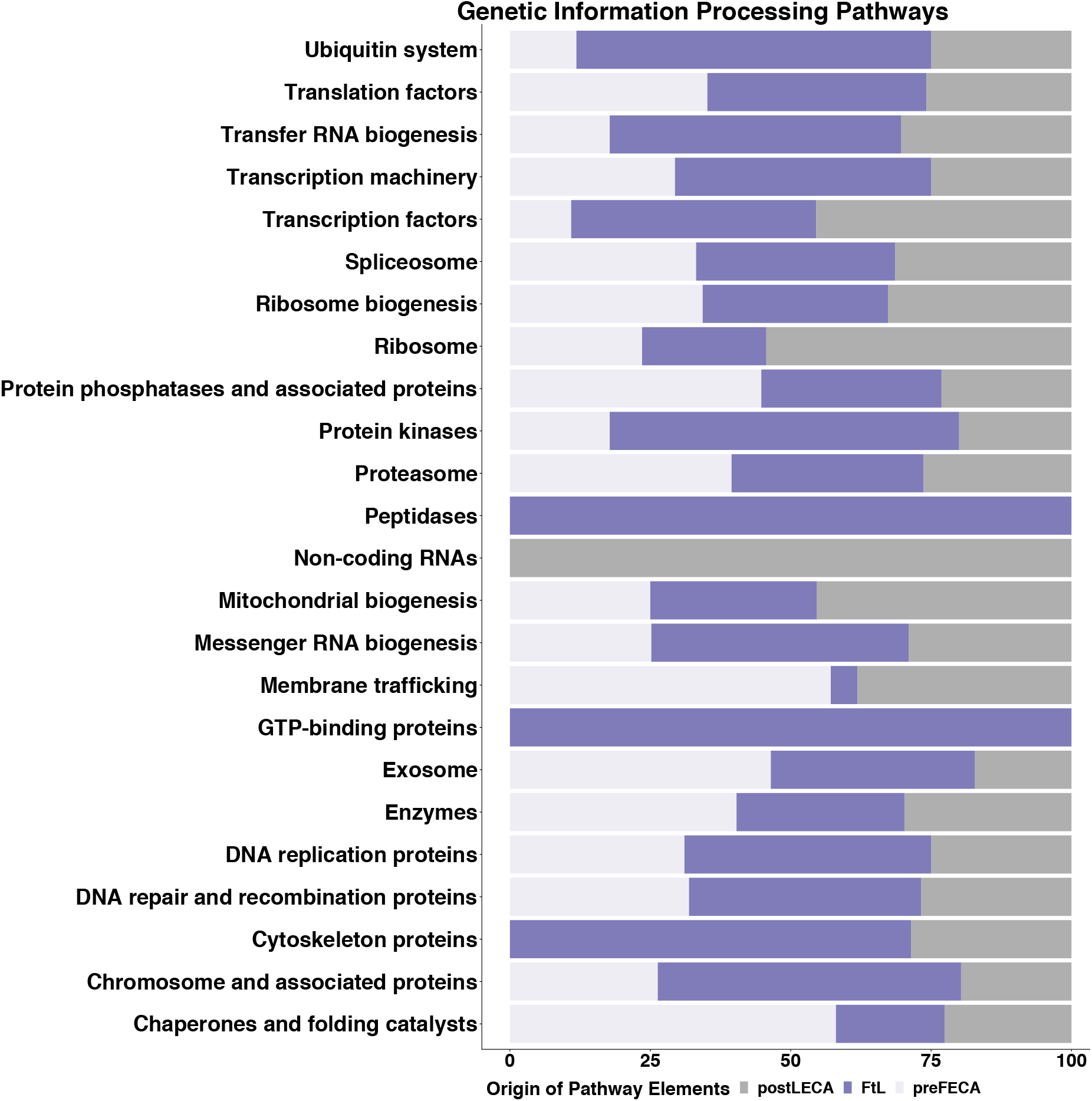
Proportion of BRITE functional classifications for genes, which originate pre-FECA, during the FECA to LECA transition, or post-LECA, from Genetic Information Processing pathways that saw gene gain during the FECA to LECA transition. FtL = FECA-to-LECA transition.

**Figure 6.**
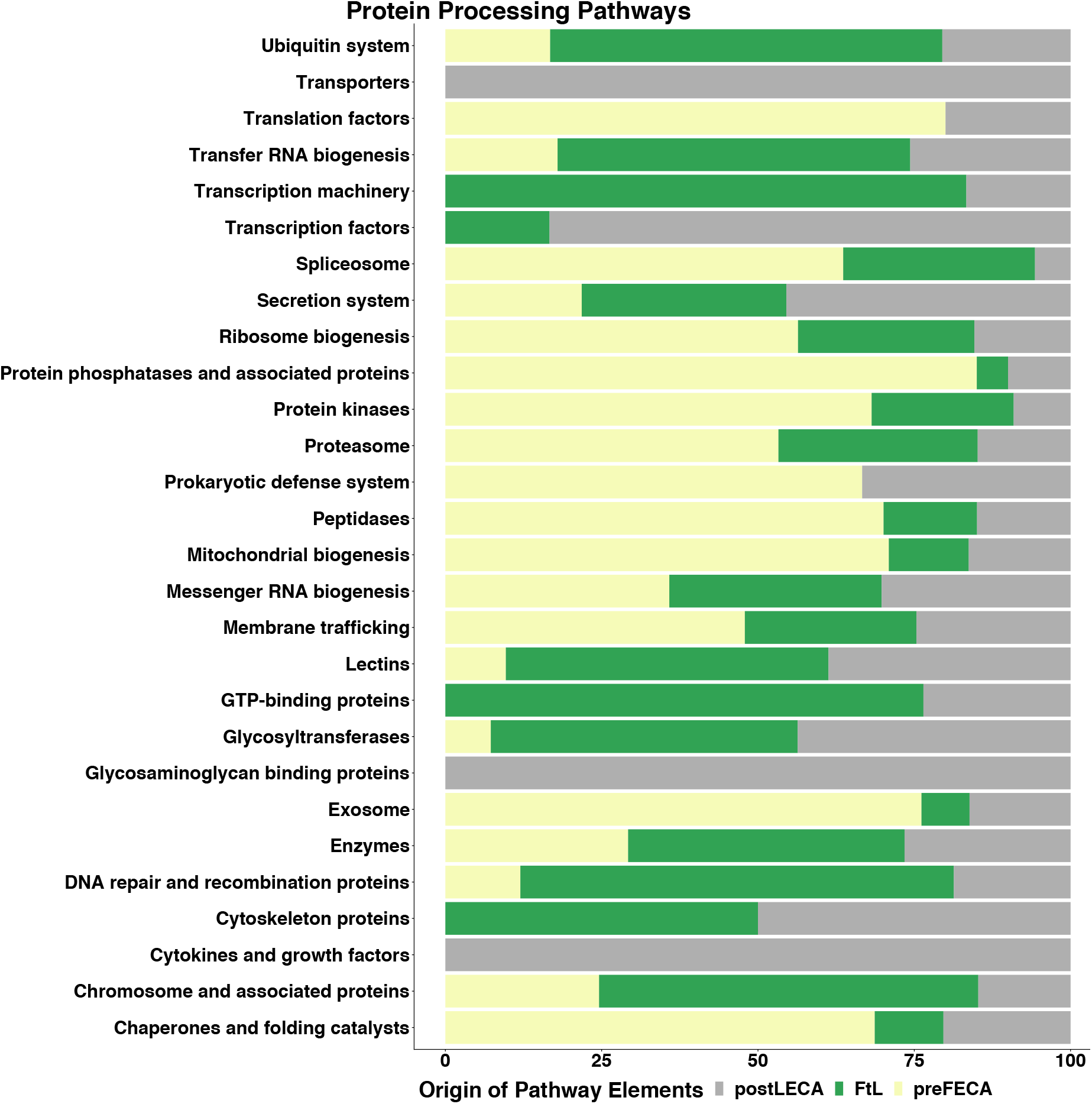
Proportion of BRITE functional classifications for genes, which originate pre-FECA, during the FECA to LECA transition, or post-LECA, from protein processing pathways that saw gene gain during the FECA to LECA transition. FtL = FECA-to-LECA transition.

**Figure 7.**
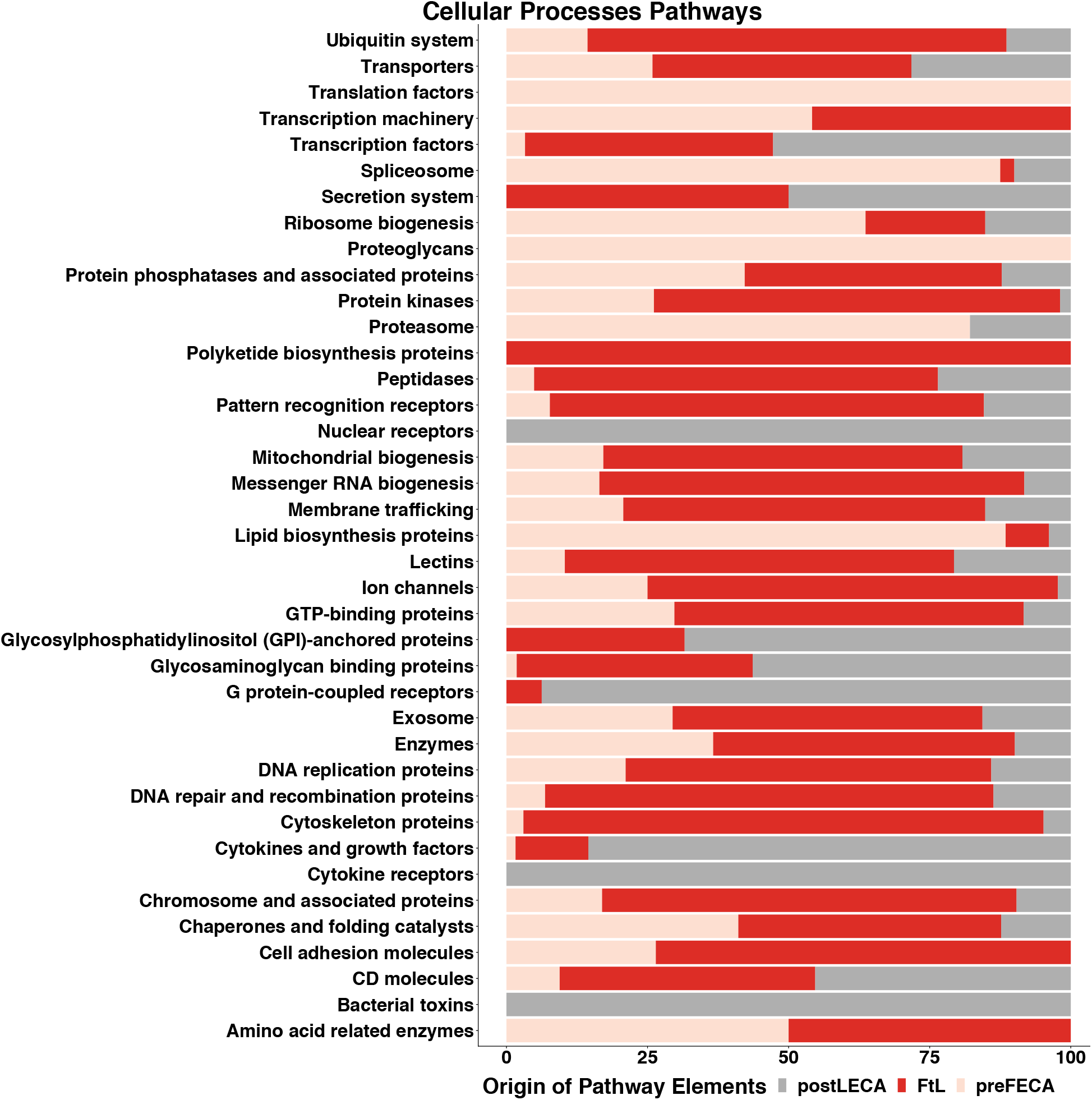
Proportion of BRITE functional classifications for genes, which originate pre-FECA, during the FECA to LECA transition, or post-LECA, from cellular processes pathways that saw gene gain during the FECA to LECA transition. FtL = FECA-to-LECA transition.

**Figure 8.**
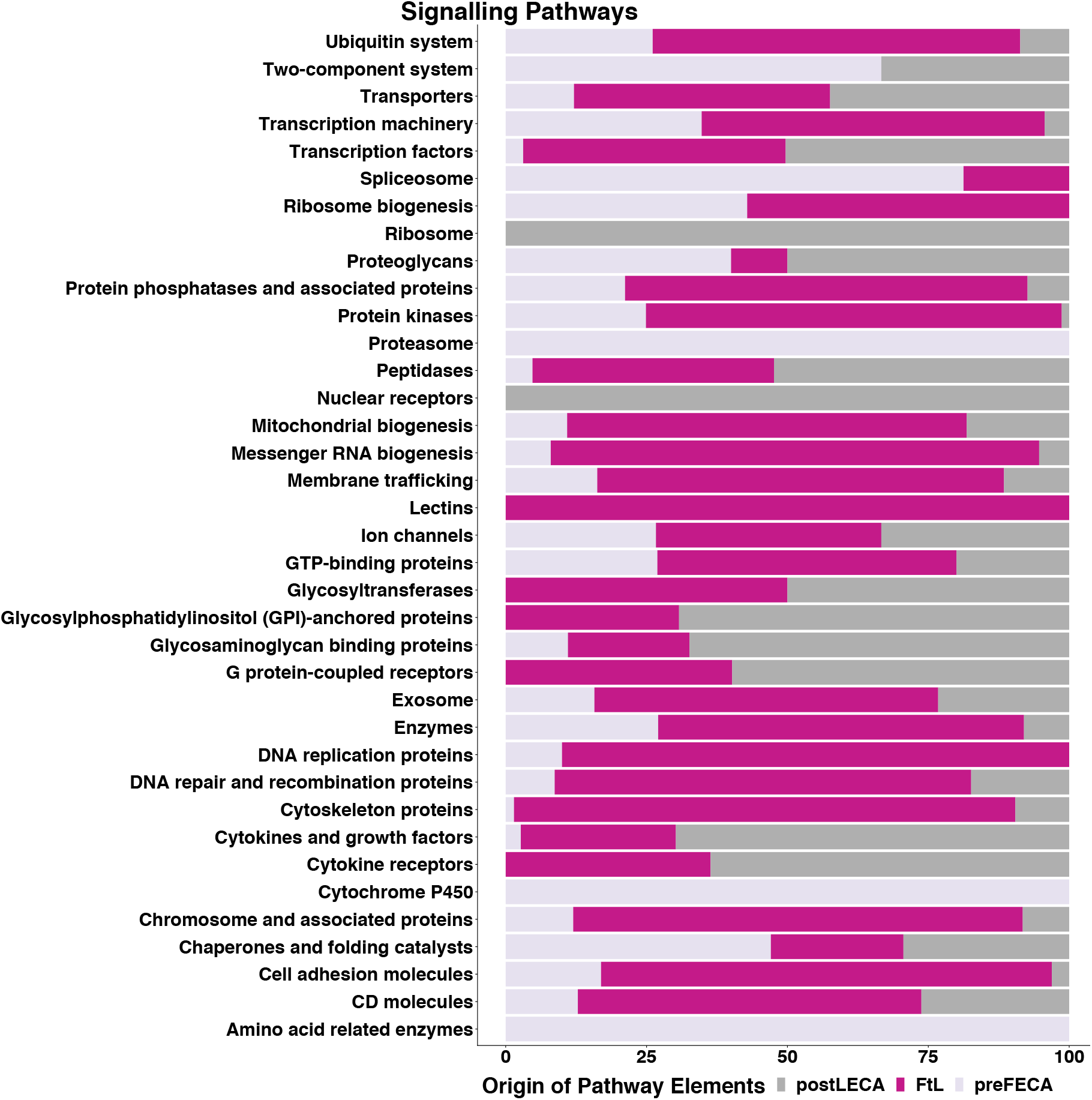
Proportion of BRITE functional classifications for genes, which originate pre-FECA, during the FECA to LECA transition, or post-LECA, from signalling pathways that saw gene gain during the FECA to LECA transition. FtL = FECA-to-LECA transition.

Genetic Information Processing pathways see the addition of GTP-binding proteins, cytoskeleton proteins, and peptidases during the FECA-to-LECA transition (Figure 5). GTP-binding proteins are signal transduction elements, which control a wide variety of processes from metabolism to gene expression ^54^. While Peptidases are a generic class of enzyme that exists across all domains of life ^55^, new family members arose in the basal transcription factors, homologous recombination, and the RNA transport pathways (SI Table 2). Cytoskeleton proteins control nuclear morphology and chromatin organization so it’s unsurprising that these types of elements were developed during this period ^51^ Specifically the element in question is the SUMO family protein SMT3 (SI Table 2), one of the main functions of sumoylation is nuclear-cytosolic transport as well as playing a role in DNA repair and recombination ^56,57^. Nuclear transport and transmitting signals to the nucleus posed new challenges for biological life and appears to have required new functional classes of genes.

Protein Processing pathways also see an addition of GTP-binding proteins and cytoskeleton proteins, as well as genes introduced to the transcription machinery and their repertoire of transcription factors (Figure 6). G-proteins have been discussed above in their myriad roles relating to signal transduction, and cytoskeleton proteins in this context interact with the organelles responsible for protein synthesis, modification, and trafficking ^58^. XRN2 adds an additional transcription machinery component relating to the RNA degradation, which promotes termination of transcription ^59^, and XBP1 adds a transcription factor relating to cellular stress response in the protein processing in endoplasmic reticulum pathways ^60^. Signal transduction and cytoskeleton elements, needed for coordinating the newly established structures of the cell are a reoccurring theme across these pathway categories. We also see novel elements in how transcription and translation operated in early eukaryotes.

G protein-coupled receptors, Secretion system, Glycosylphosphatidylinositol (GPI)-anchored proteins, and Polyketide biosynthesis proteins are introduced to the Cellular Processes pathways during this period (Figure 7). The Polyketide biosynthesis proteins class sees the addition of the Flavonoid biosynthesis gene CHS ^61^, which is present in the circadian rhythm pathway of plants as a downstream target (SI Table 2). Polyketides are common to plants, bacteria, and some ophistikonts, largely fungi ^61,62^. GPI-anchored proteins are minor plasma membrane components, involved in signal transduction and, perhaps critically in this context, play a role in clathrin-independent endocytosis ^63^. G protein-coupled receptors are part of the G-protein signal transduction pathways, another example of expansions in signal transduction machinery occurring during this evolutionary period. The secretion system elements belong to the Phagosome pathway and comprise all subunits of the ER membrane protein translocator Sec61 (SI Table 2). The bacterial gene SecY is widely reported as a homolog of Sec61 genes in the literature, these genes were cross-referenced against the gene trees in the KEGG database and not shown to possess bacterial homologs there, so we investigated this apparent false positive. Sequence analysis shows a little sequence similarity has been conserved between secY and the A1 (e-value of 0.011) and A2 subunit (e-value of 0.016), and almost none against the other subunits, with the sequence conservation being so poor it is unsurprising our method failed to identify this gene ^64^.

Signaling pathways see the largest number of classes added with G protein-coupled receptors, Cytokine receptors, Glycosylphosphatidylinositol (GPI)-anchored proteins, lectins, and glycosyltransferases (Figure 8). Given many of the other genes with novel classifications in the other categories relate to signal transduction this is perhaps unsurprising. Many of the classes are similar to the other pathway categories with the signal transduction G proteins and GPI-anchored proteins; additionally the cytokine receptors are a diverse class of signal transduction molecules ^65^. Glycosyltransferases and lectins, are involved in the production or recognition of glycoproteins ^66,67^, which is a reoccurring theme in this data. The development of signalling pathway elements, both in these pathways, but also present in the other categories discussed, reflects how critical systems of coordination become when cellular subcompartmentalisation and increasing cellular size makes passive diffusion ineffective and active transport across internal membranes is necessary.

## Discussion

Previous studies have shown that diverse representatives of the different eukaryote supergroups possess traits that can be mapped back to LECA ^2,13,14,41,68-70^. The results of all these reconstructions, based for the most part on distributions of phenotypic data across eukaryotes, consistently paint a picture of a LECA that already possessed significant complexity and contained many of the signature functional systems and structures of the modern eukaryotic cell. The benefit of whole-genome analyses is that we have greater precision in understanding the functional potential of LECA as well as the functions that predate this time period so that we can better understand these traits. Our reconstructed LECA genome is composed of 4,462 gene clusters when restricted to monophyletic clusters and 5,938 gene clusters when we include gene families with a complex history. Future reconstructions will no doubt refine and expand these results to identify more hypothetical LECA genes that are not identifiable by the current sampling. It is, of course, also likely that there will be a subset of genes in LECA that have been lost in the last two billion years of evolution and will therefore remain impossible to reconstruct with this type of method regardless of sampling. Nonetheless, we can reconstruct a vivid picture of LECA and its lifestyle. Our results suggest that LECA was likely to have been a facultatively anaerobic heterotroph, generating energy by a number of mechanisms, though not capable of photosynthesis (Figure 1, Figures S1-S5). LECA was able to engulf its prey using phagocytosis; but it remains unclear whether the elements with prokaryotic origin were sufficient to imbue FECA with a rudimentary ancestral manner of the process, leaving us unable to infer support for either the phagotrophic or autotrophic models of eukaryotic origin ^71^. LECA had a functioning Golgi apparatus and active vesicle trafficking within the cell but was not capable of contemporary forms of apoptosis. The cells were quite complex and RNA processing was carried out by spliceosomes (Figure 1, Figures S1-S5). It is likely that LECA had an endoplasmic reticulum and engaged in protein processing in this organelle. LECA cells divided by a recognisable mitosis, and recombination and cell division were also occurring via meiosis (Figure 1, Figures S1-S5).

A substantial portion of the genes from the pathways we reconstructed arose during the FECA-to-LECA transitional phase of eukaryote evolution, 1,554 genes of the 5,938 genes in total. The majority of these came from pathways relating to genetic information processing, protein production and modification, membrane-bound organelles, signalling, and lipid biosynthesis. These all constitute areas of biological function that were greatly modified as nuclear material was moved into the nucleus, the processes of transcription and translation were isolated and localised to specific regions within the cell forcing the machinery for them to be moved into the correct organelle and the products of these processes transported correctly ^13,72^. As processes became specialised and sub-localised in their own organelles and vesicles, the categories of pathways affected are entirely in-line with those that needed to have become heavily modified ^14,44^. Accordingly the functional annotations of the genes involved show a large propensity to be involved in Membrane Trafficking, various aspects of Genetic Information Processing, and signal transduction, consistent with establishing a system of intracellular biology of sub-compartmentalised processes and the regulatory systems for controlling them. The inclusion of substantial additions to DNA repair and recombination proteins specifically is also interesting because the basis of meiosis has been argued to have involved the recruitment and modification of this class of proteins ^73^.

The genes, which display novel functional classes, show a huge overlap across the different categories, with G protein pathway components as well as glycoprotein biosynthesis and interacting factors both being seen in all but one of the largest categories; Lipid Metabolism pathways lack G protein pathway components and Genetic Information Processing pathways lack Glycoprotein factors. The other classes we see that are widespread across these pathways are cytoskeleton proteins involved in coordinating both the nucleus and endomembrane system and a plethora of receptors, both nuclear and cell surface.

The functional classes of genes that were created during this transition from FECA to LECA primarily appear to have prokaryote origins, which lends additional credence to an increasing trend in the literature. There’s been a plethora of research in recent years on prokaryote encroachment into many aspects of biology that have been traditionally considered eukaryote-specific ^6-8,39,74-76^, it is perhaps worth examining the basis for the competing theories of eukaryogenesis. The apparent conflict between mito-late and mito-early theories hinges on our understanding of the complexity of prokaryotes. The distinction between the theories become largely semantic; one man’s “proto-eukaryote” is simply another’s Archaeal host, if we accept that archaea are increasingly eukaryote-like ^6-8,39,74-76^.

Overall our results show an interconnected collection of signalling elements, trafficking and transport complexes, nuclear and cell surface receptors, as well as to changes to genetic information and protein processing, and structural elements relating to the newly forming organelles (Figures 4-8, SI Table 2). These cellular components appear to be a consequence of having to cope with the challenges of establishing subcompartments that need to be navigated in a coordinated manner and functioning at a different scale to bacteria; signalling and membrane regulation and trafficking become increasingly important as diffusion becomes less feasible as a mechanism to transport necessities around one’s cell ^77^. These methods can’t disentangle whether the establishment of organelles or increasing cell size were downstream of the other or occurred in parallel. However it seems likely that a driving force for this process would be the increased levels of protein production allowed by the increased bioenergetic potential eukaryotes gained from endosymbiosis, which alongside the increasing availability of oxygen in their environment ^78^, led to levels of protein production at a level orders of magnitude higher than life was ever capable of before ^79,80^. The idea that mitochondria were necessary to fuel the size increase and subcompartmentalisation, which led to the need for the increases in membrane trafficking and signal transduction related gene functions has enormous appeal due to the neatness of the concept; endosymbiosis and the resultant mitochondria as the defining feature of eukaryotic life and driving force of all other observable differences.

## Methods and materials

### Genome choice

A dataset consisting of 32 taxonomically diverse eukaryote genomes, and 105 prokaryotes, comprising 36 Archaea and 69 Bacteria (total 855,880 genes) was acquired from publicly available genome databases, either Ensembl, Genomes version 33, or PATRIC (SI Table 1) ^81,82^.

### Gene Cluster Creation

Eukaryotes were selected to provide at least two representatives from all the major phylogenetic supergroups ^83^, including the SAR supergroup, opisthokonts, amoebozoa, archeaplastids, excavates, and orphan species. An all-versus-all BLAST search with an e-value cut-off of 10^−5^ was performed to identify homologous protein sequences. The BLAST output was clustered using the Markov Chain cluster algorithm (MCL) ^84^ with an inflation value of two after a series of parameter tests. The clusters were filtered according to their level of conservation across the six major taxa. Gene clusters conserved across at least four taxa were considered to be of a sufficient level of conservation to be strong LECA candidate genes (SI Data 3).

### Phylogenetic analysis

Clusters were aligned using MAAFT ^85^ and alignments were trimmed with TRIMAL ^86^ using the heuristic algorithm automated1. In the case of TRIMAL producing uninformative alignments the trimming step was skipped and the full sequence alignment was used. Phylogenetic hypotheses were reconstructed using the IQtree software ^87^, with parameters set to combine ModelFinder, tree search, SH-aLRT test and ultrafast bootstrap with 1000 replicates. Monophyly amongst the eukaryote sequences in these phylogenetic trees was assessed using the python package ETE3 ^88^, in order to be able to control for HGT. We then attempted to functionally characterise two datasets, a purely monophyletic dataset, and all sufficiently conserved gene clusters regardless of them demonstrating monophyly or a more complex evolutionary history.

### Functional annotation of Clusters

We used KEGG pathways from modern organisms that relate to metabolism, genetic information processing, environmental information processing, and cellular processes in order to ask whether a comparable pathway was likely to have been present in LECA (SI Table 2). We used exemplar protein sequences from the KEGG model organisms *Homo sapiens, Saccharomyces cerevisiae, Arabidopsis thaliana, Dictyostelium discoideum, Methanobrevibacter smithii*, and *Escherichia coli* where available and compared sequences from these organisms to our gene clusters in order to determine presence or absence. Homology to the highly conserved gene clusters was determined using BLAST with an e-value cutoff of 10^−6^.

### Dataset Subsetting and Analysis

For the analysis of the FECA to LECA evolutionary transition the KEGG pathway mapped data was subsetted to include only clusters that did not possess prokaryote homologs ^89^. Each KEGG pathway element that matched to a eukaryote specific gene cluster was cross-referenced against the gene tree in the KEGG database to independently verify the evolutionary origin of the gene, producing a list of elements found in LECA with no known prokaryote homology. This list of genes created during the FECA to LECA transition were then categorised and functionally annotated extracting functional information from the KEGG database BRITE classifications.

The categories used in this analysis of genes created in the FECA to LECA transition were modified slightly from the KEGG classifications as follows: The Energy Metabolism category is altered from the KEGG database schema to include the 1.1 Carbohydrate metabolism pathway for Glycolysis (0010). Lipid Metabolism and Cellular Processes categories are identical in composition to the KEGG database categories. The Metabolism category collects 1.1 Carbohydrate metabolism pathways, excluding Glycolysis and incorporating 1.9 Metabolism of terpenoids and polyketides. The Protein Processing category combines the Genetic Information Processing sub-category 2.3 Folding, sorting and degradation with the 1.7 Glycan biosynthesis and metabolism pathways. Genetic Information Processing is composed of the remaining KEGG categories – 2.1 Transcription, 2.2 Translation and 2.4 Replication and repair. The Signalling category is composed of pathways grouped under the Environmental Information Processing category in the KEGG database. Amino Acid and Nucleotide Metabolism combines the three categories of 1.4 Nucleotide metabolism, 1.5 Amino acid metabolism and 1.6 Metabolism of other amino acids.

### Author Affiliations

FJW has received funding from the European Union’s Horizon 2020 research and innovation programme under the Marie Skłodowska-Curie grant agreement No. 793818. JOM was awarded funding from the Templeton Foundation to support the work of DWN and MM. JOM was awarded funding from the BBSRC to support the work of MJR and FJW.

---

**Supplementary Data 1 – LECA Reconstruction Neo4j Database.**

https://github.com/JMcInerneyLab/LECA_Project

**Supplementary Data 2 – Raw data of the clusters that match to KEGG pathway elements.**

https://github.com/JMcInerneyLab/LECA_Project

**Supplementary Data 3 – LECA Gene Cluster lists.**

https://github.com/JMcInerneyLab/LECA_Project

**Supplementary Information Table 1 – List of Genomes used.**

**Supplementary Information Table 2 – List of Gene Gains during FECA to LECA transition.**

**Supplementary Figure S1 – Annotated KEGG Pathways for Energy metabolism related pathways.**

**Supplementary Figure S2 – Annotated KEGG Pathways for Mitosis and Cell cycle related pathways.**

**Supplementary Figure S3 – Annotated KEGG Pathways for Meiosis related pathways.**

**Supplementary Figure S4 – Annotated KEGG Pathways for Membrane related pathways.**

**Supplementary Figure S5 – Annotated KEGG Pathways for Gene Expression, Protein Production, and MAPK signalling related pathways.**

**Supplementary Figure S6 – Heatmaps of pathway completeness for amoeba, plant, yeast, bacterial, and archaeal pathways across the consitutent genomes of the reconstructed LECA genome.**

## References

1 Mayr, E. in Proc Natl Acad Sci USA Vol. 95 9720–9723 (National Academy of Sciences, 1998).

2 Ovádi, J. & Saks, V. in Mol Cell Biochem Vol. 256 5–12 (2004).

3 Martin, W. in Philosophical Transactions of the Royal Society B: Biological Sciences Vol. 365 847–855 (The Royal Society, 2010).

4 Jékely, G. in Eukaryotic Membranes and Cytoskeleton Vol. 607 38–51 (Springer, New York, NY, 2007).

5 Cox, C. J., Foster, P. G., Hirt, R. P., Harris, S. R. & Embley, T. M. in Proc Natl Acad Sci USA Vol. 105 20356–20361 (2008).

6 Spang, A. et al. in Nature Vol. 521 nature 14447–14179 (Nature Publishing Group, 2015).

7 Zaremba-Niedzwiedzka, K. et al. in Nature Vol. 541 353–358 (2017).

8 Akıl, C. & Robinson, R. C. in Nature Vol. 562 439–443 (Nature Publishing Group, 2018).

9 Hudder, A., Nathanson, L. & Deutscher, M. P. Organization of Mammalian Cytoplasm. Molecular and Cellular Biology 23, 9318–9326, doi:10.1128/mcb.23.24.9318-9326.2003 (2003).

10 Guigas, G., Kalla, C. & Weiss, M. in FEBS Letters Vol. 581 5094–5098 (2007).

11 Dacks, J. B., Peden, A. A. & Field, M. C. in The International Journal of Biochemistry & Cell Biology Vol. 41 330–340 (2009).

12 Field, M. C. & Dacks, J. B. in Current Opinion in Cell Biology Vol. 21 4–13 (2009).

13 Martin, W. & Koonin, E. V. in Nature Vol. 440 41–45 (2006).

14 Gabaldon, T. & Pittis, A. A. Origin and evolution of metabolic sub-cellular compartmentalization in eukaryotes. Biochimie 119, 262–268, doi:10.1016/j.biochi.2015.03.021 (2015).

15 van der Giezen, M. in Journal of Eukaryotic Microbiology Vol. 56 221–231 (Wiley/Blackwell (10.1111), 2009).

16 Hjort, K., Goldberg, A. V., Tsaousis, A. D., Hirt, R. P. & Embley, T. M. in Philosophical Transactions of the Royal Society B: Biological Sciences Vol. 365 713–727 (2010).

17 van der Giezen, M. & Tovar, J. in EMBO Rep. Vol. 6 525–530 (John Wiley & Sons, Ltd, 2005).

18 Tovar, J., Fischer, A. & Clark, C. G. in Molecular Microbiology Vol. 32 1013–1021 (Blackwell Science Ltd, 1999).

19 Ciechanover, A., Orian, A. & Schwartz, A. L. in BioEssays Vol. 22 442–451 (2000).

20 Dikic, I., Wakatsuki, S. & Walters, K. J. in Nat Rev Mol Cell Biol Vol. 10 659–671 (2009).

21 Hershko, A. & Ciechanover, A. in Annu. Rev. Biochem. Vol. 67 425–479 (1998).

22 Hunter, T. in Molecular Cell Vol. 28 730–738 (2007).

23 Johnson, S. A. & Hunter, T. in Nat Meth Vol. 2 17–25 (2005).

24 Pawson, T. in Cell Vol. 116 191–203 (Cell Press, 2004).

25 Pawson, T. & Kofler, M. in Current Opinion in Cell Biology Vol. 21 147–153 (Elsevier Current Trends, 2009).

26 Till, J. H. & Hubbard, S. R. in BioEssays Vol. 69 373–398 (Annual Reviews 4139 El Camino Way, P.O. Box 10139, Palo Alto, CA 94303-0139, USA, 2003).

27 Amaral, P. P., Dinger, M. E., Mercer, T. R. & Mattick, J. S. in Science Vol. 319 1787–1789 (2008).

28 Chapman, E. J. & Carrington, J. C. in Nature Publishing Group Vol. 8 884–896 (2007).

29 D’Alessio, J. A., Wright, K. J. & Tjian, R. in Molecular Cell Vol. 36 924–931 (NIH Public Access, 2009).

30 Jacquier, A. in Nat Rev Genet Vol. 10 833–844 (2009).

31 Liu, Q. & Paroo, Z. in Annu. Rev. Biochem. Vol. 79 295–319 (2010).

32 Wilson, M. D. & Odom, D. T. in Current Opinion in Genetics & Development Vol. 19 579–585 (2009).

33 Sagan, L. in Journal of Theoretical Biology Vol. 14 225–274 (1967).

34 Poole, A. M. & Gribaldo, S. in Cold Spring Harbor Perspectives in Biology Vol. 6 a015990–a015990 (Cold Spring Harbor Lab, 2014).

35 Koonin, E. V. in Genome Biol. Vol. 11 1–12 (2010).

36 Martin, W. F., Garg, S. & Zimorski, V. in Philosophical Transactions of the Royal Society B: Biological Sciences Vol. 370 20140330 (The Royal Society, 2015).

37 Cavalier-Smith, T. in Nature Vol. 326 332–333 (1987).

38 Embley, T. M. & Martin, W. in Nature Vol. 440 623–630 (Nature Publishing Group, 2006).

39 Grant, C. R., Wan, J. & Komeili, A. in Annu. Rev. Cell Dev. Biol. Vol. 34 217–238 (2018).

40 Nickelsen, J. et al. in FEMS Microbiology Letters Vol. 315 1–5 (John Wiley & Sons, Ltd, 2011).

41 Makarova, K. S. in Nucleic Acids Research Vol. 33 4626–4638 (2005).

42 McInerney, J., pisani, D. & O’Connell, M. J. in Philosophical Transactions of the Royal Society B: Biological Sciences Vol. 370 20140323–20140625 (The Royal Society, 2015).

43 Roger, A. J., Muñoz-Gómez, S. A. & Kamikawa, R. in Current Biology Vol. 27 R1177–R1192 (Elsevier Ltd, 2017).

44 Dacks, J. B. & Field, M. C. in Current Opinion in Cell Biology Vol. 53 70–76 (Elsevier Current Trends, 2018).

45 Hunter, T. in Cell Vol. 100 113–127 (Elsevier, 2000).

46 Kelly, S. M. & Corbett, A. H. in Traffic Vol. 10 1199–1208 (NIH Public Access, 2009).

47 Mitchell, P., Petfalski, E., Shevchenko, A., Mann, M. & Tollervey, D. in Cell Vol. 91 457–466 (Cell Press, 1997).

48 Petit, C. & Sancar, A. in Biochimie Vol. 81 15–25 (Elsevier, 1999).

49 Lin, Z., Kong, H., Nei, M. & Ma, H. in Proc Natl Acad Sci USA Vol. 103 10328–10333 (National Academy of Sciences, 2006).

50 Doma, M. K. & Parker, R. in Cell Vol. 131 660–668 (Elsevier, 2007).

51 Ramdas, N. M. & Shivashankar, G. V. in J. Mol. Biol. Vol. 427 695–706 (Academic Press, 2015).

52 Takahashi, S. & Koyama, T. in The Chemical Record Vol. 6 194–205 (John Wiley & Sons, Ltd, 2006).

53 Dell, A., Galadari, A., Sastre, F. & Hitchen, P. in International Journal of Microbiology Vol. 2010 1–14 (Hindawi Limited, 2010).

54 Neves, S. R. in Science Vol. 296 1636–1639 (American Association for the Advancement of Science, 2002).

55 Rawlings, N. D. & Barrett, A. J. in Biochemical Journal Vol. 290 (Pt 1) 205–218 (1993).

56 Jalal, D., Chalissery, J. & Hassan, A. H. in Nucleic Acids Research Vol. 14 gkw1369–2261 (2017).

57 Matunis, M. J., Coutavas, E. & Blobel, G. in J Cell Biol Vol. 135 1457–1470 (1996).

58 Nielsen, P., Goelz, S. & Trachsel, H. in Cell Biology International Reports Vol. 7 245–254 (No longer published by Elsevier, 1983).

59 West, S., Gromak, N. & Proudfoot, N. J. in Nature Vol. 432 522–525 (Nature Publishing Group, 2004).

60 Yoshida, H., Nadanaka, S., Sato, R. & Mori, K. in Cell Struct. Funct. Vol. 31 117–125 (Japan Society for Cell Biology, 2006).

61 Tohge, T., Yonekura-Sakakibara, K., Niida, R., Watanabe-Takahashi, A. & Saito, K. in Pure and Applied Chemistry Vol. 79 811–823 (2007).

62 Castoe, T. A., Stephens, T., Noonan, B. P. & Calestani, C. in Gene Vol. 392 47–58 (Elsevier, 2007).

63 Imhof, I. et al. in J. Biol. Chem. Vol. 279 19614–19627 (American Society for Biochemistry and Molecular Biology, 2004).

64 Gold, V. A. M., Duong, F. & Collinson, I. in Molecular Membrane Biology Vol. 24 387–394 (Taylor & Francis, 2009).

65 Kishimoto, T., Taga, T. & Akira, S. in Cell Vol. 76 253–262 (Cell Press, 1994).

66 Breton, C. & Imberty, A. in Curr Opin Struct Biol Vol. 9 563–571 (Elsevier Current Trends, 1999).

67 Goldstein, I. J. & Hayes, C. E. in Advances in Carbohydrate Chemistry and Biochemistry Vol. 35 127–340 (Academic Press, 1978).

68 Goodenough, U. & Heitman, J. in Cold Spring Harbor Perspectives in Biology Vol. 6 a016154–a016154 (2014).

69 Loidl, J. in Annu. Rev. Genet. Vol. 50 293–316 (2016).

70 Koonin, E. V. et al. in Genome Biol. Vol. 5 R7 (2004).

71 O’Malley, M. A. in Studies in History and Philosophy of Science Part C: Studies in History and Philosophy of Biological and Biomedical Sciences Vol. 41 212–224 (Pergamon, 2010).

72 Martin, W. Archaebacteria (Archaea) and the origin of the eukaryotic nucleus. Curr Opin Microbiol 8, 630–637, doi:10.1016/j.mib.2005.10.004 (2005).

73 Marcon, E. & Moens, P. B. in BioEssays Vol. 27 795–808 (John Wiley & Sons, Ltd, 2005).

74 Trépout, S. & Wehenkel, A. M. in Trends in Microbiology 1–2 (Elsevier Ltd, 2017).

75 Errington, J. in Nature Cell Biology 2001 3:8 Vol. 5 175–178 (Nature Publishing Group, 2003).

76 Thanbichler, M., Wang, S. C. & Shapiro, L. in Journal of Cellular Biochemistry Vol. 96 506–521 (John Wiley & Sons, Ltd, 2005).

77 de Duve, C. in Sci Am Vol. 274 50–57 (1996).

78 Lenton, T. M., Boyle, R. A., Poulton, S. W., Shields-Zhou, G. A. & Butterfield, N. J. in Nature Geosci Vol. 7 257–265 (2014).

79 Lane, N. & Martin, W. in Nature Vol. 467 929–934 (Nature Publishing Group, 2010).

80 Lane, N. & Martin, W. F. in Proc Natl Acad Sci USA Vol. 112 E4823–E4823 (National Academy of Sciences, 2015).

81 Wattam, A. R. et al. in Nucleic Acids Research Vol. 45 D535–D542 (2017).

82 Zerbino, D. R. et al. in Nucleic Acids Research Vol. 46 D754–D761 (Oxford University Press, 2018).

83 Burki, F. The eukaryotic tree of life from a global phylogenomic perspective. Cold Spring Harb Perspect Biol 6, a016147, doi:10.1101/cshperspect.a016147 (2014).

84 Enright, A. J., Van Dongen, S. & Ouzounis, C. A. in Nucleic Acids Research Vol. 30 1575–1584 (Oxford University Press, 2002).

85 Katoh, K., Misawa, K., Kuma, K.-i. & Miyata, T. in Nucleic Acids Research Vol. 30 3059–3066 (Oxford University Press, 2002).

86 Capella-Gutiérrez, S., Silla-Martínez, J. M. & Gabaldón, T. in Bioinformatics Vol. 25 1972–1973 (Oxford University Press, 2009).

87 Nguyen, L.-T., Schmidt, H. A., von Haeseler, A. & Minh, B. Q. in Molecular Biology and Evolution Vol. 32 268–274 (Oxford University Press, 2015).

88 Huerta-Cepas, J., Serra, F. & Bork, P. in Molecular Biology and Evolution Vol. 33 1635–1638 (Oxford University Press, 2016).

89 Kanehisa, M., Furumichi, M., Tanabe, M., Sato, Y. & Morishima, K. in Nucleic Acids Research Vol. 45 D353–D361 (2017).

