## Supplemental Figure S1 for "Reconstructing and Analysing The Genome of The Last Eukaryote Common Ancestor to Better Understand the Transition from FECA to LECA"

A.

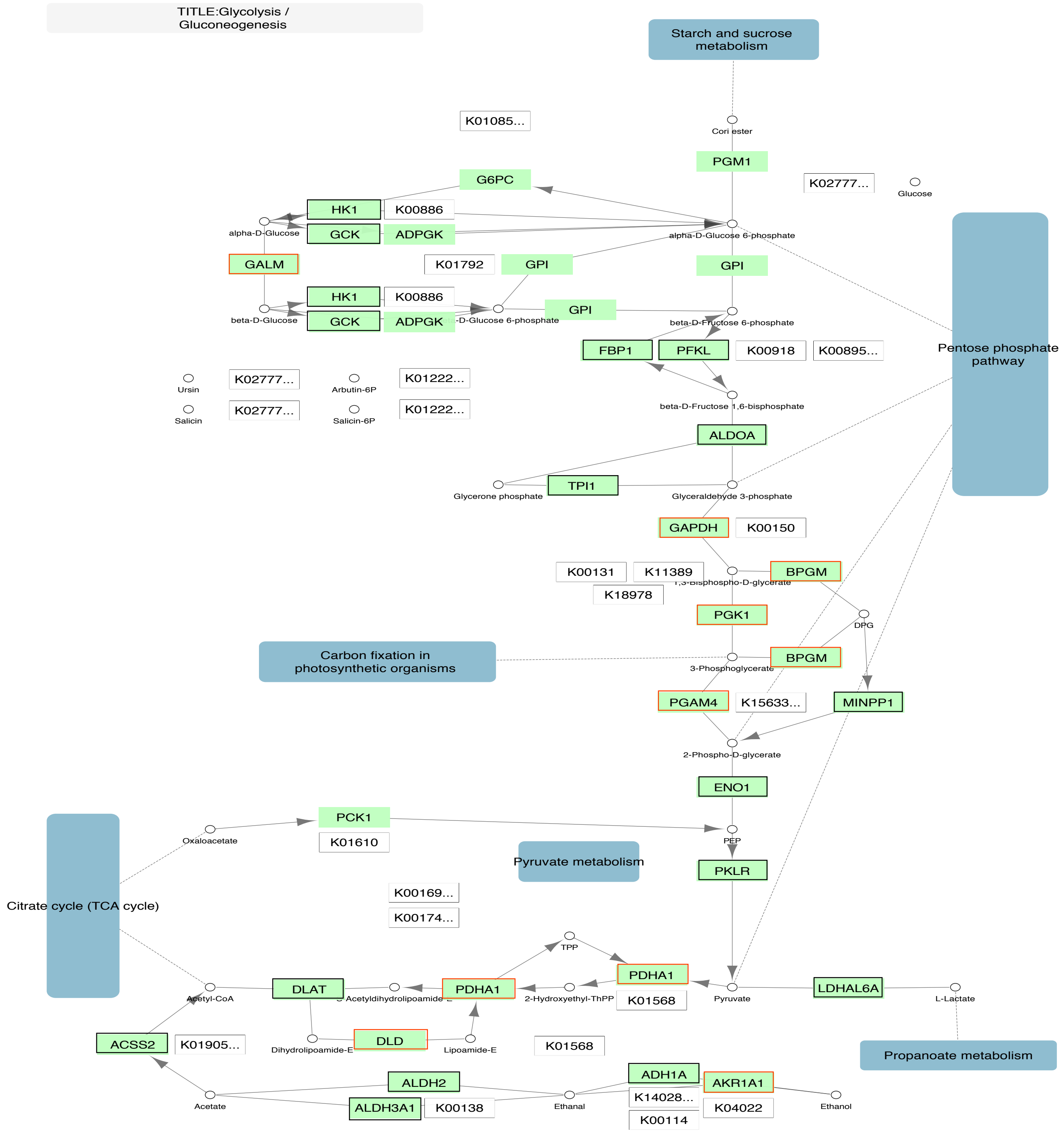
TITLE:Glycolysis / Gluconeogenesis
Starch and sucrose metabolism
K01085...
Cori ester
PGM1
K02777...
Glucose
G6PC
HK1
K00886
alpha-D-Glucose
GCK
ADPGK
alpha-D-Glucose 6-phosphate
GALM
K01792
GPI
GPI
Pentose phosphate pathway
HK1
K00886
GPI
beta-D-Glucose
GCK
ADPGK
beta-D-Fructose 6-phosphate
FBP1
PFKL
K00918
K00895...
Ursin
K02777...
Arbutin-6P
K01222...
Salicin
K02777...
Salicin-6P
K01222...
beta-D-Fructose 1,6-bisphosphate
ALDOA
Glycerone phosphate
TPI1
Glyceraldehyde 3-phosphate
GAPDH
K00150
K00131
K11389
BPGM
K18978
DPG
PGK1
Carbon fixation in photosynthetic organisms
BPGM
3-Phosphoglycerate
PGAM4
K15633...
MINPP1
2-Phospho-D-glycerate
ENO1
PCK1
Oxaloacetate
K01610
PEP
Pyruvate metabolism
PKLR
Citrate cycle (TCA cycle)
K00169...
K00174...
TPP
PDHA1
Acetyl-CoA
DLAT
PDHA1
2-Hydroxyethyl-ThPP
K01568
Pyruvate
LDHAL6A
L-Lactate
Dihydrolipoamide-E
DLD
Lipoamide-E
ACSS2
K01905...
K01568
Propanoate metabolism
ALDH2
ADH1A
AKR1A1
Acetate
ALDH3A1
K00138
Ethanal
K14028...
K04022
Ethanol
K00114

B.

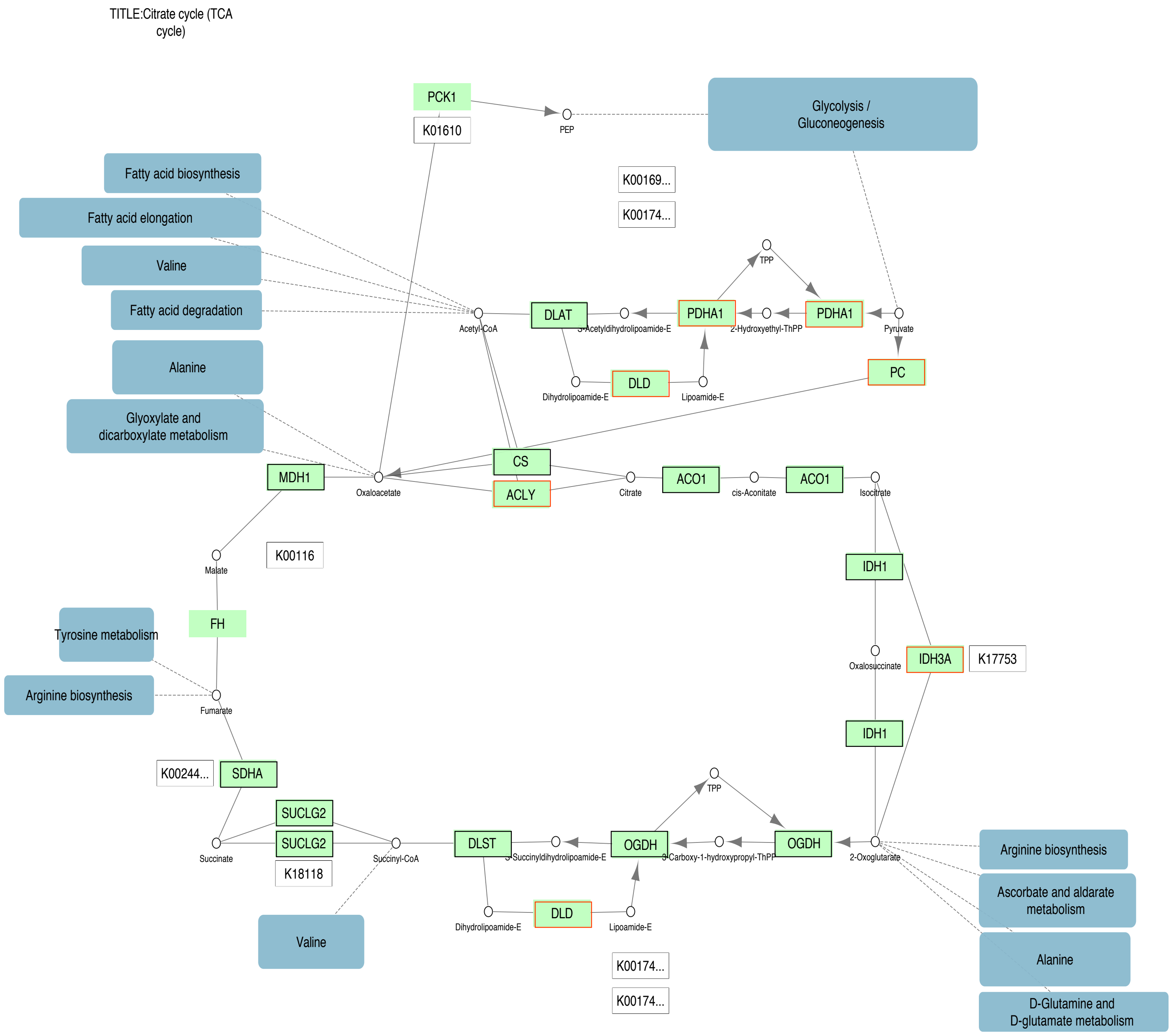

TITLE:Citrate cycle (TCA cycle)
PCK1
K01610
PEP
Glycolysis / Gluconeogenesis
Fatty acid biosynthesis
Fatty acid elongation
Valine
Fatty acid degradation
Alanine
Glyoxylate and dicarboxylate metabolism
K00169...
K00174...
TPP
Acetyl-CoA
DLAT
S-Acetyldihydrolipoamide-E
PDHA1
2-Hydroxyethyl-ThPP
PDHA1
Pyruvate
Dihydrolipoamide-E
DLD
Lipoamide-E
PC
MDH1
Oxaloacetate
CS
ACLY
Citrate
ACO1
cis-Aconitate
ACO1
Isocitrate
K00116
Malate
IDH1
FH
Tyrosine metabolism
Oxalosuccinate
IDH3A
K17753
Arginine biosynthesis
Fumarate
IDH1
K00244...
SDHA
TPP
SUCLG2
SUCLG2
Succinate
Succinyl-CoA
DLST
S-Succinyldihydrolipoamide-E
OGDH
3-Carboxy-1-hydroxypropyl-ThPP
OGDH
2-Oxoglutarate
Arginine biosynthesis
K18118
Ascorbate and aldarate metabolism
Dihydrolipoamide-E
DLD
Lipoamide-E
Valine
Alanine
K00174...
K00174...
D-Glutamine and D-glutamate metabolism

C.

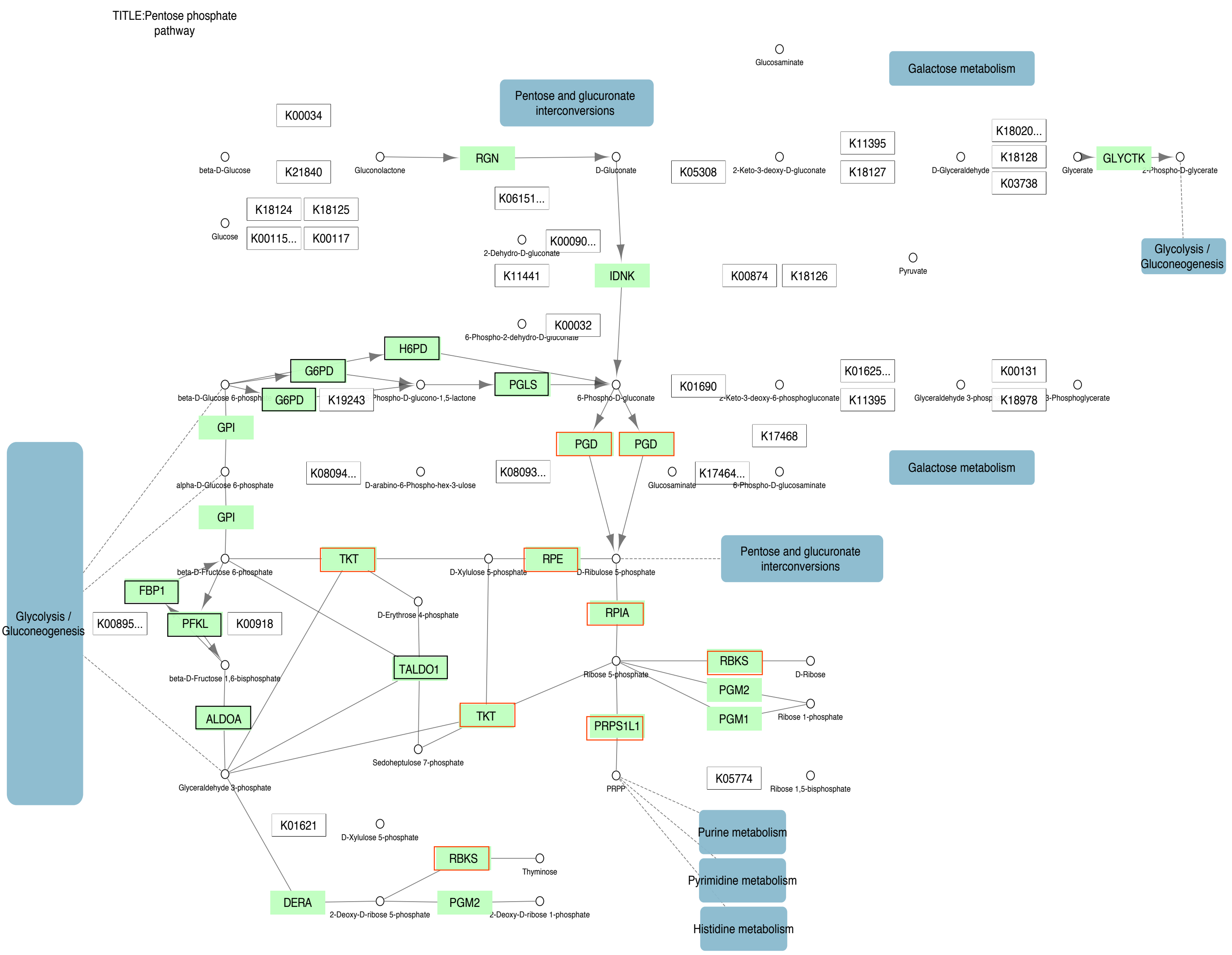

TITLE:Pentose phosphate pathway
Glucosaminate
Galactose metabolism
Pentose and glucuronate interconversions
K00034
beta-D-Glucose
K21840
Gluconolactone
RGN
D-Gluconate
K05308
2-Keto-3-deoxy-D-gluconate
K11395
K18127
K18020...
D-Glyceraldehyde
K18128
K03738
Glycerate
GLYCTK
2-Phospho-D-glycerate
K18124
K18125
Glucose
K00115...
K00117
K06151...
2-Dehydro-D-gluconate
K00090...
K11441
IDNK
K00874
K18126
Pyruvate
Glycolysis / Gluconeogenesis
6-Phospho-2-dehydro-D-gluconate
K00032
H6PD
G6PD
PGLS
beta-D-Glucose 6-phosph
G6PD
K19243
Phospho-D-glucono-1,5-lactone
6-Phospho-D-gluconate
K01690
2-Keto-3-deoxy-6-phosphogluconate
K01625...
K11395
Glyceraldehyde 3-phosp
K00131
K18978
3-Phosphoglycerate
GPI
PGD
PGD
K17468
alpha-D-Glucose 6-phosphate
K08094...
D-arabino-6-Phospho-hex-3-ulose
K08093...
Glucosaminate
K17464...
6-Phospho-D-glucosaminate
Galactose metabolism
GPI
beta-D-Fructose 6-phosphate
TKT
D-Xylulose 5-phosphate
RPE
D-Ribulose 5-phosphate
Pentose and glucuronate interconversions
FBP1
D-Erythrose 4-phosphate
RPIA
Glycolysis / Gluconeogenesis
K00895...
PFKL
K00918
beta-D-Fructose 1,6-bisphosphate
TALDO1
Ribose 5-phosphate
RBKS
D-Ribose
PGM2
PGM1
Ribose 1-phosphate
ALDOA
TKT
PRPS1L1
Sedoheptulose 7-phosphate
Glyceraldehyde 3-phosphate
PRPP
K05774
Ribose 1,5-bisphosphate
K01621
D-Xylulose 5-phosphate
Purine metabolism
RBKS
Thyminose
Pyrimidine metabolism
DERA
2-Deoxy-D-ribose 5-phosphate
PGM2
2-Deoxy-D-ribose 1-phosphate
Histidine metabolism

D.

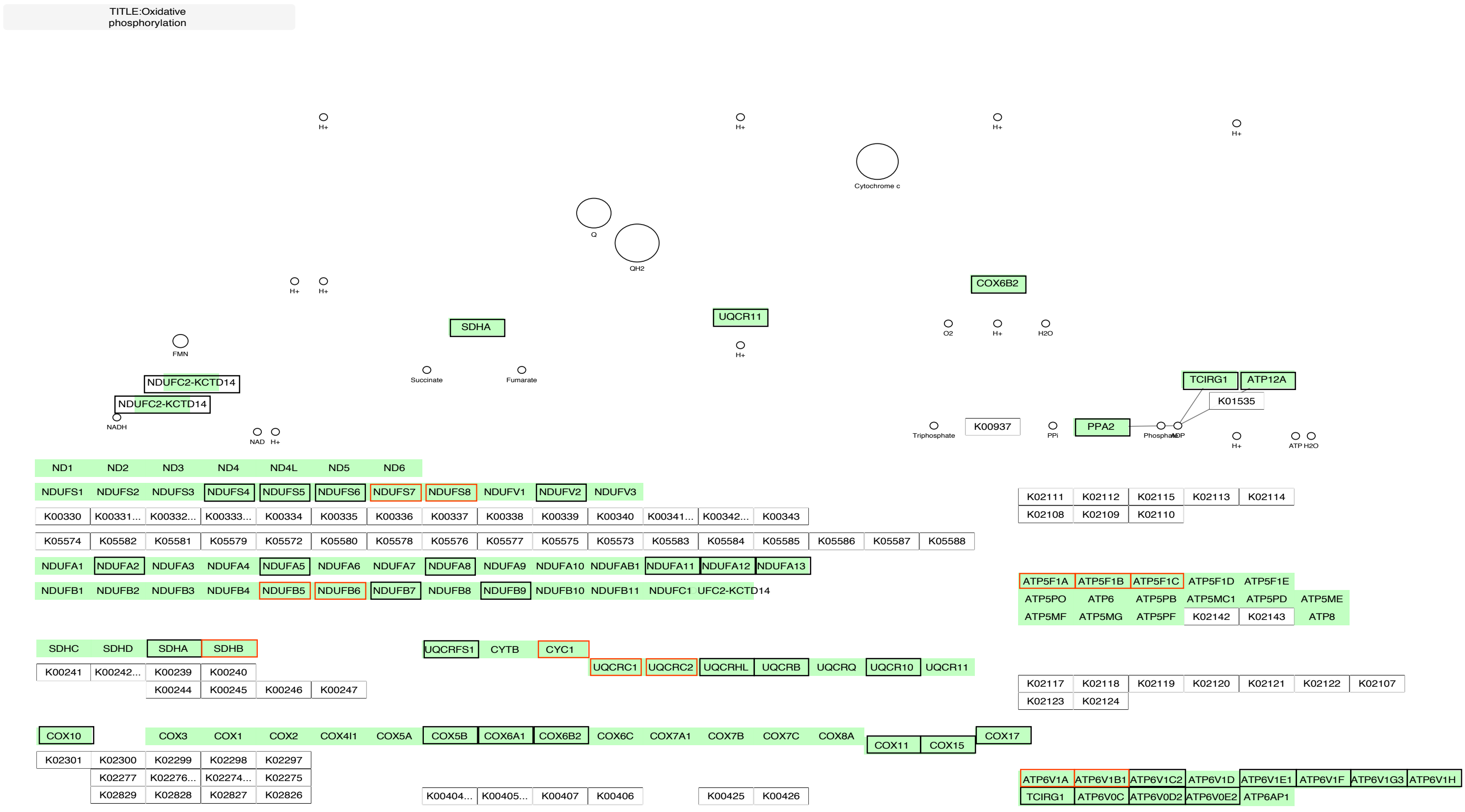
TITLE:Oxidative phosphorylation
H+
H+
H+
H+
Cytochrome c
Q
QH2
H+
H+
COX6B2
SDHA
UQCR11
O2
H+
H2O
FMN
H+
Succinate
Fumarate
NDUFC2-KCTD14
NDUFC2-KCTD14
NADH
NAD
H+
TCIRG1
ATP12A
K01535
Triphosphate
K00937
PPi
PPA2
Phosphate
ADP
H+
ATP
H2O
ND1
ND2
ND3
ND4
ND4L
ND5
ND6
NDUFS1
NDUFS2
NDUFS3
NDUFS4
NDUFS5
NDUFS6
NDUFS7
NDUFS8
NDUFV1
NDUFV2
NDUFV3
K00330
K00331...
K00332...
K00333...
K00334
K00335
K00336
K00337
K00338
K00339
K00340
K00341...
K00342...
K00343
K05574
K05582
K05581
K05579
K05572
K05580
K05578
K05576
K05577
K05575
K05573
K05583
K05584
K05585
K05586
K05587
K05588
NDUFA1
NDUFA2
NDUFA3
NDUFA4
NDUFA5
NDUFA6
NDUFA7
NDUFA8
NDUFA9
NDUFA10
NDUFAB1
NDUFA11
NDUFA12
NDUFA13
NDUFB1
NDUFB2
NDUFB3
NDUFB4
NDUFB5
NDUFB6
NDUFB7
NDUFB8
NDUFB9
NDUFB10
NDUFB11
NDUFC1
UFC2-KCTD14
K02111
K02112
K02115
K02113
K02114
K02108
K02109
K02110
ATP5F1A
ATP5F1B
ATP5F1C
ATP5F1D
ATP5F1E
ATP5PO
ATP6
ATP5PB
ATP5MC1
ATP5PD
ATP5ME
ATP5MF
ATP5MG
ATP5PF
K02142
K02143
ATP8
SDHC
SDHD
SDHA
SDHB
K00241
K00242...
K00239
K00240
K00244
K00245
K00246
K00247
UQCRFS1
CYTB
CYC1
UQCRC1
UQCRC2
UQCRHL
UQCRB
UQCRQ
UQCR10
UQCR11
K02117
K02118
K02119
K02120
K02121
K02122
K02107
K02123
K02124
COX10
COX3
COX1
COX2
COX4I1
COX5A
COX5B
COX6A1
COX6B2
COX6C
COX7A1
COX7B
COX7C
COX8A
COX11
COX15
COX17
K02301
K02300
K02299
K02298
K02297
K02277
K02276...
K02274...
K02275
K02829
K02828
K02827
K02826
K00404...
K00405...
K00407
K00406
K00425
K00426
ATP6V1A
ATP6V1B1
ATP6V1C2
ATP6V1D
ATP6V1E1
ATP6V1F
ATP6V1G3
ATP6V1H
TCIRG1
ATP6V0C
ATP6V0D2
ATP6V0E2
ATP6AP1

E.

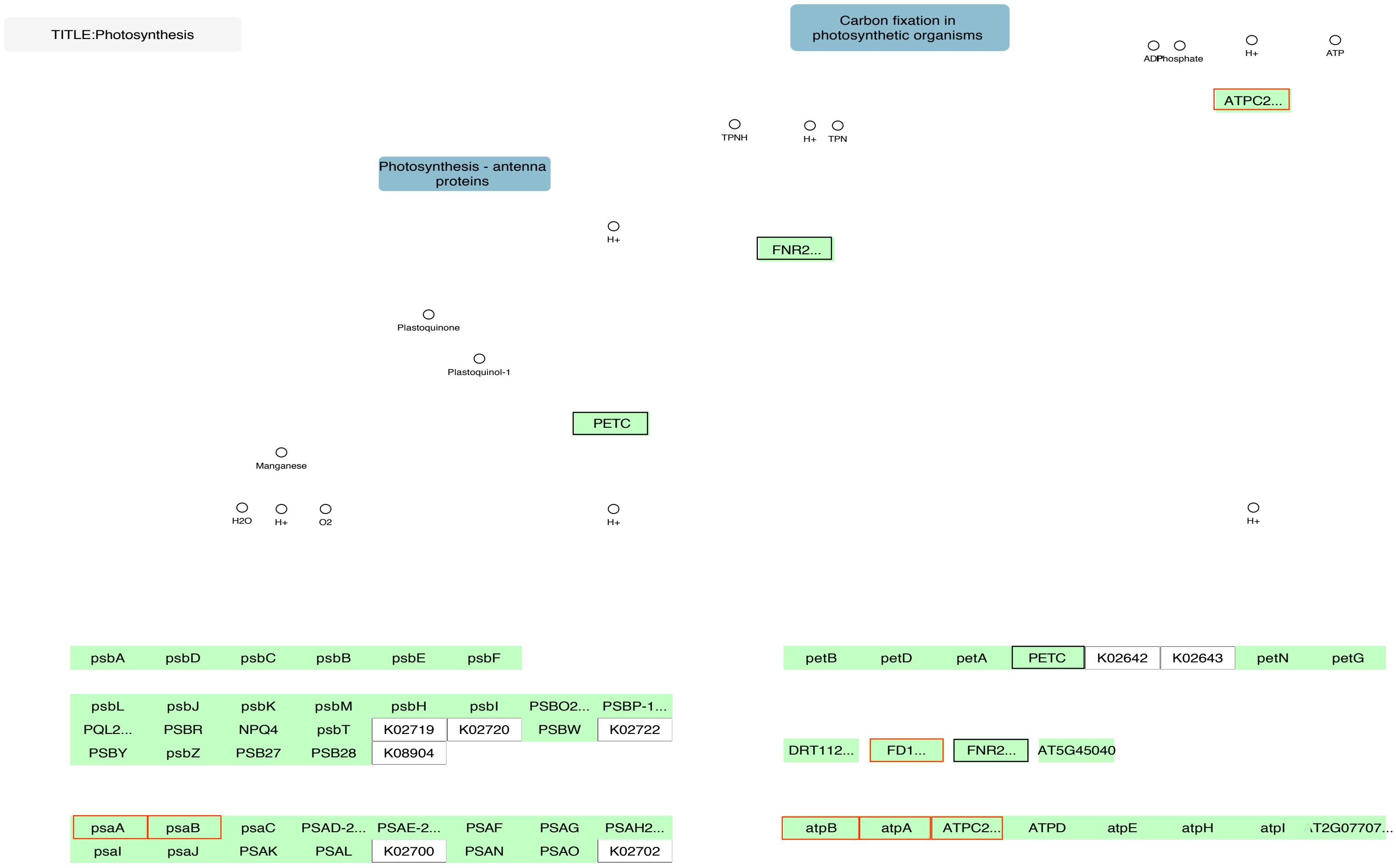
TITLE:Photosynthesis
Carbon fixation in photosynthetic organisms
ADP
Phosphate
H+
ATP
ATPC2...
TPNH
H+
TPN
Photosynthesis - antenna proteins
H+
FNR2...
Plastoquinone
Plastoquinol-1
PETC
Manganese
H2O
H+
O2
H+
H+
psbA psbD psbC psbB psbE psbF
psbL psbJ psbK psbM psbH psbI PSBO2... PSBP-1...
PQL2... PSBR NPQ4 psbT K02719 K02720 PSBW K02722
PSBY psbZ PSB27 PSB28 K08904
psaA psaB psaC PSAD-2... PSAE-2... PSAF PSAG PSAH2...
psaI psaJ PSAK PSAL K02700 PSAN PSAO K02702
petB petD petA PETC K02642 K02643 petN petG
DRT112... FD1... FNR2... AT5G45040
atpB atpA ATPC2... ATPD atpE atpH atpI AT2G07707...

Figure S1. Annotated KEGG pathways for Energy Metabolism. Human pathways for Central metabolism: A. Glycolysis-Gluconeogenesis (hsa00010), B. Citrate cycle (hsa00020), C. Pentose Phosphate pathway (hsa00030), and D. Oxidative Phosphorylation (hsa00190), and Arabidopsis thaliana pathway for E. Photosynthesis (ath00195). Genes in the path- ways that are found in the monophyletic clusters are highlighted with a black outline and complex history clusters are highlighted with a red outline.
