## Supplemental Figure S2 for "Reconstructing and Analysing The Genome of The Last Eukaryote Common Ancestor to Better Understand the Transition from FECA to LECA"

A.

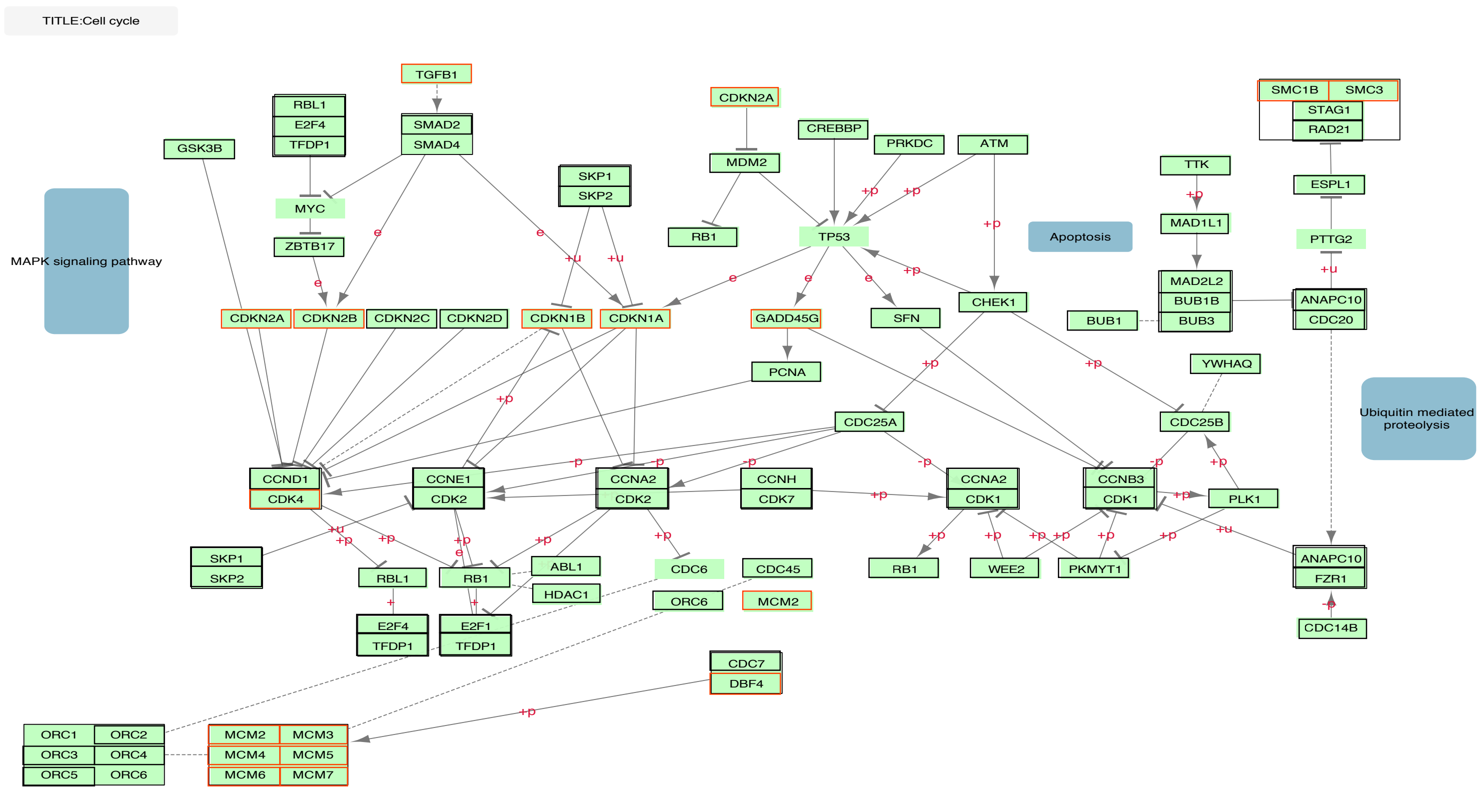

TITLE:Cell cycle
MAPK signaling pathway
TGFB1
SMAD2
SMAD4
RBL1
E2F4
TFDP1
GSK3B
MYC
ZBTB17
CDKN2A
MDM2
RB1
CREBBP
PRKDC
ATM
TP53
SKP1
SKP2
Apoptosis
TTK
MAD1L1
MAD2L2
BUB1B
BUB3
BUB1
SMC1B
SMC3
STAG1
RAD21
ESPL1
PTTG2
ANAPC10
CDC20
CDKN2A
CDKN2B
CDKN2C
CDKN2D
CDKN1B
CDKN1A
GADD45G
SFN
CHEK1
PCNA
YWHAQ
CDC25A
CDC25B
Ubiquitin mediated proteolysis
CCND1
CDK4
CCNE1
CDK2
CCNA2
CDK2
CCNH
CDK7
CCNA2
CDK1
CCNB3
CDK1
PLK1
SKP1
SKP2
RBL1
RB1
ABL1
HDAC1
CDC6
CDC45
ORC6
MCM2
RB1
WEE2
PKMYT1
ANAPC10
FZR1
CDC14B
E2F4
TFDP1
E2F1
TFDP1
CDC7
DBF4
ORC1
ORC2
ORC3
ORC4
ORC5
ORC6
MCM2
MCM3
MCM4
MCM5
MCM6
MCM7

B.

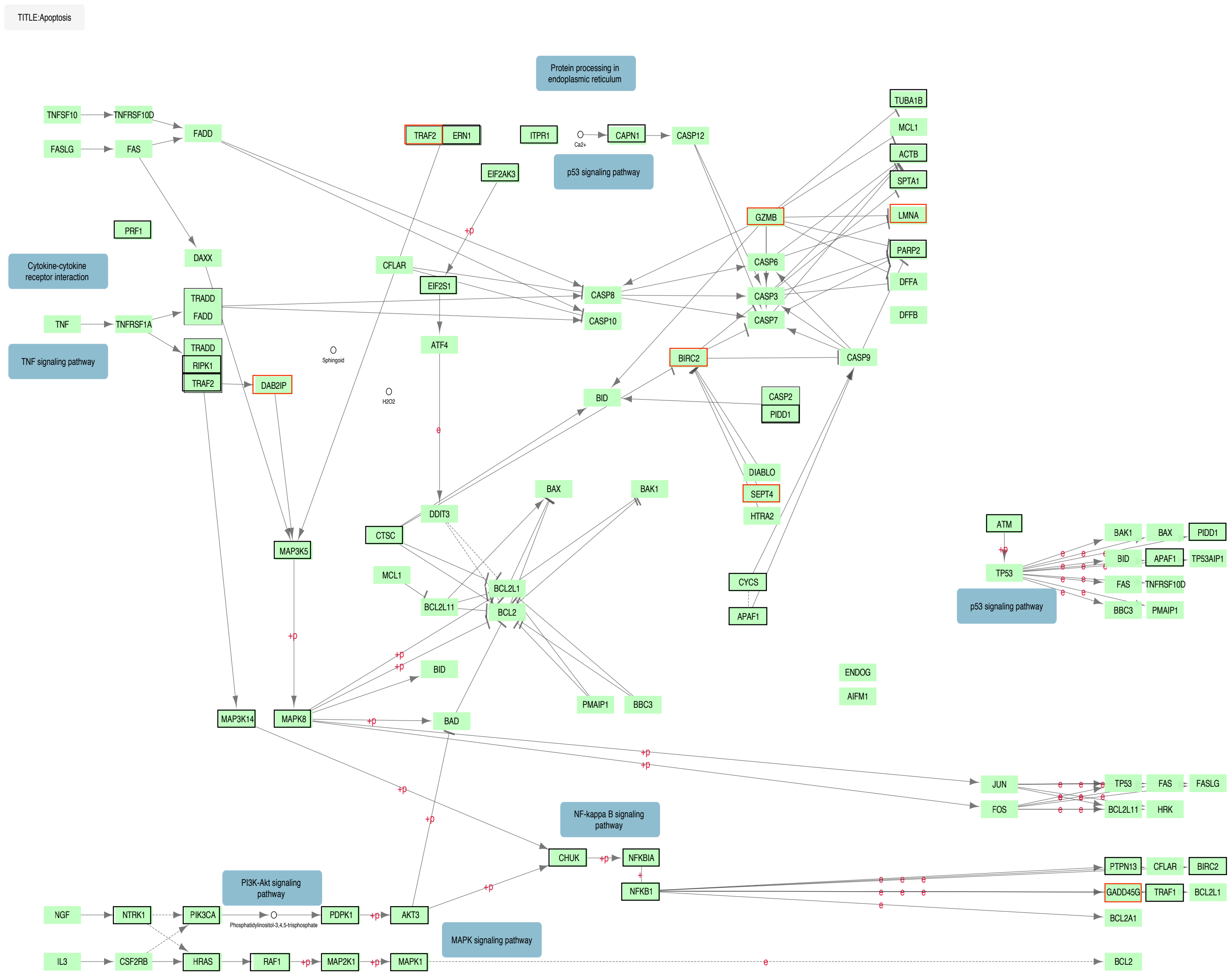
TITLE:Apoptosis
Protein processing in endoplasmic reticulum
p53 signaling pathway
Cytokine-cytokine receptor interaction
TNF signaling pathway
NF-kappa B signaling pathway
PI3K-Akt signaling pathway
MAPK signaling pathway
TNFSF10
TNFRSF10D
FASLG
FAS
FADD
PRF1
DAXX
TRADD
TNF
TNFRSF1A
RIPK1
TRAF2
DAB2IP
ERN1
ITPR1
Ca2+
CAPN1
CASP12
EIF2AK3
CFLAR
EIF2S1
ATF4
Sphingoid
H2O2
CASP8
CASP10
GZMB
CASP6
CASP3
CASP7
TUBA1B
MCL1
ACTB
SPTA1
LMNA
PARP2
DFFA
DFFB
BIRC2
CASP9
BID
CASP2
PIDD1
DIABLO
SEPT4
HTRA2
CYCS
APAF1
DDIT3
CTSC
BAX
BAK1
BCL2L1
BCL2
BCL2L11
MAP3K5
PMAIP1
BBC3
BAD
ENDOG
AIFM1
MAP3K14
MAPK8
ATM
TP53
TP53AIP1
JUN
FOS
HRK
CHUK
NFKBIA
NFKB1
PTPN13
GADD45G
TRAF1
BCL2A1
NGF
NTRK1
PIK3CA
Phosphatidylinositol-3,4,5-trisphosphate
PDPK1
AKT3
IL3
CSF2RB
HRAS
RAF1
MAP2K1
MAPK1
+p
e

C.

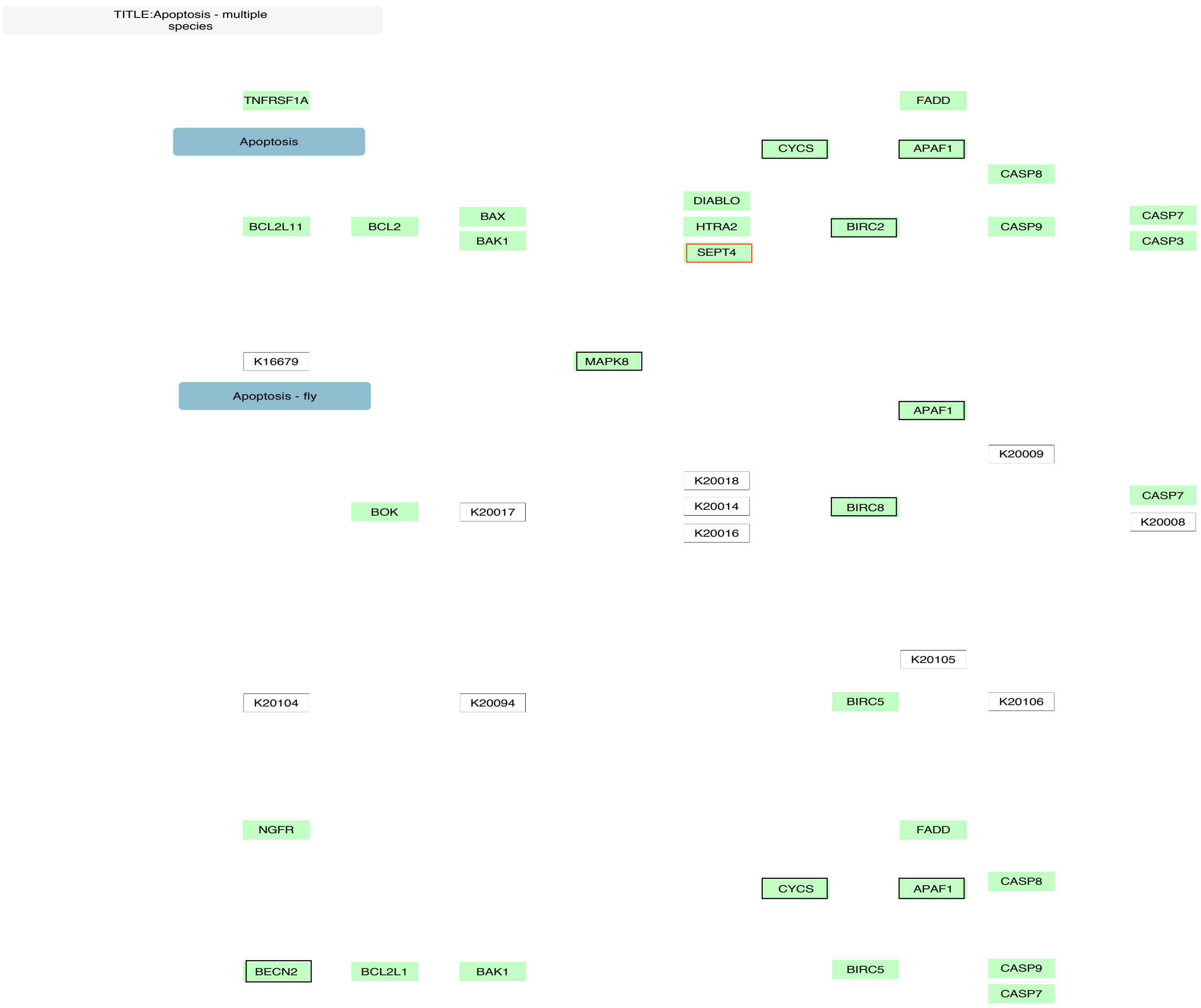

TITLE:Apoptosis - multiple species
TNFRSF1A
Apoptosis
FADD
CYCS
APAF1
CASP8
DIABLO
BCL2L11
BCL2
BAX
BAK1
HTRA2
SEPT4
BIRC2
CASP9
CASP7
CASP3
K16679
MAPK8
Apoptosis - fly
APAF1
K20009
K20018
BOK
K20017
K20014
K20016
BIRC8
CASP7
K20008
K20105
K20104
K20094
BIRC5
K20106
NGFR
FADD
CYCS
APAF1
CASP8
BECN2
BCL2L1
BAK1
BIRC5
CASP9
CASP7

D.

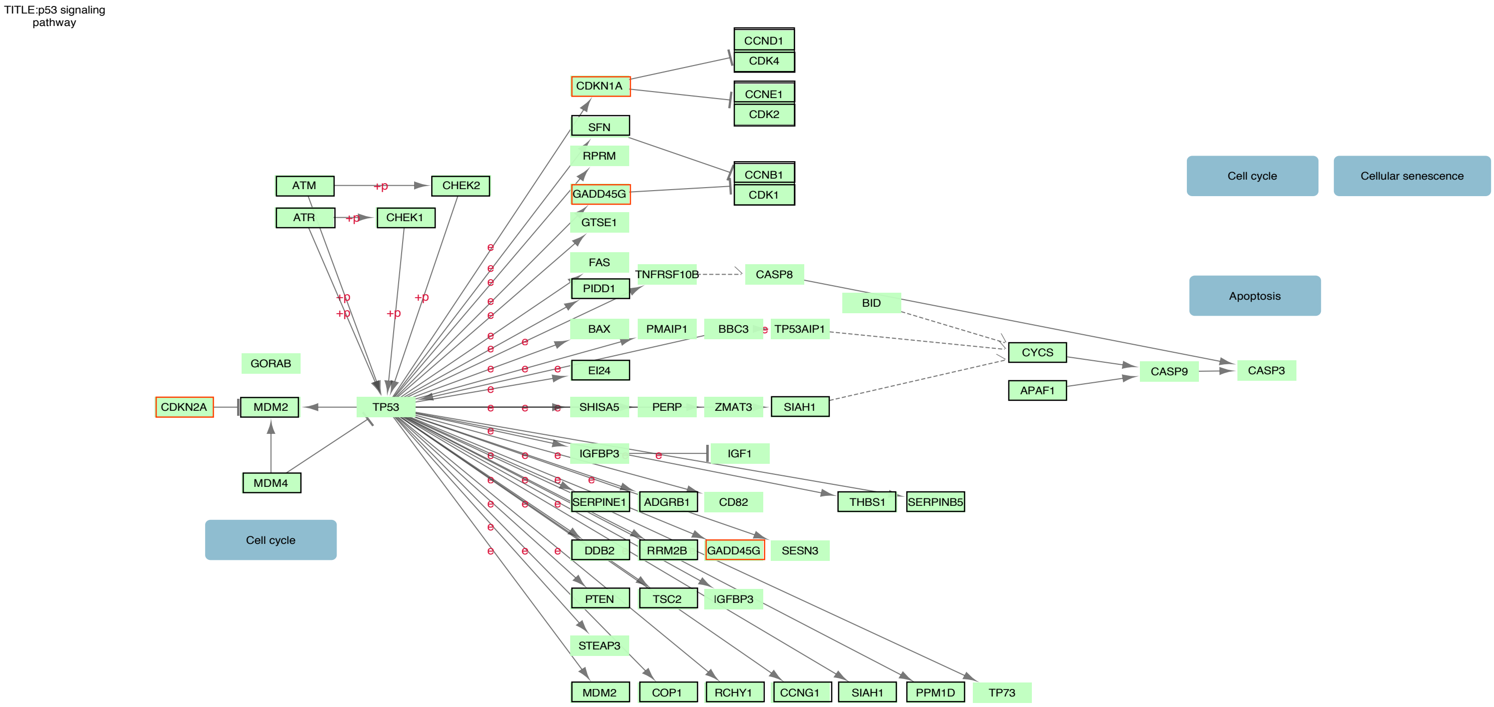
TITLE:p53 signaling pathway
CCND1
CDK4
CDKN1A
CCNE1
CDK2
SFN
RPRM
CCNB1
CDK1
GADD45G
ATM
CHEK2
ATR
CHEK1
GTSE1
Cell cycle
Cellular senescence
FAS
TNFRSF10B
CASP8
PIDD1
BID
Apoptosis
BAX
PMAIP1
BBC3
TP53AIP1
CYCS
GORAB
EI24
CASP9
CASP3
APAF1
CDKN2A
MDM2
TP53
SHISA5
PERP
ZMAT3
SIAH1
IGFBP3
IGF1
MDM4
SERPINE1
ADGRB1
CD82
THBS1
SERPINB5
Cell cycle
DDB2
RRM2B
GADD45G
SESN3
PTEN
TSC2
IGFBP3
STEAP3
MDM2
COP1
RCHY1
CCNG1
SIAH1
PPM1D
TP73
+p
e

E.

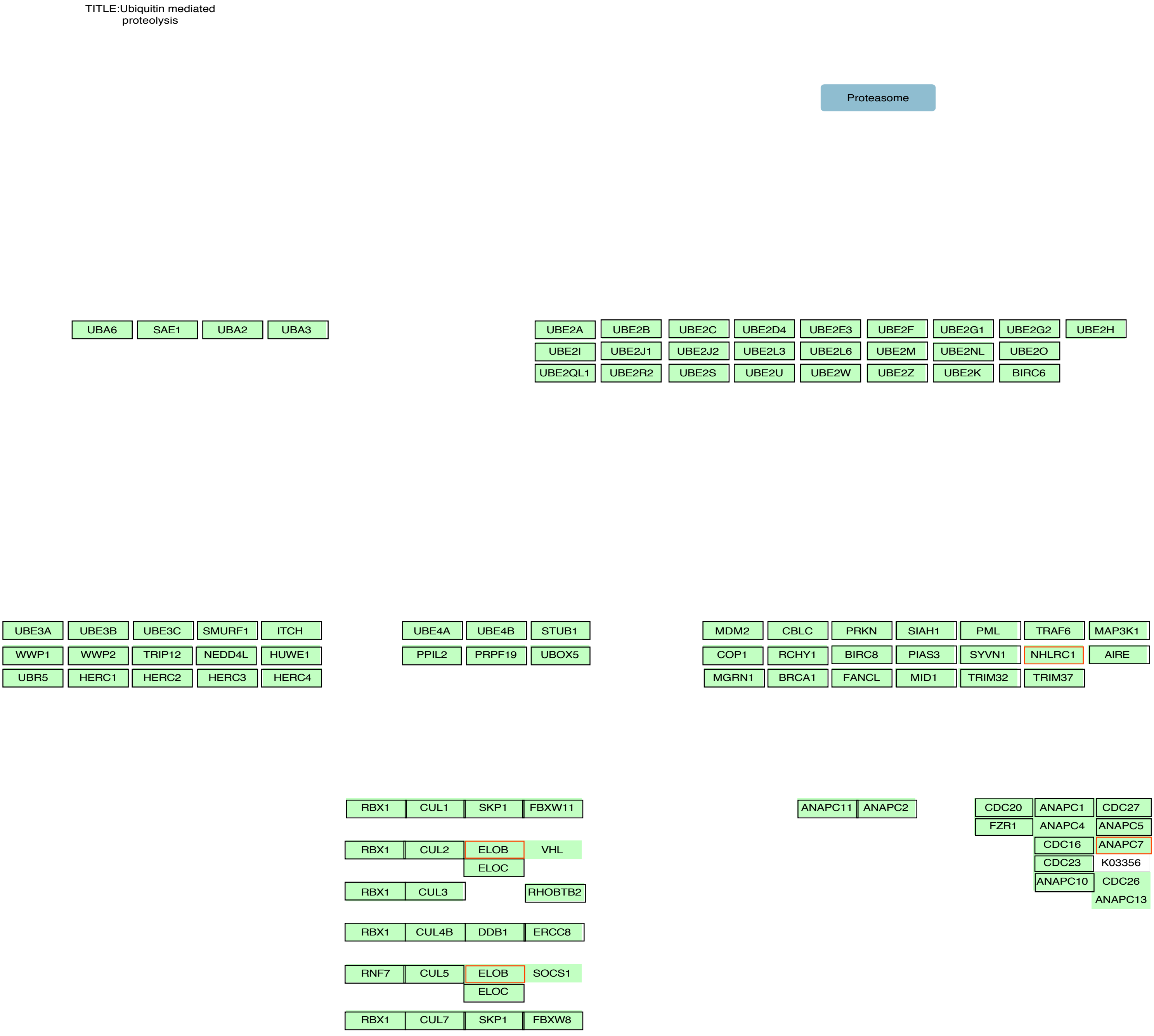
TITLE:Ubiquitin mediated proteolysis
Proteasome
UBA6 SAE1 UBA2 UBA3
UBE2A UBE2B UBE2C UBE2D4 UBE2E3 UBE2F UBE2G1 UBE2G2 UBE2H
UBE2I UBE2J1 UBE2J2 UBE2L3 UBE2L6 UBE2M UBE2NL UBE2O
UBE2QL1 UBE2R2 UBE2S UBE2U UBE2W UBE2Z UBE2K BIRC6
UBE3A UBE3B UBE3C SMURF1 ITCH
WWP1 WWP2 TRIP12 NEDD4L HUWE1
UBR5 HERC1 HERC2 HERC3 HERC4
UBE4A UBE4B STUB1
PPIL2 PRPF19 UBOX5
MDM2 CBLC PRKN SIAH1 PML TRAF6 MAP3K1
COP1 RCHY1 BIRC8 PIAS3 SYVN1 NHLRC1 AIRE
MGRN1 BRCA1 FANCL MID1 TRIM32 TRIM37
RBX1 CUL1 SKP1 FBXW11
RBX1 CUL2 ELOB VHL
ELOC
RBX1 CUL3 RHOBTB2
RBX1 CUL4B DDB1 ERCC8
RNF7 CUL5 ELOB SOCS1
ELOC
RBX1 CUL7 SKP1 FBXW8
ANAPC11 ANAPC2
CDC20 ANAPC1 CDC27
FZR1 ANAPC4 ANAPC5
CDC16 ANAPC7
CDC23 K03356
ANAPC10 CDC26
ANAPC13

Figure S2. Annotated KEGG pathways for Mitosis and the Cell cycle. Human pathways for Mitosis and the cell cycle: A. Cell cycle (hsa04110), B. Apoptosis (hsa04210), C. Apoptosis (hsa04215), D. p53 signalling pathway (hsa04115), and E. Ubiquitin mediated proteolysis (hsa04120). Genes in the pathways that are found in the monophyletic clusters are highlighted with a black outline and complex history clusters are highlighted with a red outline.
