## Supplemental Figure S3 for "Reconstructing and Analysing The Genome of The Last Eukaryote Common Ancestor to Better Understand the Transition from FECA to LECA"

A.

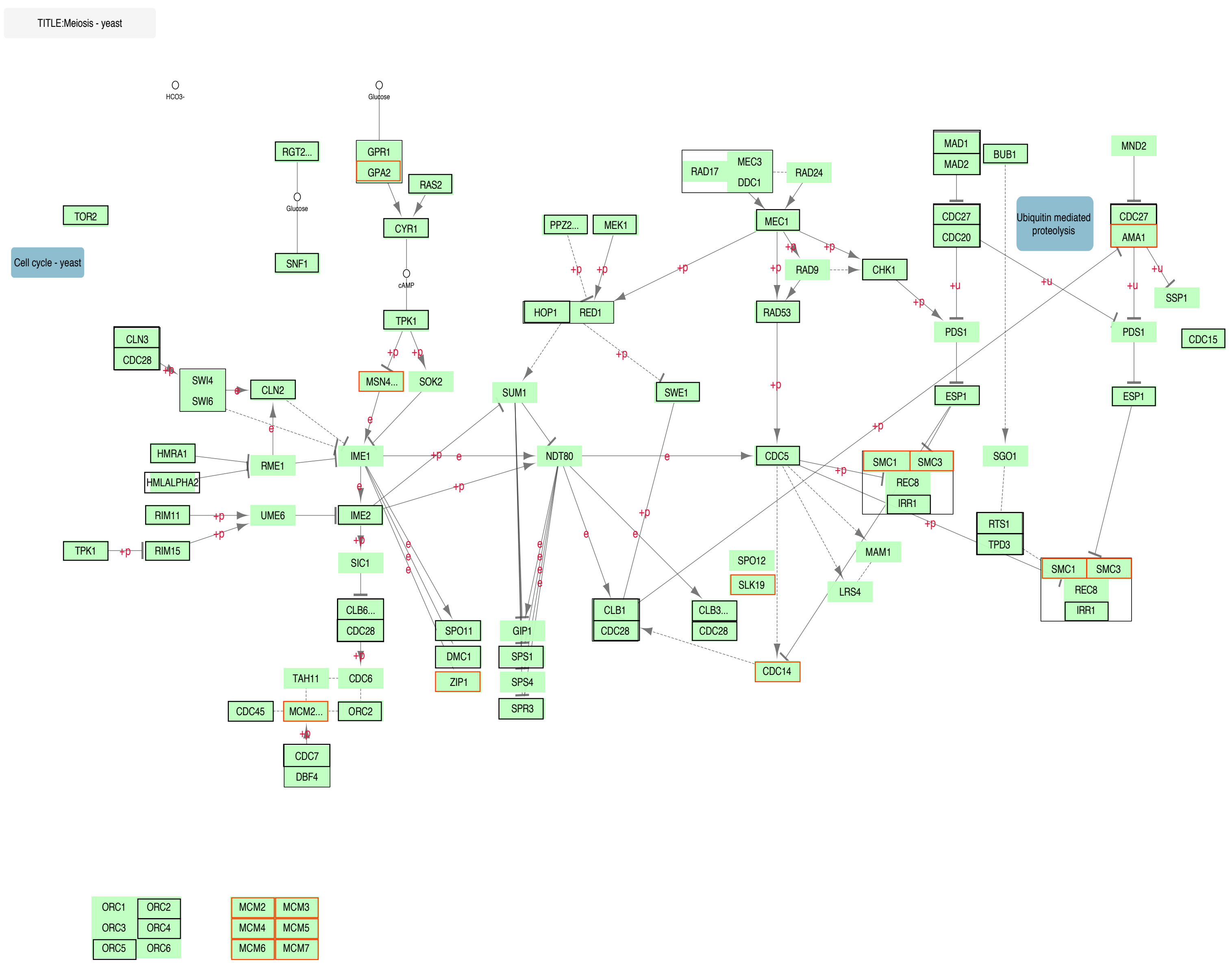
TITLE:Meiosis - yeast
HCO3-
Glucose
RGT2...
GPR1
GPA2
RAS2
Glucose
CYR1
SNF1
TOR2
Cell cycle - yeast
cAMP
TPK1
CLN3
CDC28
SWI4
SWI6
CLN2
MSN4...
SOK2
HMRA1
HMLALPHA2
RME1
IME1
RIM11
UME6
IME2
TPK1
RIM15
SIC1
CLB6...
CDC28
TAH11
CDC6
CDC45
MCM2...
ORC2
CDC7
DBF4
PPZ2...
MEK1
HOP1
RED1
SUM1
NDT80
SWE1
SPO11
DMC1
ZIP1
GIP1
SPS1
SPS4
SPR3
CLB1
CDC28
CLB3...
CDC28
RAD17
MEC3
DDC1
RAD24
MEC1
RAD9
CHK1
RAD53
CDC5
SPO12
SLK19
CDC14
MAM1
LRS4
MAD1
MAD2
BUB1
CDC27
CDC20
Ubiquitin mediated proteolysis
MND2
CDC27
AMA1
SSP1
PDS1
PDS1
CDC15
ESP1
ESP1
SMC1
SMC3
REC8
IRR1
SGO1
RTS1
TPD3
SMC1
SMC3
REC8
IRR1
ORC1
ORC2
ORC3
ORC4
ORC5
ORC6
MCM2
MCM3
MCM4
MCM5
MCM6
MCM7

B.

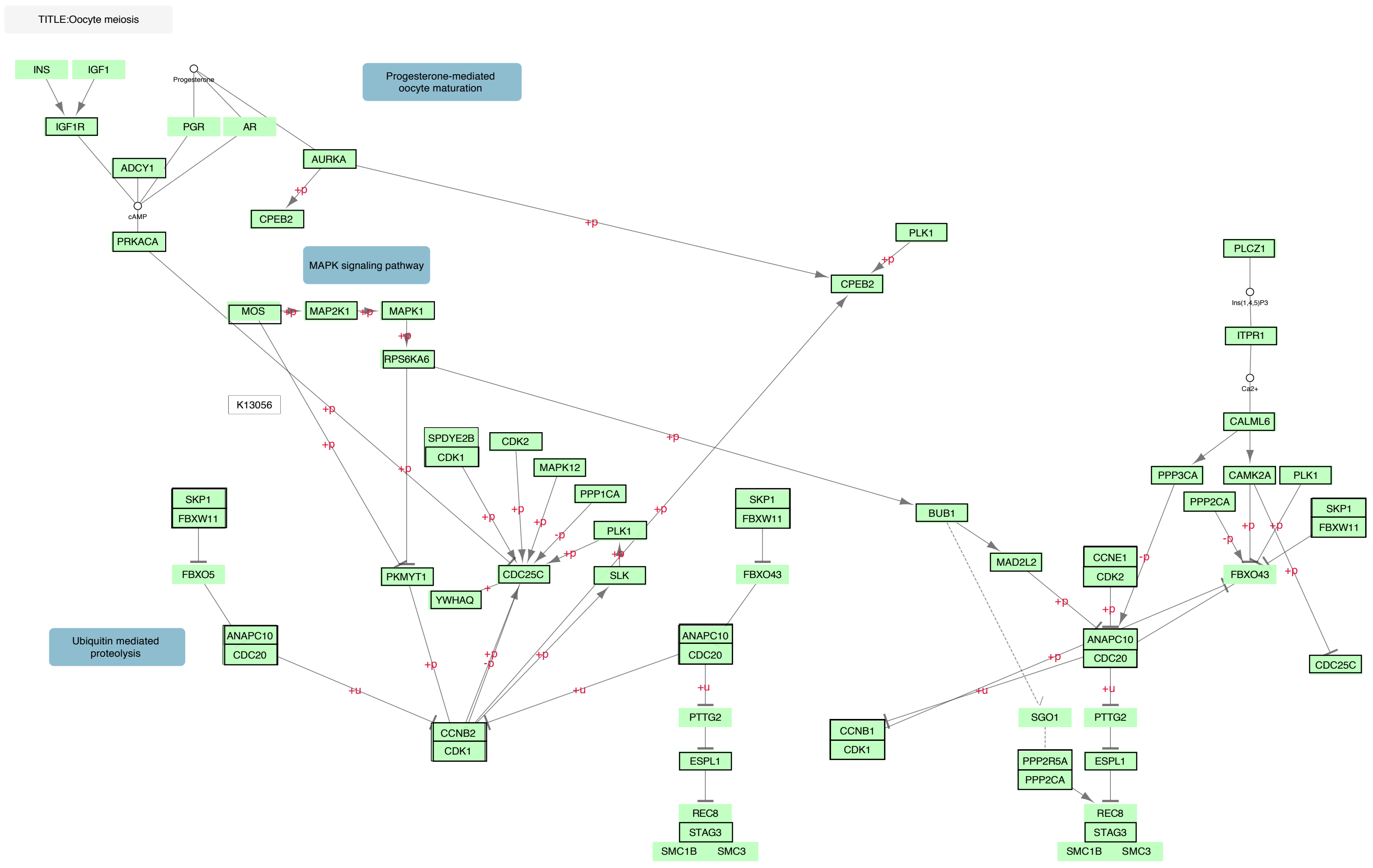
TITLE:Oocyte meiosis
Progesterone-mediated oocyte maturation
MAPK signaling pathway
Ubiquitin mediated proteolysis
INS
IGF1
IGF1R
Progesterone
PGR
AR
ADCY1
cAMP
PRKACA
AURKA
CPEB2
PLK1
MOS
MAP2K1
MAPK1
RPS6KA6
K13056
SPDYE2B
CDK1
CDK2
MAPK12
PPP1CA
SLK
CDC25C
YWHAQ
PKMYT1
SKP1
FBXW11
FBXO5
ANAPC10
CDC20
CCNB2
FBXO43
PTTG2
ESPL1
REC8
STAG3
SMC1B
SMC3
BUB1
MAD2L2
CCNE1
CCNB1
SGO1
PPP2R5A
PPP2CA
PLCZ1
Ins(1,4,5)P3
ITPR1
Ca2+
CALML6
PPP3CA
CAMK2A
+p
-p
+u

C.

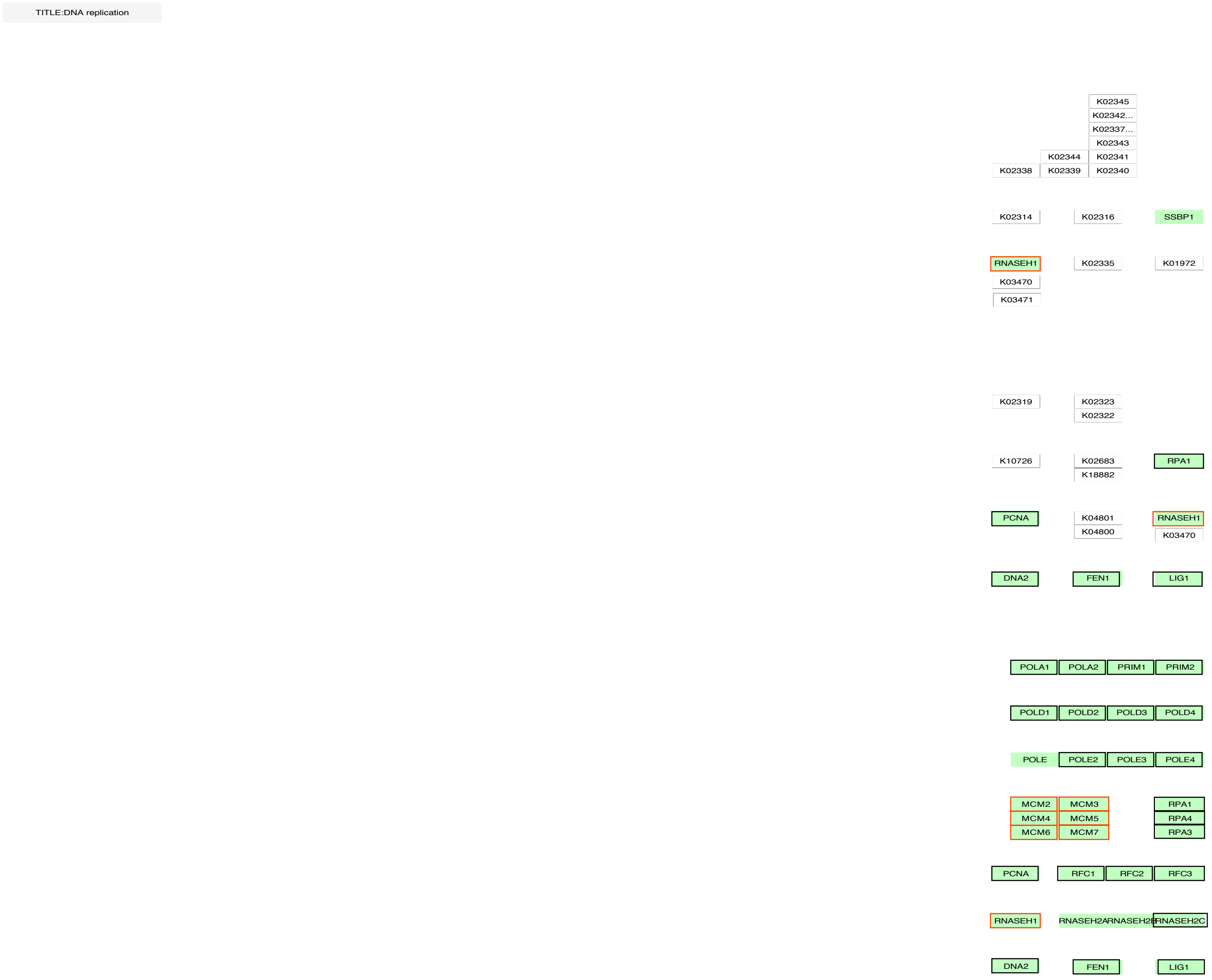
TITLE:DNA replication
K02345
K02342...
K02337...
K02343
K02344
K02341
K02338
K02339
K02340
K02314
K02316
SSBP1
RNASEH1
K02335
K01972
K03470
K03471
K02319
K02323
K02322
K10726
K02683
K18882
RPA1
PCNA
K04801
K04800
RNASEH1
K03470
DNA2
FEN1
LIG1
POLA1
POLA2
PRIM1
PRIM2
POLD1
POLD2
POLD3
POLD4
POLE
POLE2
POLE3
POLE4
MCM2
MCM3
MCM4
MCM5
MCM6
MCM7
RPA1
RPA4
RPA3
PCNA
RFC1
RFC2
RFC3
RNASEH1
RNASEH2A
RNASEH2B
RNASEH2C
DNA2
FEN1
LIG1

D.

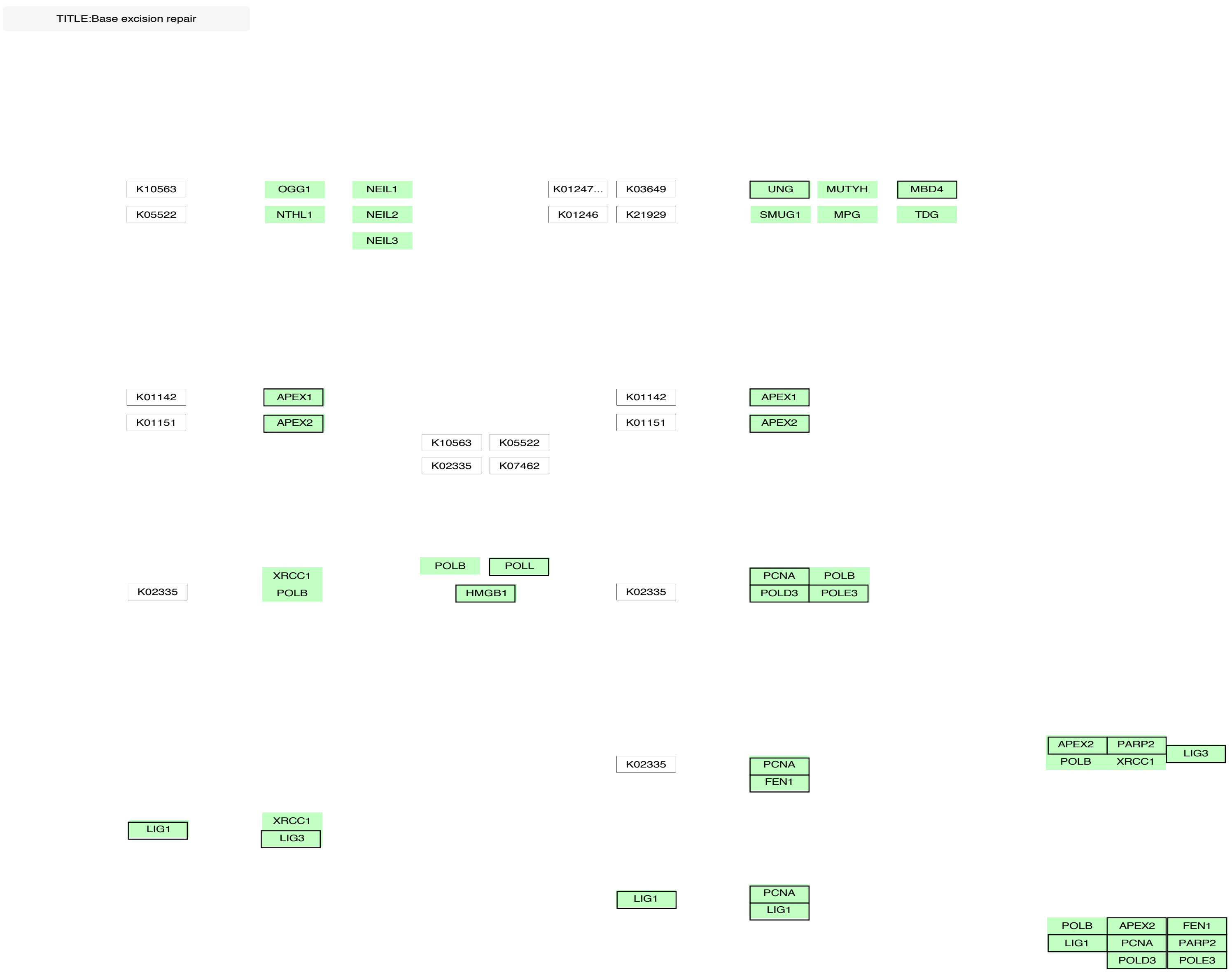
TITLE:Base excision repair
K10563
K05522
OGG1
NTHL1
NEIL1
NEIL2
NEIL3
K01247...
K01246
K03649
K21929
UNG
SMUG1
MUTYH
MPG
MBD4
TDG
K01142
K01151
APEX1
APEX2
K10563
K05522
K02335
K07462
K01142
K01151
APEX1
APEX2
K02335
XRCC1
POLB
POLB
POLL
HMGB1
K02335
PCNA
POLB
POLD3
POLE3
K02335
PCNA
FEN1
APEX2
PARP2
POLB
XRCC1
LIG3
LIG1
XRCC1
LIG3
LIG1
PCNA
LIG1
POLB
APEX2
FEN1
LIG1
PCNA
PARP2
POLD3
POLE3

E.

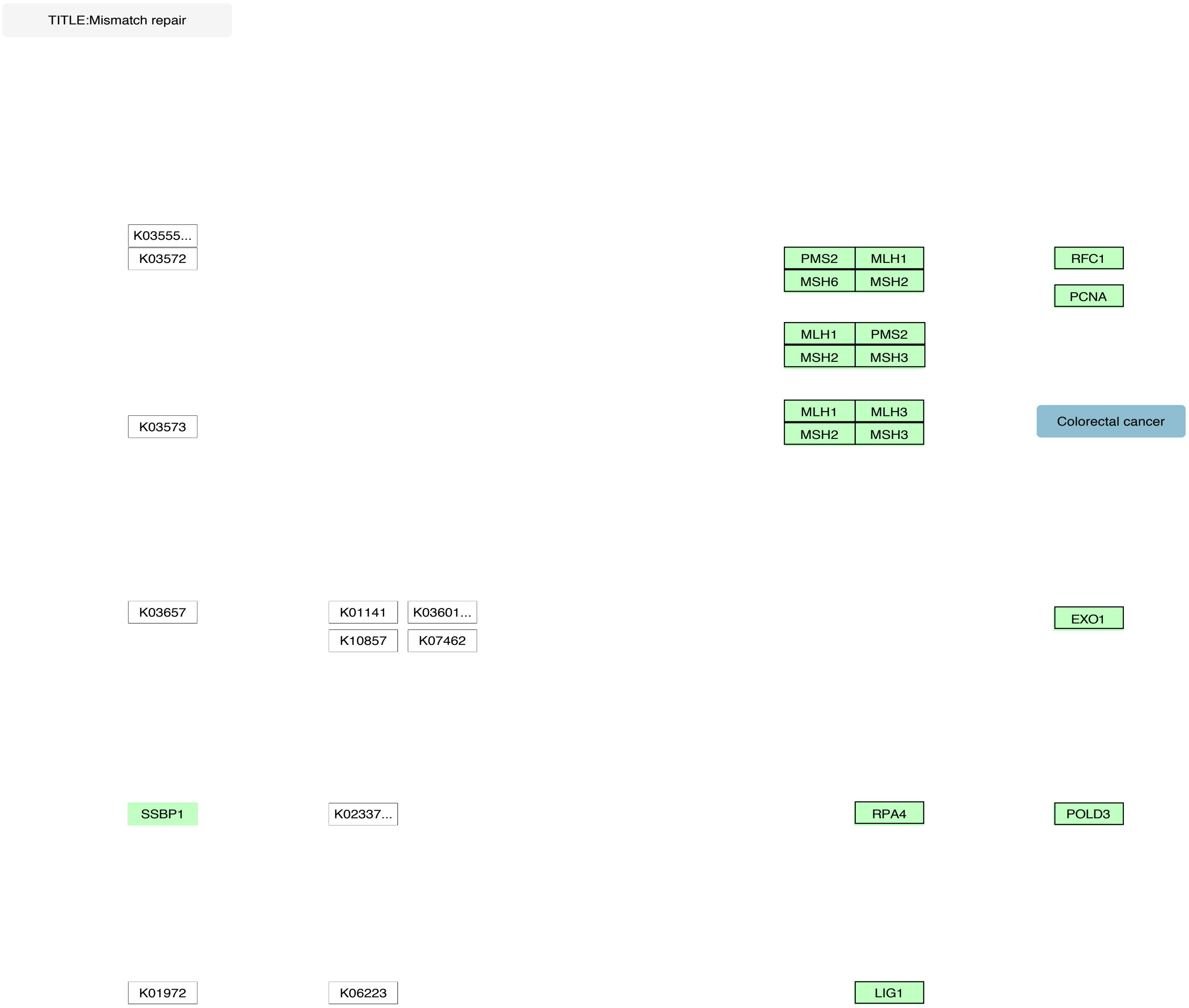
TITLE:Mismatch repair
K03555...
K03572
PMS2
MLH1
MSH6
MSH2
RFC1
PCNA
MLH1
PMS2
MSH2
MSH3
K03573
MLH1
MLH3
MSH2
MSH3
Colorectal cancer
K03657
K01141
K03601...
K10857
K07462
EXO1
SSBP1
K02337...
RPA4
POLD3
K01972
K06223
LIG1

F.

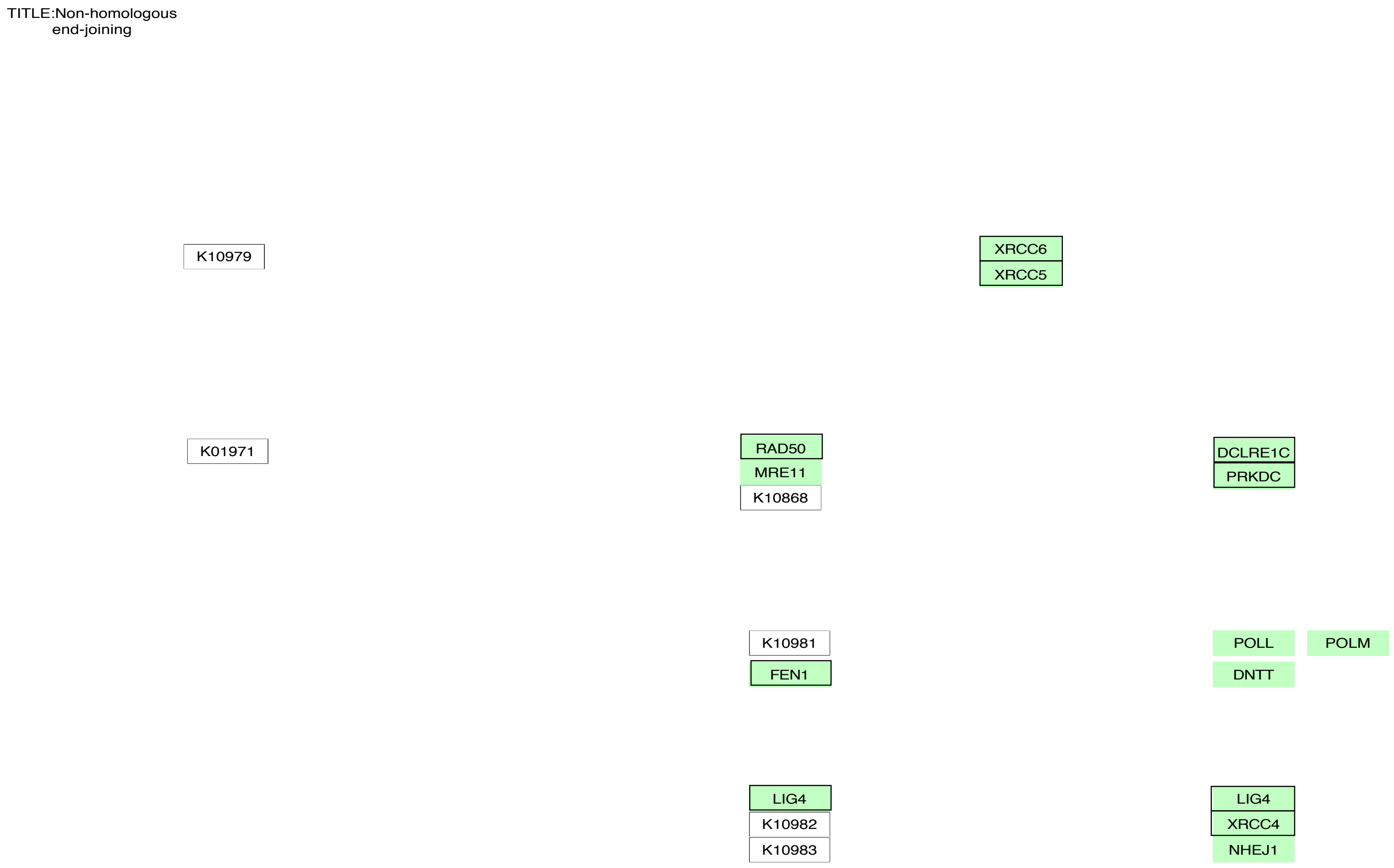
TITLE:Non-homologous end-joining
K10979
XRCC6
XRCC5
K01971
RAD50
MRE11
K10868
DCLRE1C
PRKDC
K10981
FEN1
POLL
POLM
DNTT
LIG4
K10982
K10983
LIG4
XRCC4
NHEJ1

G.

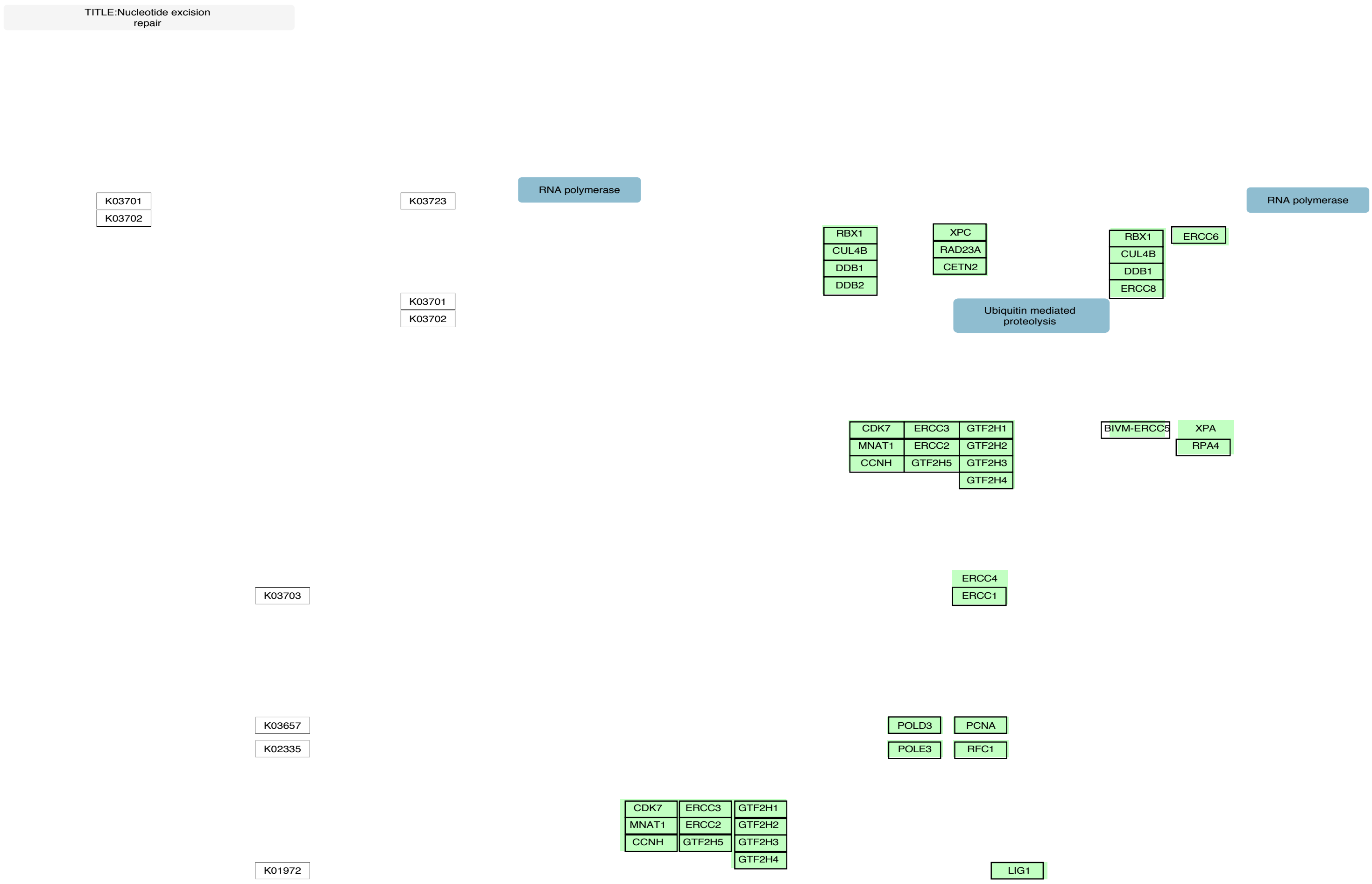
TITLE:Nucleotide excision repair
K03701
K03702
K03723
RNA polymerase
K03701
K03702
RBX1
CUL4B
DDB1
DDB2
XPC
RAD23A
CETN2
Ubiquitin mediated proteolysis
RBX1
CUL4B
DDB1
ERCC8
ERCC6
RNA polymerase
CDK7
MNAT1
CCNH
ERCC3
ERCC2
GTF2H5
GTF2H1
GTF2H2
GTF2H3
GTF2H4
BIVM-ERCC5
XPA
RPA4
K03703
ERCC4
ERCC1
K03657
K02335
POLD3
POLE3
PCNA
RFC1
CDK7
MNAT1
CCNH
ERCC3
ERCC2
GTF2H5
GTF2H1
GTF2H2
GTF2H3
GTF2H4
K01972
LIG1

Figure S3. Annotated KEGG pathways for Meiosis and the Cell cycle. Yeast and human pathways for Meiosis and the cell cycle: A. Meiosis yeast (sce04113), B. Oocyte meiosis (hsa04114), C. DNA replication (hsa03030), D. Base excision repair (hsa03410), E. Mismatch repair (hsa03430), F. Non-homologous end joining (hsa03450), and G. Nucleotide excision repair (hsa03420). Genes in the pathways that are found in the monophyletic clusters are highlighted with a black outline and complex history clusters are highlighted with a red outline.
