## Supplemental Figure S4 for "Reconstructing and Analysing The Genome of The Last Eukaryote Common Ancestor to Better Understand the Transition from FECA to LECA"

A.

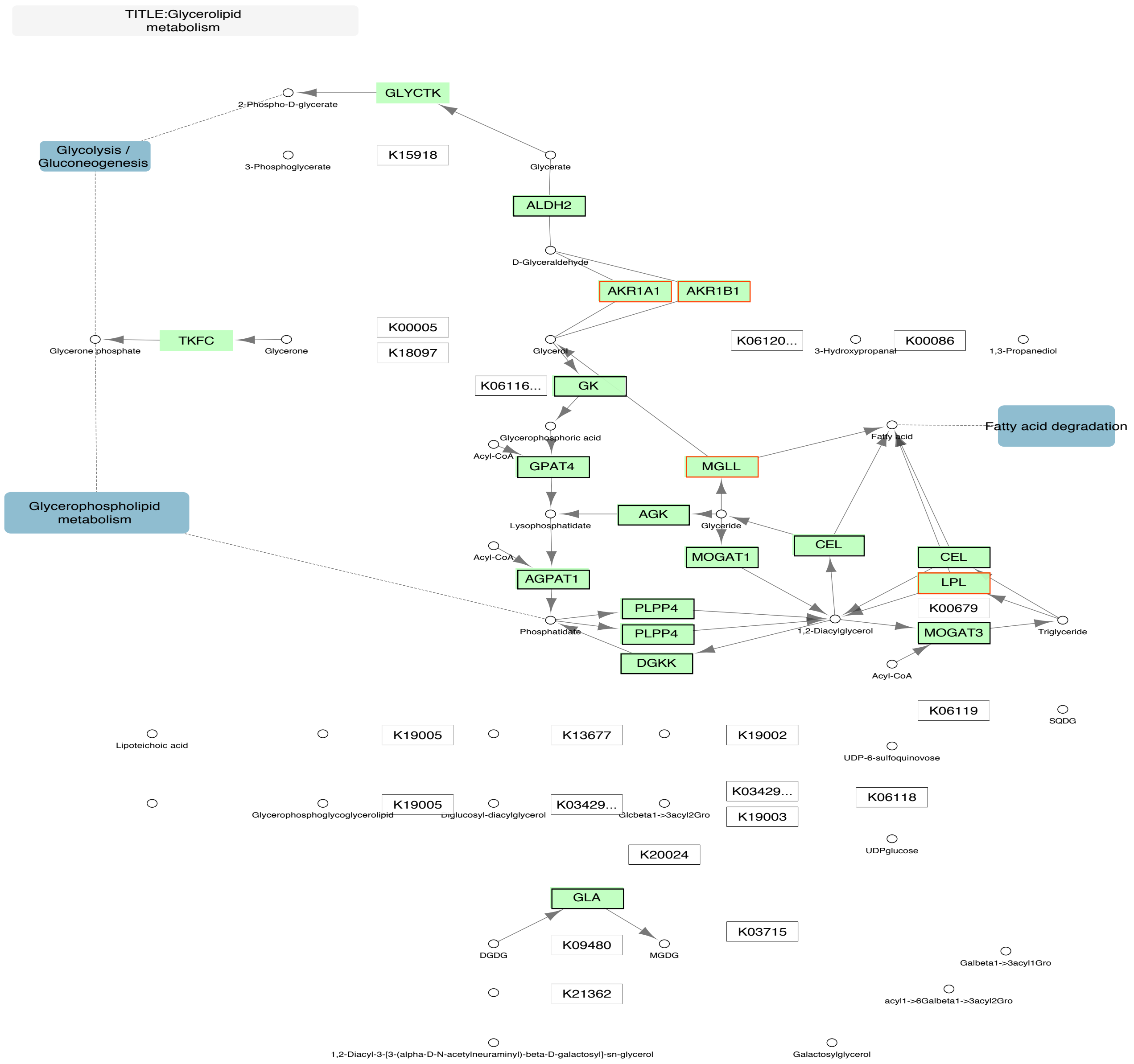
TITLE:Glycerolipid metabolism
GLYCTK
2-Phospho-D-glycerate
Glycolysis / Gluconeogenesis
3-Phosphoglycerate
K15918
Glycerate
ALDH2
D-Glyceraldehyde
AKR1A1
AKR1B1
Glycerone phosphate
TKFC
Glycerone
K00005
K18097
Glycerol
K06120...
3-Hydroxypropanal
K00086
1,3-Propanediol
K06116...
GK
Fatty acid
Fatty acid degradation
Glycerophosphoric acid
Acyl-CoA
GPAT4
MGLL
Glycerophospholipid metabolism
AGK
Lysophosphatidate
Glyceride
CEL
Acyl-CoA
MOGAT1
CEL
LPL
AGPAT1
PLPP4
K00679
Phosphatidate
PLPP4
1,2-Diacylglycerol
MOGAT3
Triglyceride
DGKK
Acyl-CoA
K06119
SQDG
Lipoteichoic acid
K19005
K13677
K19002
UDP-6-sulfoquinovose
K03429...
K06118
Glycerophosphoglycoglycerolipid
K19005
Diglucosyl-diacylglycerol
K03429...
Glcbeta1->3acyl2Gro
K19003
K20024
UDPglucose
GLA
K03715
DGDG
K09480
MGDG
Galbeta1->3acyl1Gro
K21362
acyl1->6Galbeta1->3acyl2Gro
1,2-Diacyl-3-[3-(alpha-D-N-acetylneuraminyl)-beta-D-galactosyl]-sn-glycerol
Galactosylglycerol

B.

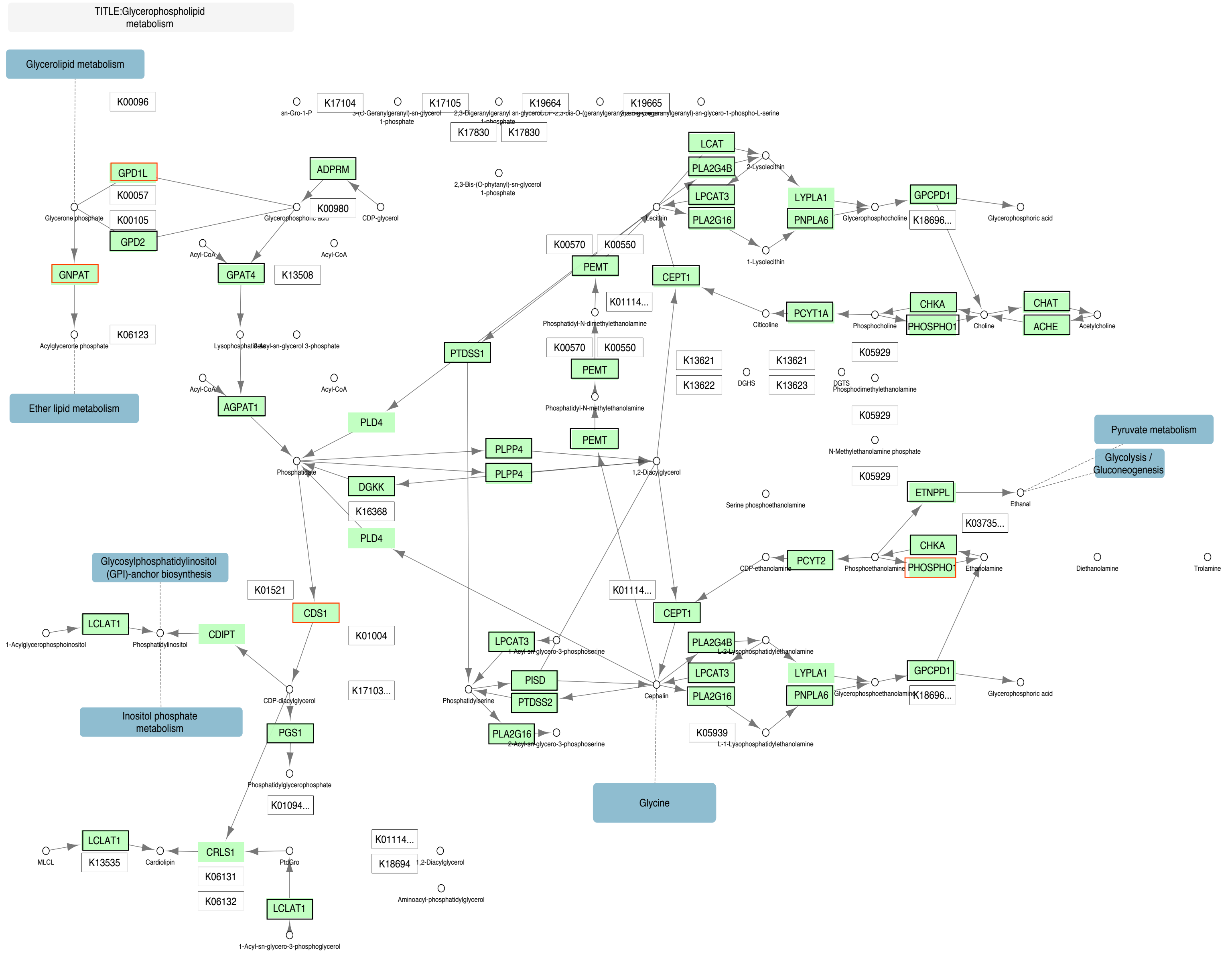
TITLE:Glycerophospholipid metabolism
Glycerolipid metabolism
K00096
GPD1L
K00057
K00105
GPD2
Glycerone phosphate
GNPAT
Acylglycerone phosphate
K06123
Ether lipid metabolism
ADPRM
K00980
Glycerophosphoric acid
CDP-glycerol
Acyl-CoA
GPAT4
K13508
Lysophosphatidate
1-Acyl-sn-glycerol 3-phosphate
AGPAT1
sn-Gro-1-P
K17104
K17105
K19664
K19665
K17830
2,3-Bis-(O-phytanyl)-sn-glycerol 1-phosphate
LCAT
PLA2G4B
2-Lysolecithin
LPCAT3
PLA2G16
Lecithin
LYPLA1
PNPLA6
1-Lysolecithin
Glycerophosphocholine
GPCPD1
K18696...
Glycerophosphoric acid
K00570
K00550
PEMT
CEPT1
K01114...
Phosphatidyl-N-dimethylethanolamine
Phosphatidyl-N-methylethanolamine
Citicoline
PCYT1A
Phosphocholine
CHKA
PHOSPHO1
Choline
CHAT
ACHE
Acetylcholine
K05929
K13621
K13622
K13623
DGHS
DGTS
Phosphodimethylethanolamine
N-Methylethanolamine phosphate
PTDSS1
PLD4
PLPP4
DGKK
K16368
Phosphatidate
1,2-Diacylglycerol
Serine phosphoethanolamine
ETNPPL
Ethanal
K03735...
Pyruvate metabolism
Glycolysis / Gluconeogenesis
CDP-ethanolamine
PCYT2
Phosphoethanolamine
Ethanolamine
Diethanolamine
Trolamine
Glycosylphosphatidylinositol (GPI)-anchor biosynthesis
K01521
CDS1
K01004
K17103...
LCLAT1
1-Acylglycerophosphoinositol
Phosphatidylinositol
CDIPT
CDP-diacylglycerol
Inositol phosphate metabolism
PGS1
Phosphatidylglycerophosphate
K01094...
LPCAT3
1-Acyl-sn-glycero-3-phosphoserine
PISD
PTDSS2
Phosphatidylserine
PLA2G16
2-Acyl-sn-glycero-3-phosphoserine
Cephalin
2-Lysophosphatidylethanolamine
Glycerophosphoethanolamine
K05939
L-1-Lysophosphatidylethanolamine
Glycine
MLCL
K13535
Cardiolipin
CRLS1
K06131
K06132
PtdGro
1-Acyl-sn-glycero-3-phosphoglycerol
K01114...
K18694
Aminoacyl-phosphatidylglycerol

C.

TITLE:Inositol phosphate metabolism

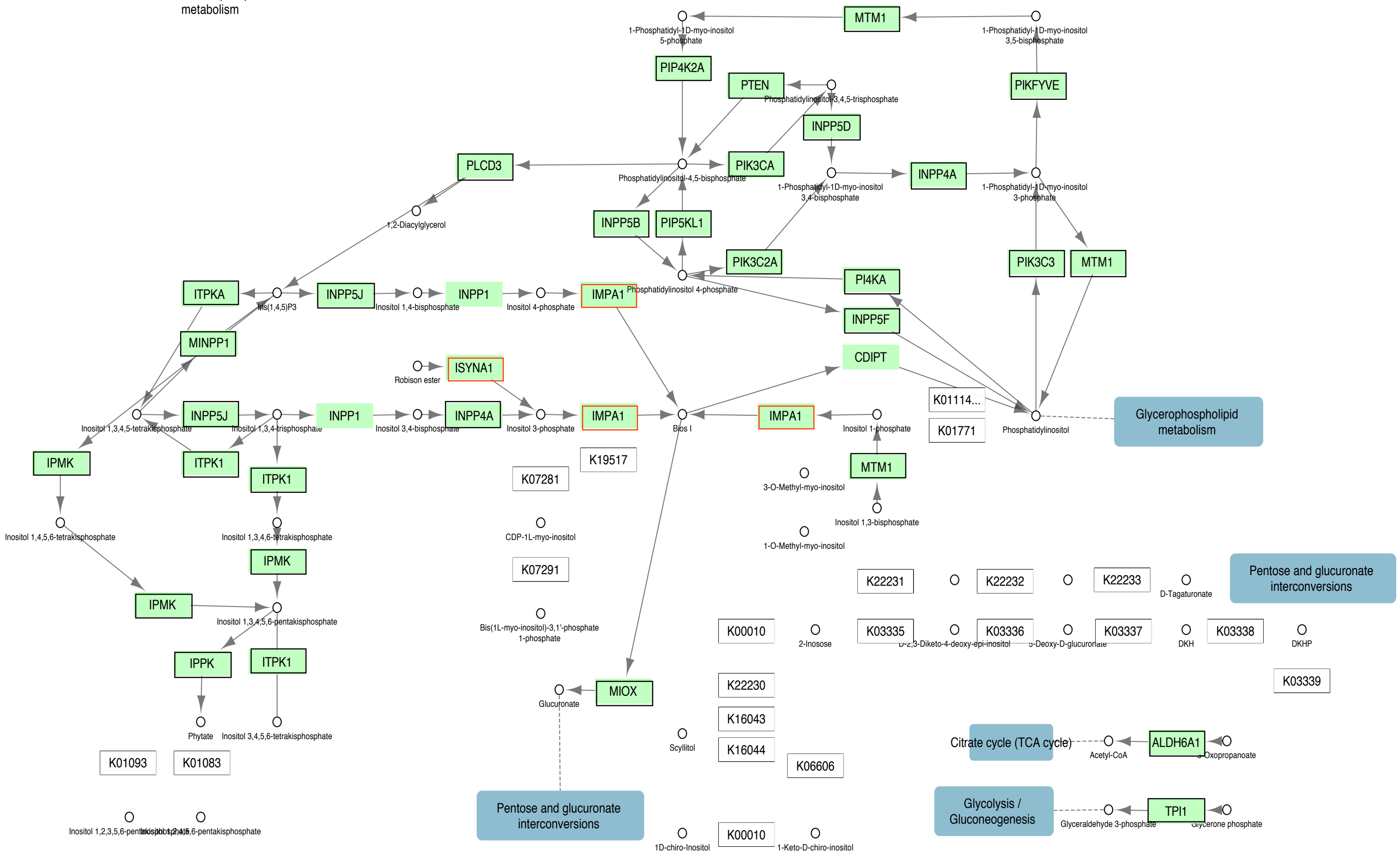

D.

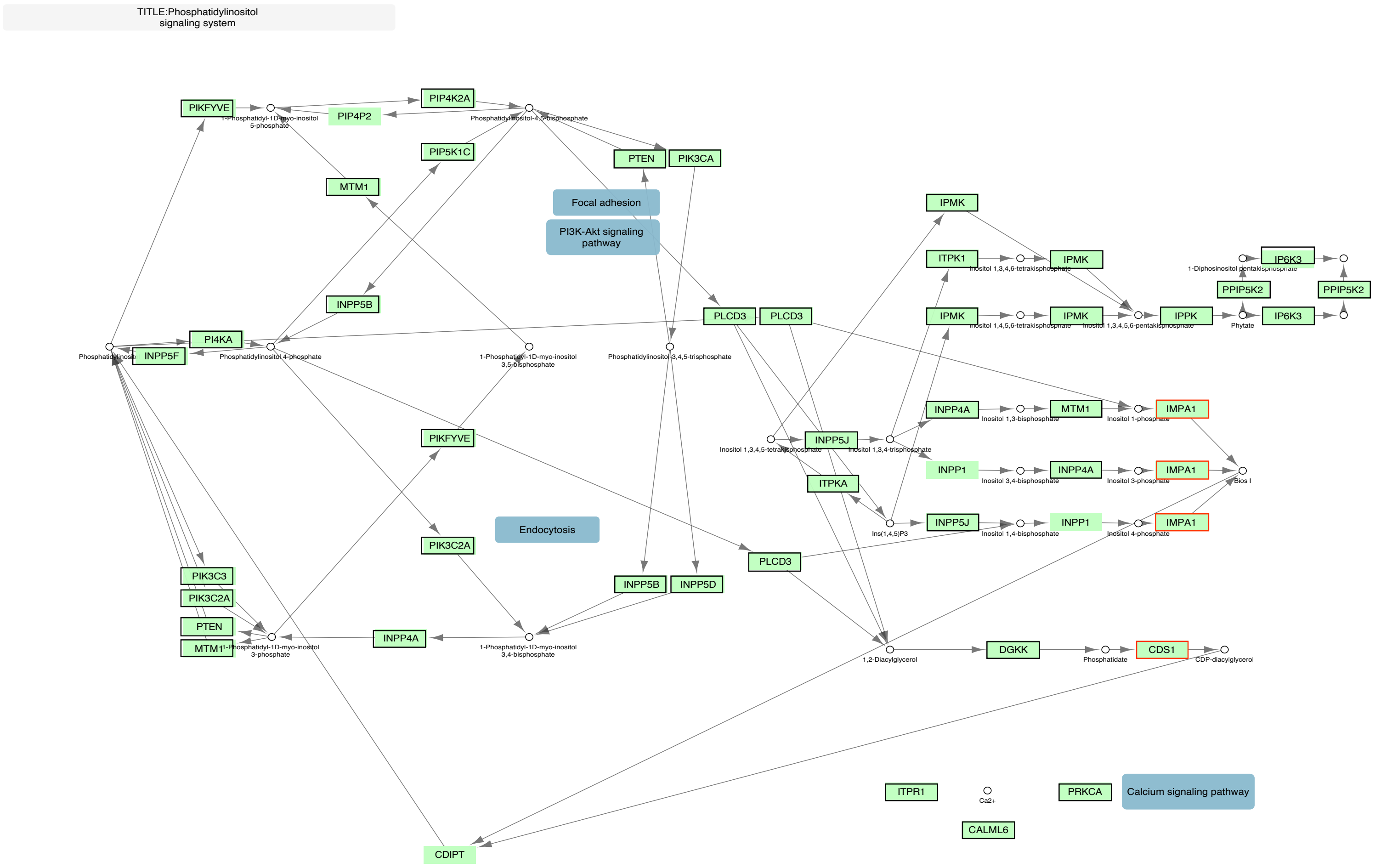

TITLE:Phosphatidylinositol signaling system
PIKFYVE
1-Phosphatidyl-1D-myo-inositol 5-phosphate
PIP4K2A
PIP4P2
Phosphatidylinositol-4,5-bisphosphate
PIP5K1C
PTEN
PIK3CA
MTM1
Focal adhesion
PI3K-Akt signaling pathway
INPP5B
PLCD3
PI4KA
INPP5F
Phosphatidylinositol 4-phosphate
1-Phosphatidyl-1D-myo-inositol 3,5-bisphosphate
Phosphatidylinositol-3,4,5-trisphosphate
PIKFYVE
Endocytosis
PIK3C2A
PIK3C3
PIK3C2A
PTEN
MTM1
1-Phosphatidyl-1D-myo-inositol 3-phosphate
INPP4A
1-Phosphatidyl-1D-myo-inositol 3,4-bisphosphate
INPP5B
INPP5D
IPMK
ITPK1
Inositol 1,3,4,6-tetrakisphosphate
IPMK
IP6K3
1-Diphosinositol pentakisphosphate
PPIP5K2
PPIP5K2
Inositol 1,4,5,6-tetrakisphosphate
Inositol 1,3,4,5,6-pentakisphosphate
IPPK
Phytate
IP6K3
INPP4A
Inositol 1,3-bisphosphate
MTM1
Inositol 1-phosphate
IMPA1
Inositol 1,3,4,5-tetrakisphosphate
INPP5J
Inositol 1,3,4-trisphosphate
INPP1
Inositol 3,4-bisphosphate
INPP4A
Inositol 3-phosphate
IMPA1
ITPKA
Ins(1,4,5)P3
INPP5J
Inositol 1,4-bisphosphate
INPP1
Inositol 4-phosphate
IMPA1
PLCD3
1,2-Diacylglycerol
DGKK
Phosphatidate
CDS1
CDP-diacylglycerol
ITPR1
Ca2+
PRKCA
Calcium signaling pathway
CALML6
CDIPT

E.

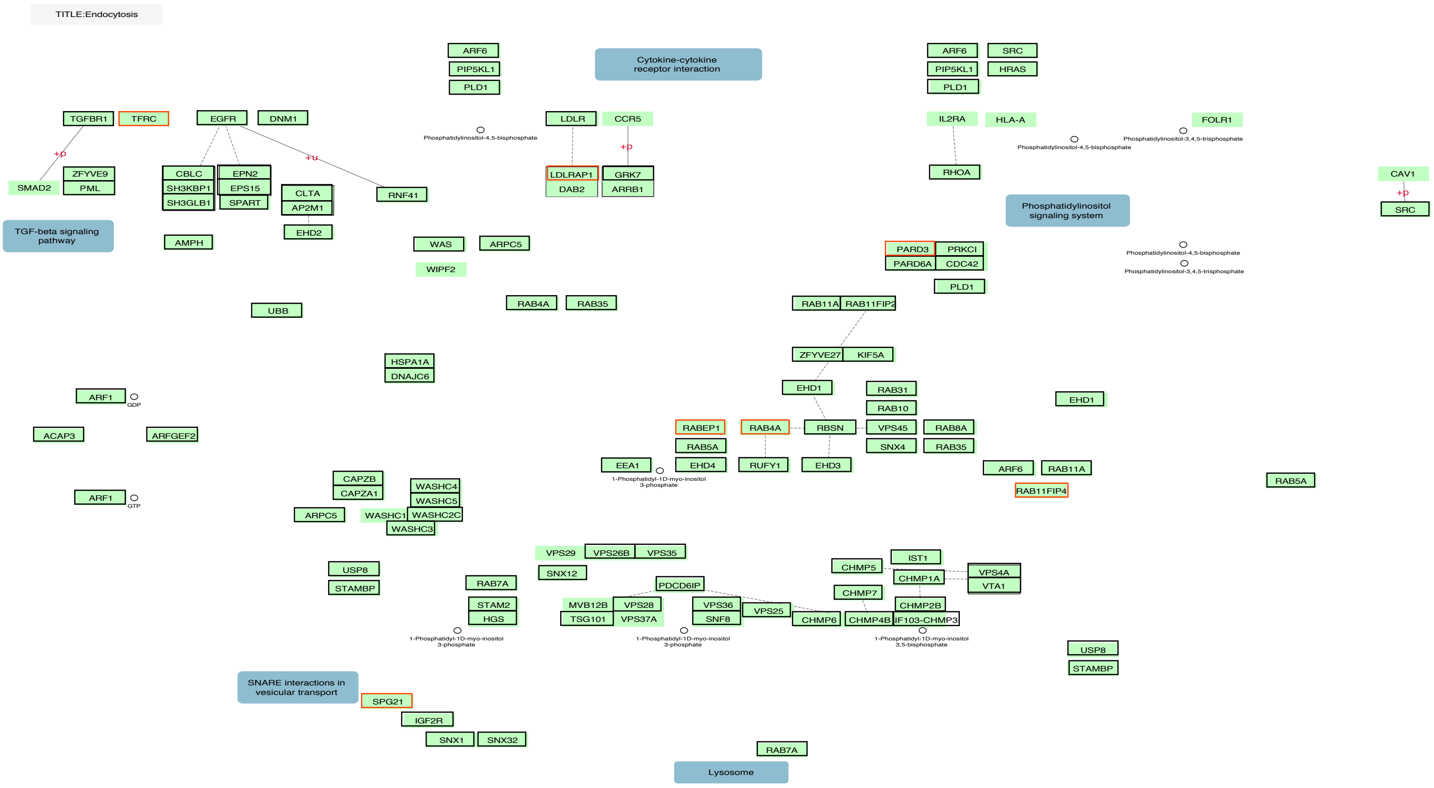
TITLE:Endocytosis
ARF6
PIP5KL1
PLD1
Cytokine-cytokine receptor interaction
ARF6
SRC
PIP5KL1
HRAS
PLD1
TGFBR1
TFRC
EGFR
DNM1
LDLR
CCR5
IL2RA
HLA-A
FOLR1
Phosphatidylinositol-4,5-bisphosphate
Phosphatidylinositol-4,5-bisphosphate
Phosphatidylinositol-3,4,5-trisphosphate
+p
+u
+p
SMAD2
ZFYVE9
PML
CBLC
SH3KBP1
SH3GLB1
EPN2
EPS15
SPART
CLTA
AP2M1
EHD2
RNF41
LDLRAP1
DAB2
GRK7
ARRB1
RHOA
CAV1
+p
SRC
Phosphatidylinositol signaling system
TGF-beta signaling pathway
AMPH
WAS
ARPC5
WIPF2
PARD3
PRKCI
PARD6A
CDC42
PLD1
Phosphatidylinositol-4,5-bisphosphate
Phosphatidylinositol-3,4,5-trisphosphate
UBB
RAB4A
RAB35
RAB11A
RAB11FIP2
HSPA1A
DNAJC6
ZFYVE27
KIF5A
ARF1
GDP
EHD1
RAB31
RAB10
EHD1
ACAP3
ARFGEF2
RABEP1
RAB4A
RBSN
VPS45
RAB8A
RAB5A
SNX4
RAB35
EEA1
EHD4
RUFY1
EHD3
1-Phosphatidyl-1D-myo-inositol 3-phosphate
ARF6
RAB11A
RAB11FIP4
RAB5A
ARF1
GTP
CAPZB
CAPZA1
WASHC4
WASHC5
ARPC5
WASHC1
WASHC2C
WASHC3
VPS29
VPS26B
VPS35
SNX12
USP8
STAMBP
RAB7A
PDCD6IP
CHMP5
IST1
CHMP1A
VPS4A
VTA1
CHMP7
STAM2
HGS
MVB12B
TSG101
VPS28
VPS37A
VPS36
SNF8
VPS25
CHMP6
CHMP4B
CHMP2B
CHMP3
1-Phosphatidyl-1D-myo-inositol 3-phosphate
1-Phosphatidyl-1D-myo-inositol 3-phosphate
1-Phosphatidyl-1D-myo-inositol 3,5-bisphosphate
USP8
STAMBP
SNARE interactions in vesicular transport
SPG21
IGF2R
SNX1
SNX32
RAB7A
Lysosome

F.

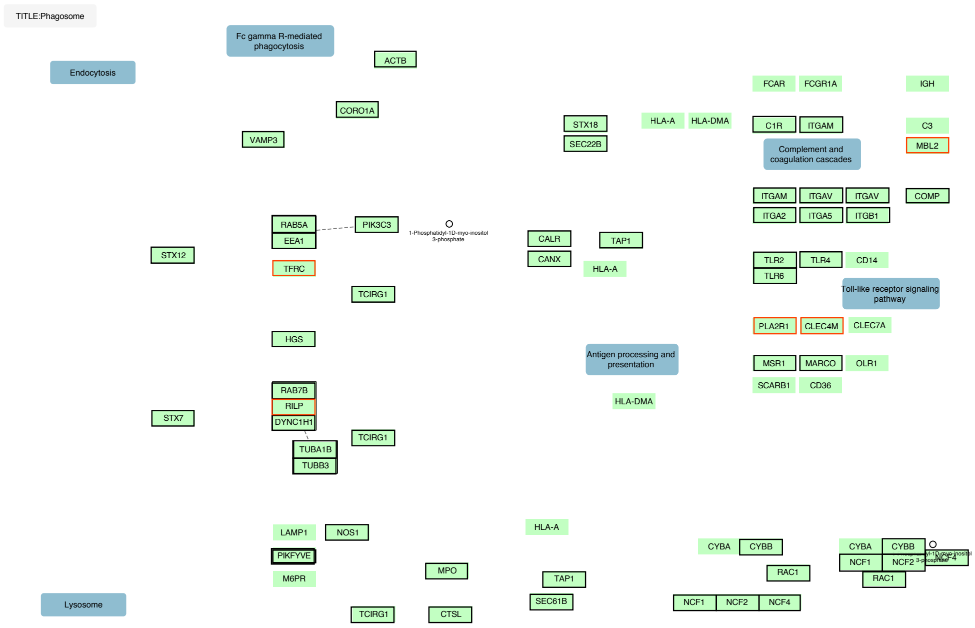
TITLE:Phagosome
Fc gamma R-mediated phagocytosis
Endocytosis
ACTB
CORO1A
VAMP3
STX18
SEC22B
HLA-A
HLA-DMA
FCAR
FCGR1A
IGH
C1R
ITGAM
C3
MBL2
Complement and coagulation cascades
ITGAM
ITGAV
ITGAV
COMP
ITGA2
ITGA5
ITGB1
RAB5A
EEA1
PIK3C3
1-Phosphatidyl-1D-myo-inositol 3-phosphate
CALR
CANX
TAP1
HLA-A
STX12
TFRC
TCIRG1
TLR2
TLR4
CD14
TLR6
Toll-like receptor signaling pathway
HGS
PLA2R1
CLEC4M
CLEC7A
Antigen processing and presentation
MSR1
MARCO
OLR1
SCARB1
CD36
RAB7B
RILP
DYNC1H1
HLA-DMA
STX7
TCIRG1
TUBA1B
TUBB3
HLA-A
LAMP1
NOS1
PIKFYVE
MPO
M6PR
TAP1
SEC61B
CYBA
CYBB
RAC1
CYBA
CYBB
NCF1
NCF2
NCF4
RAC1
NCF1
NCF2
NCF4
Lysosome
TCIRG1
CTSL

Figure S4. Annotated KEGG pathways for Membranes and cytosis. Human pathways for membranes, endocytosis and phagosome: A. Glycerolipid metabolism (hsa00561), B. Glycerophospholipid metabolism (hsa00564), C. Inositol phosphate metabolism (hsa00562), D. Phosphatidylinositol signaling system (hsa04070), E. Endocytosis (hsa04144), and F. Phagosome (hsa04145). Genes in the pathways that are found in the monophyletic clusters are highlighted with a black outline and complex history clusters are highlighted with a red outline.
