## Supplemental Figure S5 for "Reconstructing and Analysing The Genome of The Last Eukaryote Common Ancestor to Better Understand the Transition from FECA to LECA"

A.

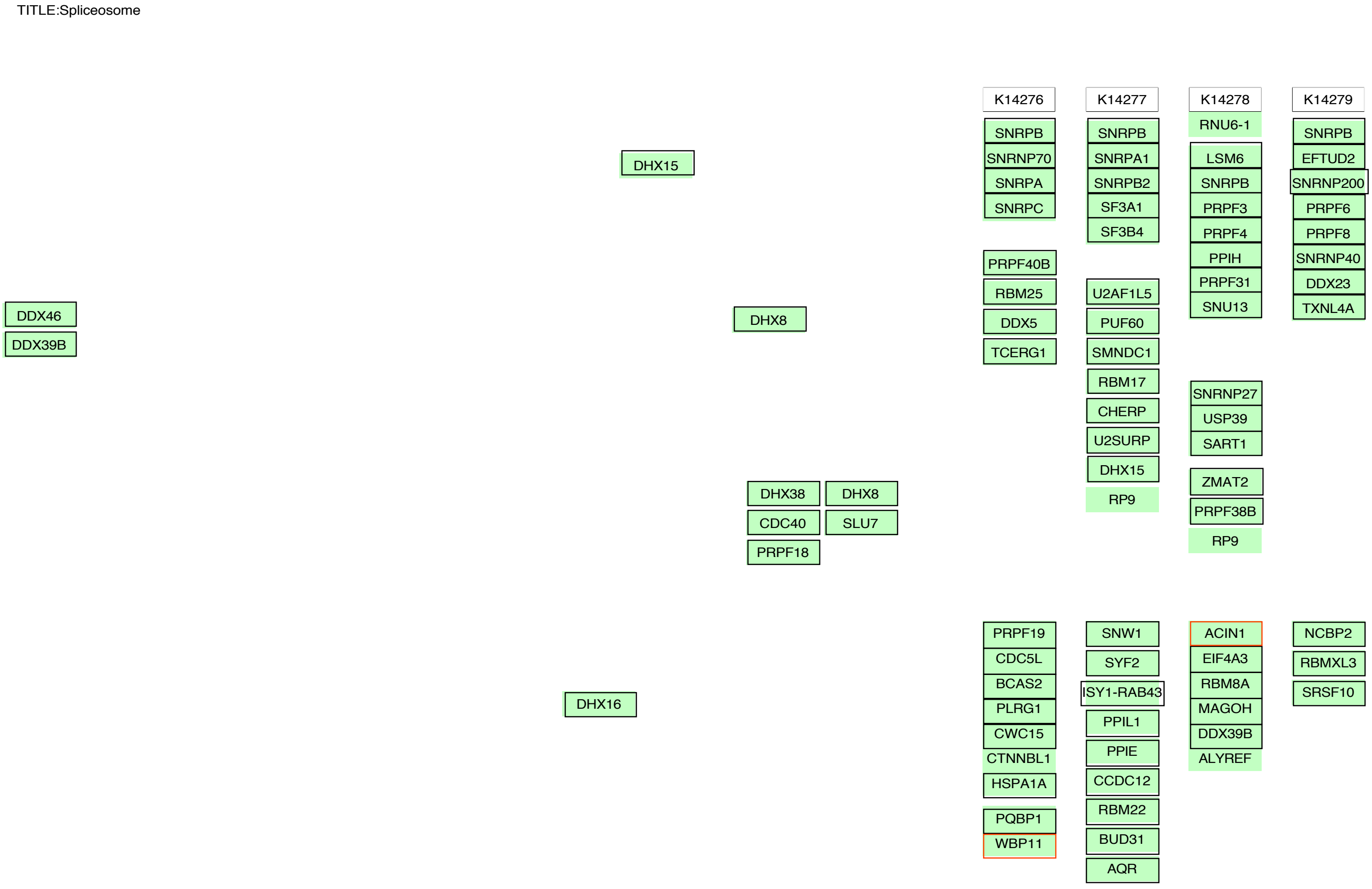
TITLE:Spliceosome
DHX15
DDX46
DDX39B
DHX8
DHX38
CDC40
PRPF18
DHX8
SLU7
DHX16
K14276
SNRPB
SNRNP70
SNRPA
SNRPC
PRPF40B
RBM25
DDX5
TCERG1
PRPF19
CDC5L
BCAS2
PLRG1
CWC15
CTNNBL1
HSPA1A
PQBP1
WBP11
K14277
SNRPB
SNRPA1
SNRPB2
SF3A1
SF3B4
U2AF1L5
PUF60
SMNDC1
RBM17
CHERP
U2SURP
DHX15
RP9
SNW1
SYF2
ISY1-RAB43
PPIL1
PPIE
CCDC12
RBM22
BUD31
AQR
K14278
RNU6-1
LSM6
SNRPB
PRPF3
PRPF4
PPIH
PRPF31
SNU13
SNRNP27
USP39
SART1
ZMAT2
PRPF38B
RP9
ACIN1
EIF4A3
RBM8A
MAGOH
DDX39B
ALYREF
K14279
SNRPB
EFTUD2
SNRNP200
PRPF6
PRPF8
SNRNP40
DDX23
TXNL4A
NCBP2
RBMXL3
SRSF10

B.

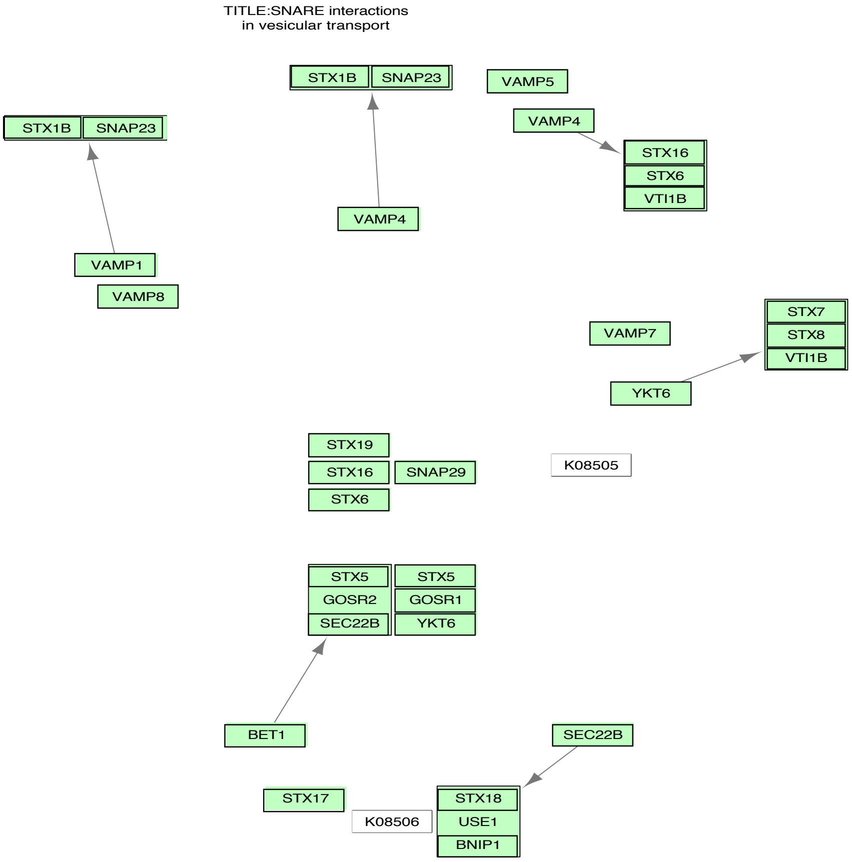
TITLE:SNARE interactions in vesicular transport
STX1B
SNAP23
STX1B
SNAP23
VAMP5
VAMP4
STX16
STX6
VTI1B
VAMP4
VAMP1
VAMP8
VAMP7
STX7
STX8
VTI1B
YKT6
STX19
STX16
SNAP29
STX6
K08505
STX5
GOSR2
SEC22B
STX5
GOSR1
YKT6
BET1
SEC22B
STX17
K08506
STX18
USE1
BNIP1

C.

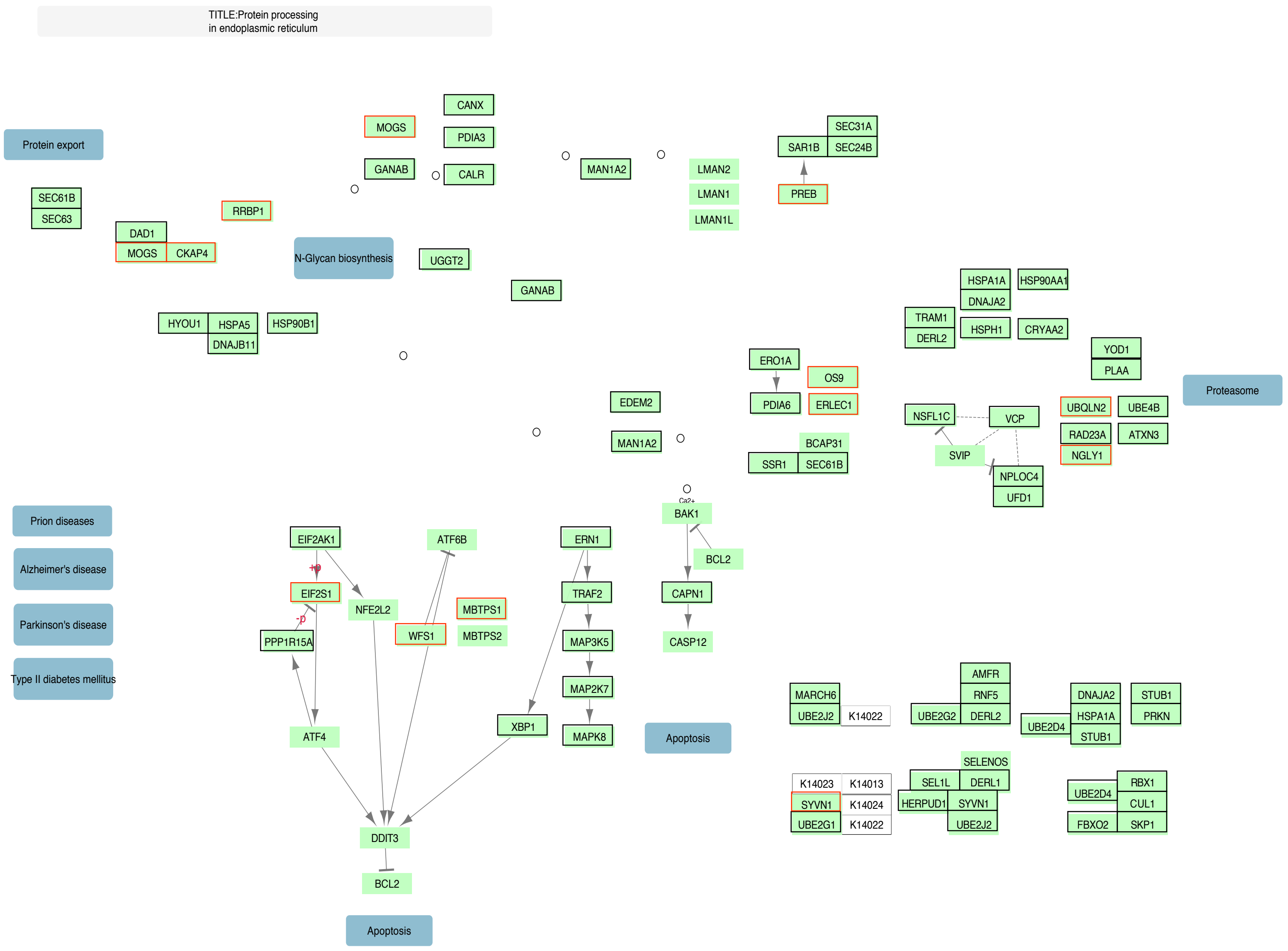
TITLE:Protein processing in endoplasmic reticulum
Protein export
SEC61B
SEC63
DAD1
MOGS
CKAP4
RRBP1
MOGS
GANAB
CANX
PDIA3
CALR
MAN1A2
LMAN2
LMAN1
LMAN1L
SEC31A
SAR1B
SEC24B
PREB
N-Glycan biosynthesis
UGGT2
GANAB
HYOU1
HSPA5
DNAJB11
HSP90B1
HSPA1A
HSP90AA1
DNAJA2
TRAM1
DERL2
HSPH1
CRYAA2
YOD1
PLAA
Proteasome
ERO1A
PDIA6
OS9
ERLEC1
EDEM2
MAN1A2
BCAP31
SSR1
SEC61B
NSFL1C
VCP
SVIP
NPLOC4
UFD1
UBQLN2
UBE4B
RAD23A
ATXN3
NGLY1
Prion diseases
Alzheimer's disease
Parkinson's disease
Type II diabetes mellitus
EIF2AK1
+p
EIF2S1
-p
PPP1R15A
NFE2L2
ATF6B
WFS1
MBTPS1
MBTPS2
ERN1
TRAF2
MAP3K5
MAP2K7
MAPK8
XBP1
Ca2+
BAK1
BCL2
CAPN1
CASP12
Apoptosis
ATF4
DDIT3
BCL2
Apoptosis
MARCH6
UBE2J2
K14022
AMFR
RNF5
UBE2G2
DERL2
DNAJA2
HSPA1A
STUB1
STUB1
PRKN
UBE2D4
SELENOS
K14023
K14013
SEL1L
DERL1
SYVN1
K14024
HERPUD1
SYVN1
UBE2G1
K14022
UBE2J2
RBX1
UBE2D4
CUL1
FBXO2
SKP1

D.

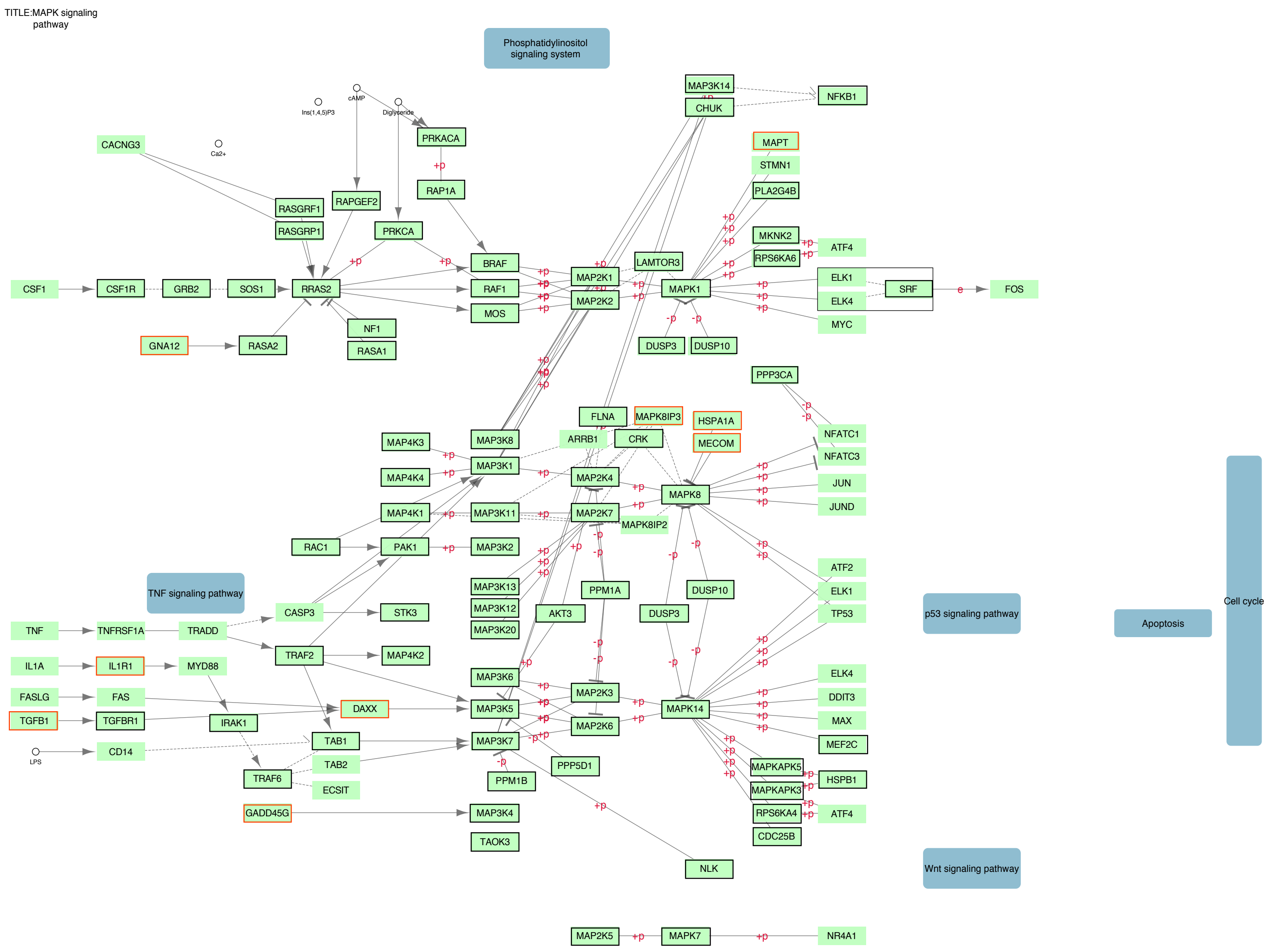
TITLE:MAPK signaling pathway
Phosphatidylinositol signaling system
TNF signaling pathway
p53 signaling pathway
Apoptosis
Cell cycle
Wnt signaling pathway
CACNG3
Ca2+
Ins(1,4,5)P3
cAMP
Diglyceride
PRKACA
RAP1A
RASGRF1
RASGRP1
RAPGEF2
PRKCA
CSF1
CSF1R
GRB2
SOS1
RRAS2
BRAF
RAF1
MOS
NF1
RASA1
GNA12
RASA2
MAP2K1
MAP2K2
LAMTOR3
MAPK1
MAP3K14
CHUK
NFKB1
MAPT
STMN1
PLA2G4B
MKNK2
RPS6KA6
ATF4
ELK1
ELK4
SRF
FOS
MYC
DUSP3
DUSP10
PPP3CA
NFATC1
NFATC3
JUN
JUND
FLNA
MAPK8IP3
HSPA1A
MECOM
ARRB1
CRK
MAP4K3
MAP4K4
MAP4K1
MAP3K8
MAP3K1
MAP3K11
MAP2K4
MAP2K7
MAPK8
MAPK8IP2
RAC1
PAK1
MAP3K2
MAP3K13
MAP3K12
MAP3K20
AKT3
PPM1A
ATF2
TP53
TNF
TNFRSF1A
TRADD
CASP3
STK3
IL1A
IL1R1
MYD88
TRAF2
MAP4K2
FASLG
FAS
DAXX
MAP3K6
MAP3K5
MAP2K3
MAP2K6
MAPK14
DDIT3
MAX
MEF2C
TGFB1
TGFBR1
IRAK1
LPS
CD14
TAB1
TAB2
TRAF6
ECSIT
MAP3K7
PPM1B
PPP5D1
MAPKAPK5
MAPKAPK3
HSPB1
RPS6KA4
CDC25B
GADD45G
MAP3K4
TAOK3
NLK
MAP2K5
MAPK7
NR4A1
+p
-p
e

Figure S5. Annotated KEGG pathways for Gene Expression, Protein Production, and MAPK signaling. Human pathways for gene expression, protein production and MAPK signaling: A. Spliceosome (hsa03040), B. SNARE interactions in vesicular transport (hsa04130), C. Protein processing in endoplasmic reticulum (hsa04141), and D. MAPK signaling pathway (hsa04010). Genes in the pathways that are found in the monophyletic clusters are highlighted with a black outline and complex history clusters are highlighted with a red outline.
