## Supplemental Figure S6 for "Reconstructing and Analysing The Genome of The Last Eukaryote Common Ancestor to Better Understand the Transition from FECA to LECA"

A.

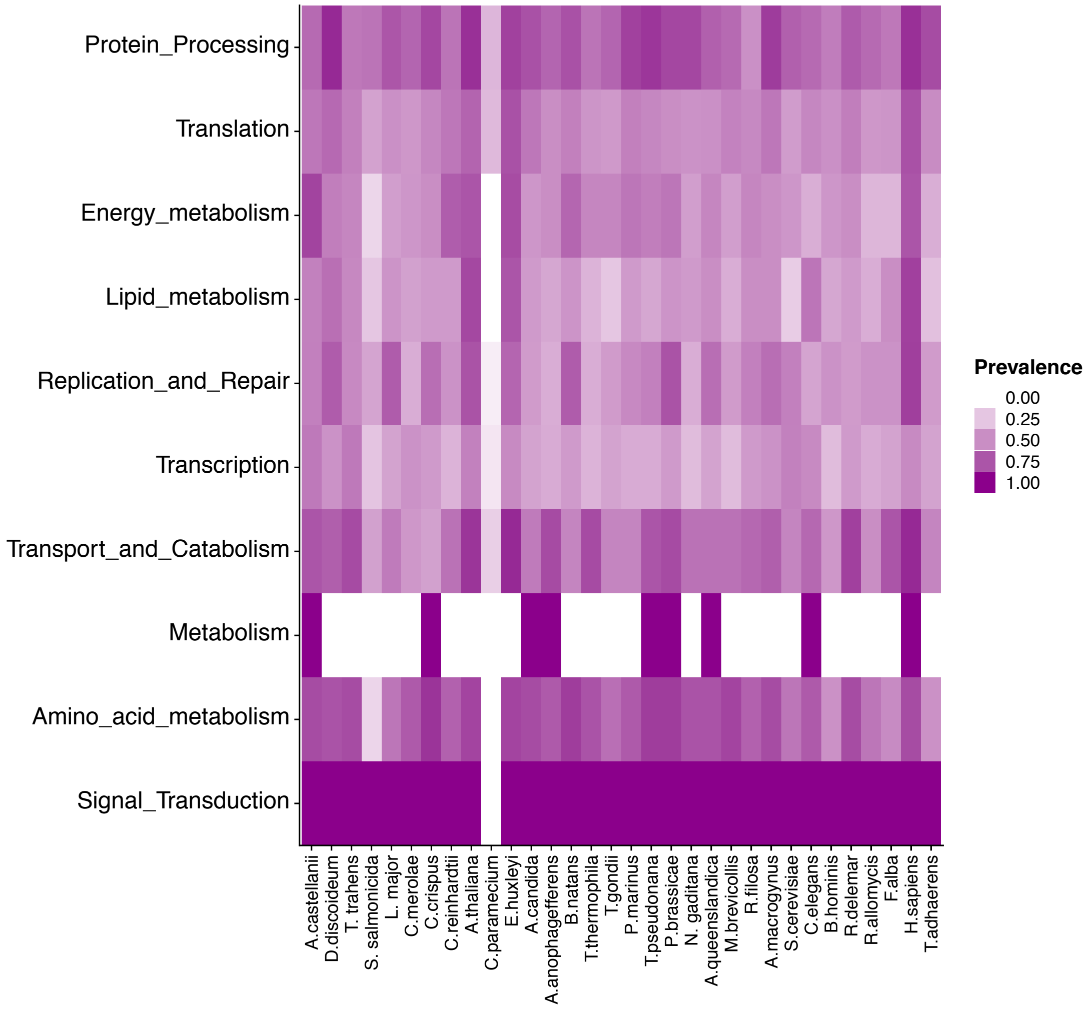
Protein_Processing
Translation
Energy_metabolism
Lipid_metabolism
Replication_and_Repair
Transcription
Transport_and_Catabolism
Metabolism
Amino_acid_metabolism
Signal_Transduction
A.castellanii
D.discoideum
T. trahens
S. salmonicida
L. major
C.merolae
C.crispus
C.reinhardtii
A.thaliana
C.paramecium
E.huxleyi
A.candida
A.anophagefferens
B.natans
T.thermophila
T.gondii
P.marinus
T.pseudonana
P.brassicae
N. gaditana
A.queenslandica
M.brevicollis
R.filosa
A.macrogynus
S.cerevisiae
C.elegans
B.hominis
R.delemar
R.allomycis
F.alba
H.sapiens
T.adhaerens
Prevalence
0.00
0.25
0.50
0.75
1.00

B.

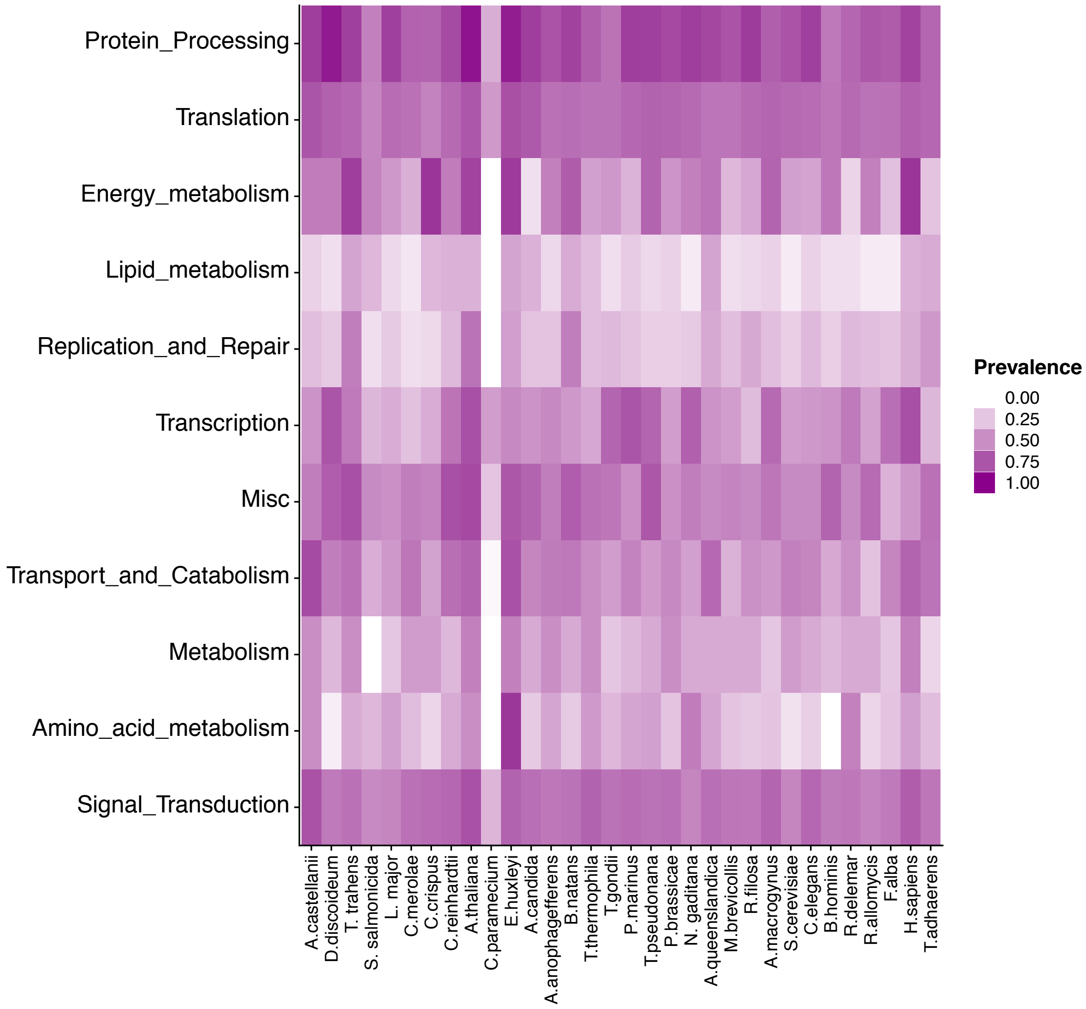
Protein_Processing
Translation
Energy_metabolism
Lipid_metabolism
Replication_and_Repair
Transcription
Misc
Transport_and_Catabolism
Metabolism
Amino_acid_metabolism
Signal_Transduction
A.castellanii
D.discoideum
T. trahens
S. salmonicida
L. major
C.merolae
C.crispus
C.reinhardtii
A.thaliana
C.paramecium
E.huxleyi
A.candida
A.anophagefferens
B.natans
T.thermophila
T.gondii
P.marinus
T.pseudonana
P.brassicae
N. gaditana
A.queenslandica
M.brevicollis
R.filosa
A.macrogynus
S.cerevisiae
C.elegans
B.hominis
R.delemar
R.allomycis
F.alba
H.sapiens
T.adhaerens
Prevalence
0.00
0.25
0.50
0.75
1.00

C.

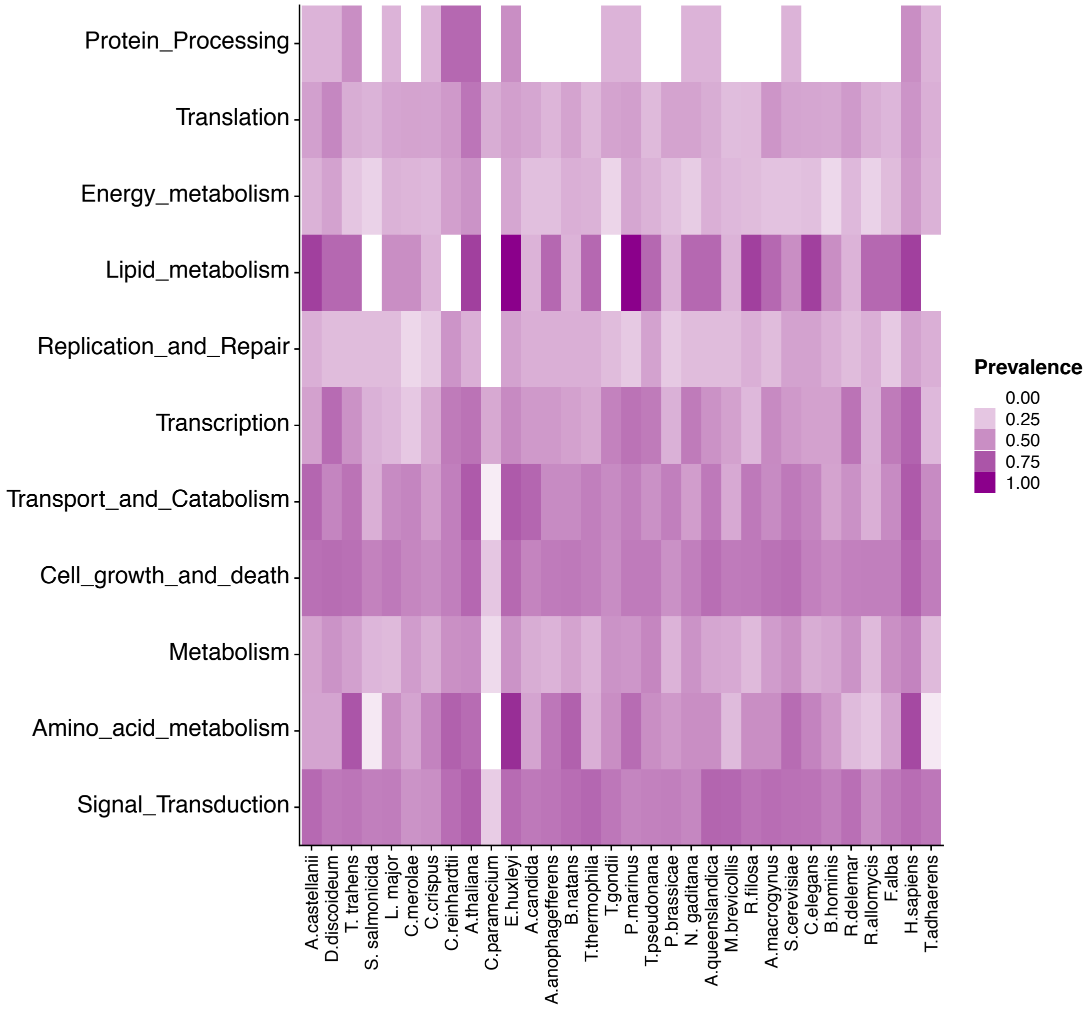
Protein_Processing
Translation
Energy_metabolism
Lipid_metabolism
Replication_and_Repair
Transcription
Transport_and_Catabolism
Cell_growth_and_death
Metabolism
Amino_acid_metabolism
Signal_Transduction
A.castellanii
D.discoideum
T. trahens
S. salmonicida
L. major
C.merolae
C.crispus
C.reinhardtii
A.thaliana
C.paramecium
E.huxleyi
A.candida
A.anophagefferens
B.natans
T.thermophila
T.gondii
P.marinus
T.pseudonana
P.brassicae
N. gaditana
A.queenslandica
M.brevicollis
R.filosa
A.macrogynus
S.cerevisiae
C.elegans
B.hominis
R.delemar
R.allomycis
F.alba
H.sapiens
T.adhaerens
Prevalence
0.00
0.25
0.50
0.75
1.00

**D.**

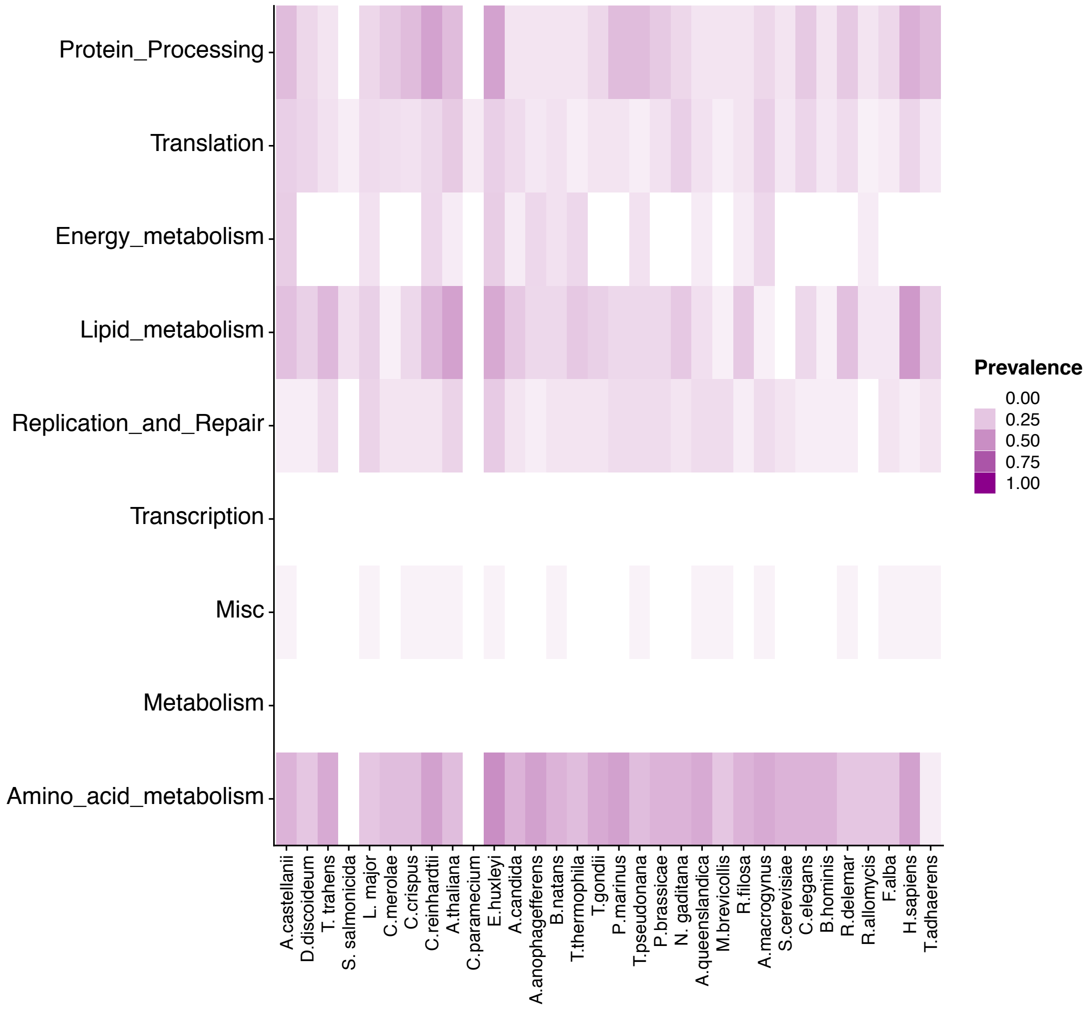
Protein_Processing
Translation
Energy_metabolism
Lipid_metabolism
Replication_and_Repair
Transcription
Misc
Metabolism
Amino_acid_metabolism
A.castellanii
D.discoideum
T. trahens
S. salmonicida
L. major
C.merolae
C.crispus
C.reinhardtii
A.thaliana
C.paramecium
E.huxleyi
A.candida
A.anophagefferens
B.natans
T.thermophila
T.gondii
P.marinus
T.pseudonana
P.brassicae
N. gaditana
A.queenslandica
M.brevicollis
R.filosa
A.macrogynus
S.cerevisiae
C.elegans
B.hominis
R.delemar
R.allomycis
F.alba
H.sapiens
T.adhaerens
Prevalence
0.00
0.25
0.50
0.75
1.00

.

**E.**

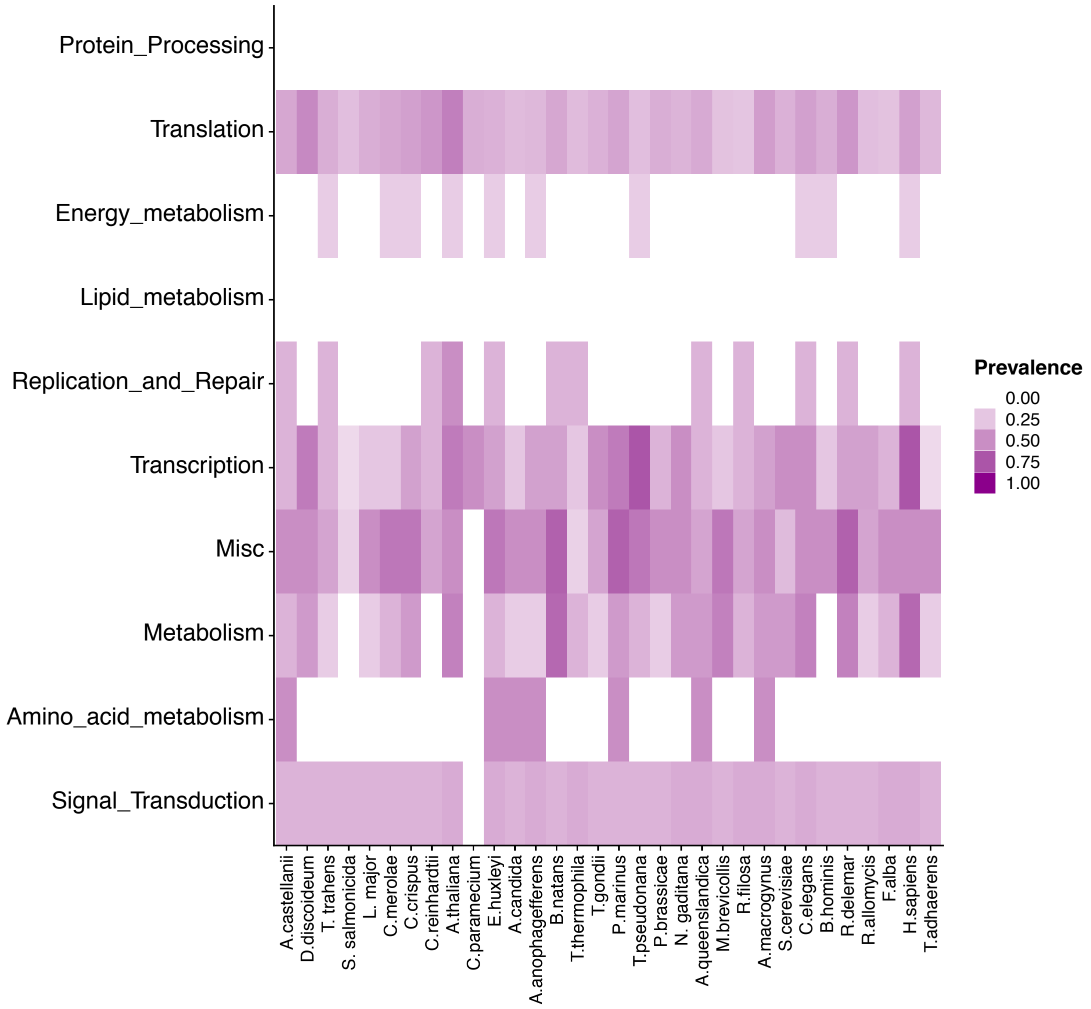


**Figure S6**. Heatmaps showing pathway completeness for (**A**) amoeba (*Dictyostelium discoideum*), (**B**) plant (Arabidopsis thaliana), (**C**) yeast (*Saccharomyces cerevisiae*), (**D**) bacterial (*Escherichia coli*) and (**E**) archaeal (*Methanobrevibacter smithii*) pathways across each of the constituent genomes of the reconstructed LECA genome. KEGG pathways categorised into groups of similar or linked function. *C. paramecium* represents a minimalist nucleomorph genome that lacks it's own metabolic genes.
