## Supplemental Figure S7 for "Reconstructing and Analysing The Genome of The Last Eukaryote Common Ancestor to Better Understand the Transition from FECA to LECA"

**A.**

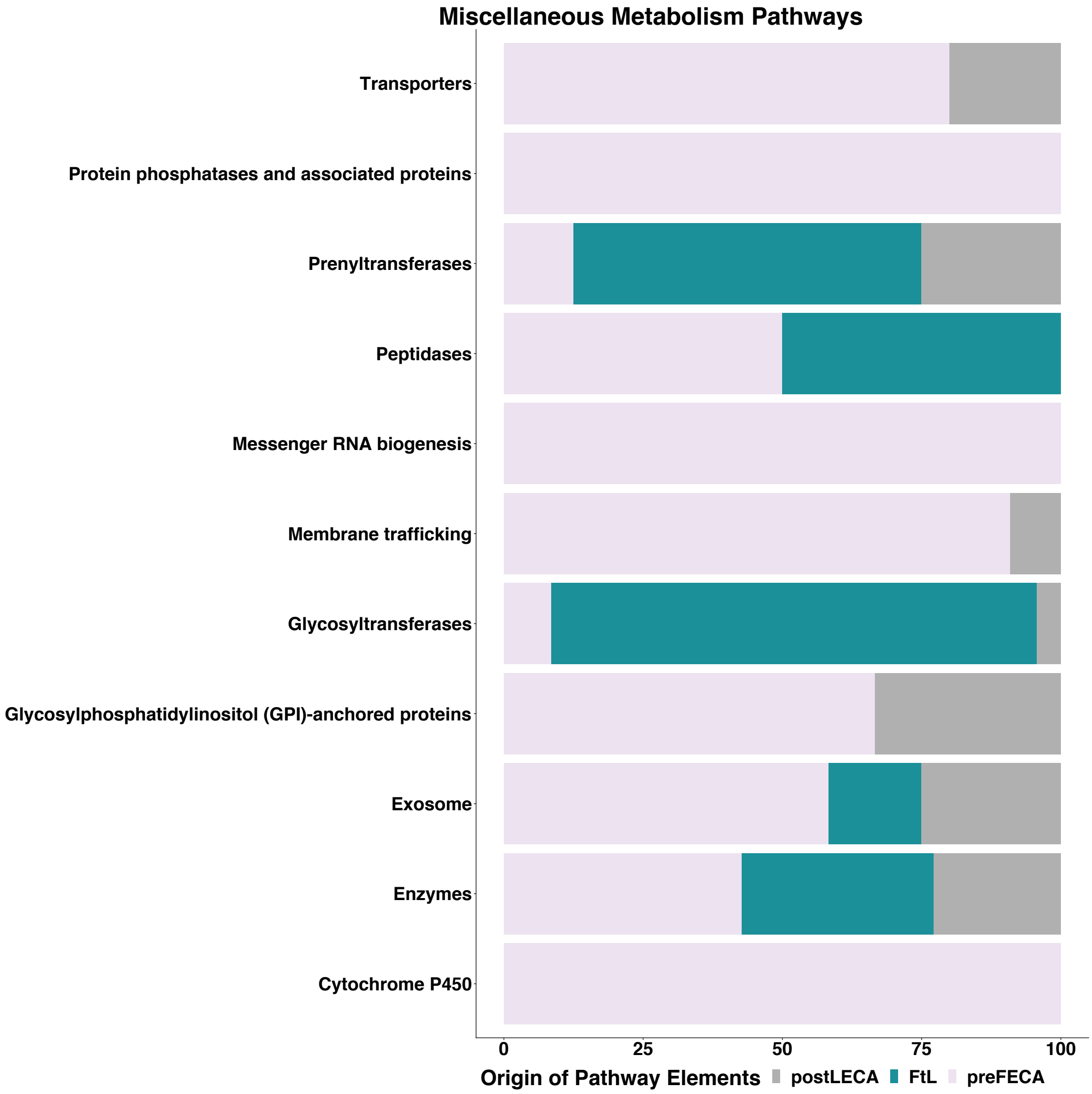
Miscellaneous Metabolism Pathways
Transporters
Protein phosphatases and associated proteins
Prenyltransferases
Peptidases
Messenger RNA biogenesis
Membrane trafficking
Glycosyltransferases
Glycosylphosphatidylinositol (GPI)-anchored proteins
Exosome
Enzymes
Cytochrome P450
0
25
50
75
100
Origin of Pathway Elements
postLECA
FtL
preFECA

**B.**

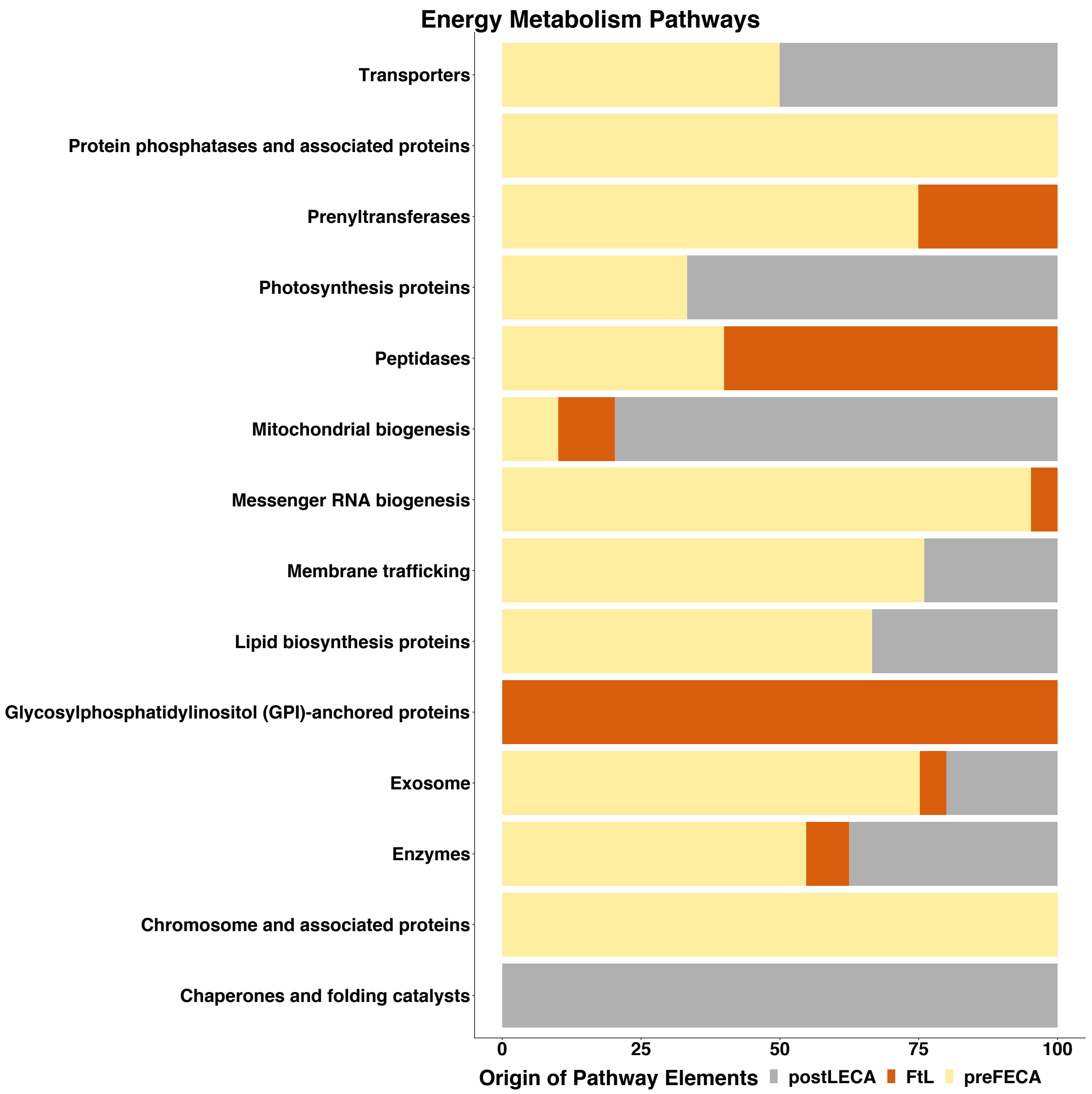
Energy Metabolism Pathways
Transporters
Protein phosphatases and associated proteins
Prenyltransferases
Photosynthesis proteins
Peptidases
Mitochondrial biogenesis
Messenger RNA biogenesis
Membrane trafficking
Lipid biosynthesis proteins
Glycosylphosphatidylinositol (GPI)-anchored proteins
Exosome
Enzymes
Chromosome and associated proteins
Chaperones and folding catalysts
0
25
50
75
100
Origin of Pathway Elements
postLECA
FtL
preFECA

C.

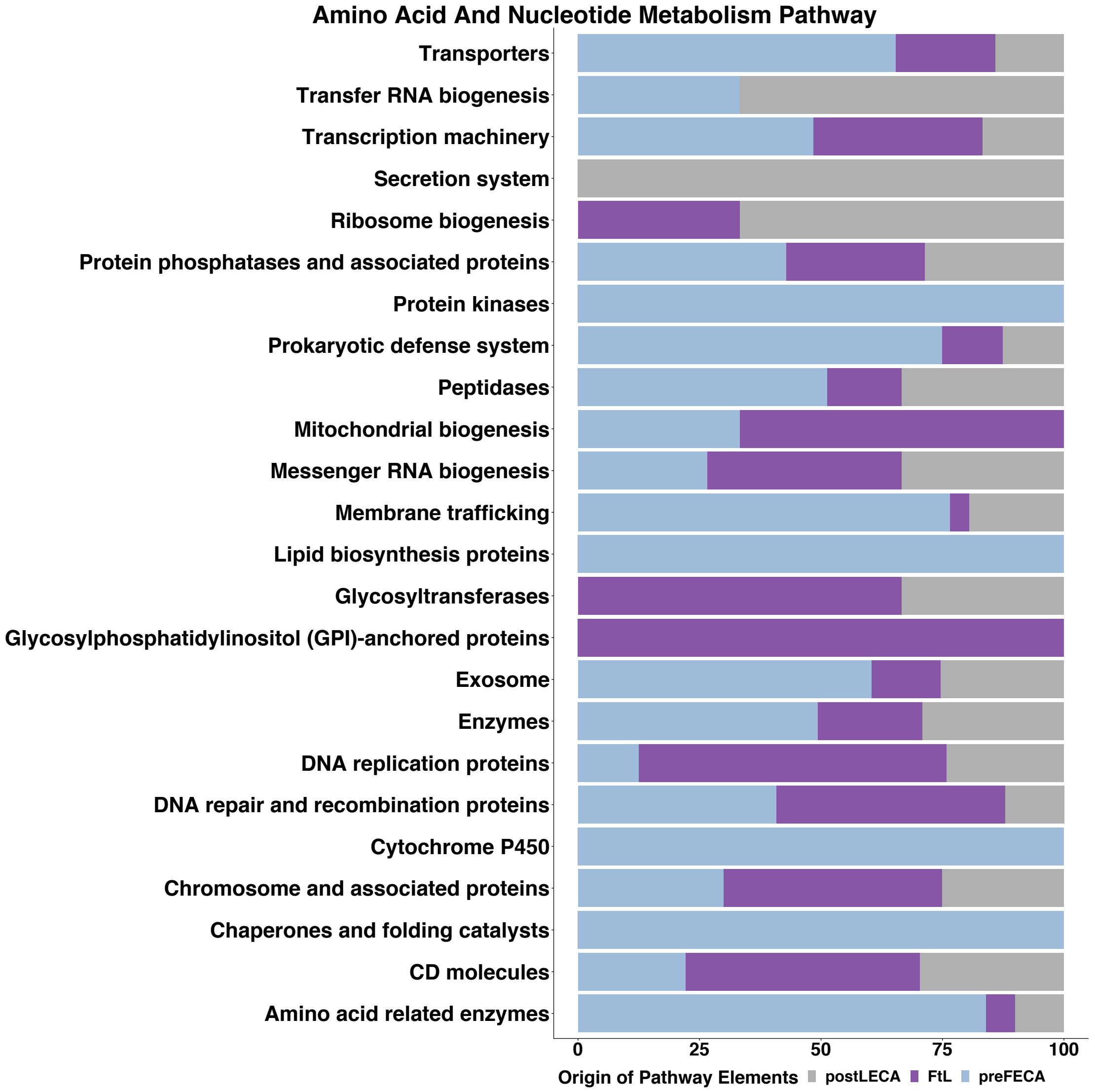


**Figure S7**. Proportion of BRITE functional classifications for genes, which originate pre-FECA, during the FECA to LECA transition, or post-LECA, from (A) Miscellaneous metabolism, (**B**) Energy metabolism, and (**C**) amino acid and nucleotide metabolism pathways that saw gene gain during the FECA to LECA transition. FtL = FECA-to-LECA transition.
