## Supplemental Table S1 for "Reconstructing and Analysing The Genome of The Last Eukaryote Common Ancestor to Better Understand the Transition from FECA to LECA"

| Eukaryote Genomes |
| --- |
| Acanthamoeba_castellanii_str_neff.Acastellanii_strNEFF_v1.pep.all.fa |
| Albugo_candida.ASM107853v1.pep.all.fa |
| Allomyces_macrogynus_atcc_38327.A_macrogynus_V3.pep.all.fa |
| Amphimedon_queenslandica.Aqu1.pep.all.fa |
| Arabidopsis_thaliana.TAIR10.pep.all.fa |
| Aureococcus_anophagefferens.v_1_0.pep.all.fa |
| Bigelowiella_natans.GCA_000320545.1.pep.all.fa |
| Blastocystis_hominis.ASM15166v1.pep.all.fa |
| Caenorhabditis_elegans.WBcel235.pep.all.fa |
| Chlamydomonas_reinhardtii.v3.1.pep.all.fa |
| Chondrus_crispus.ASM35022v2.pep.all.fa |
| Cryptomonas_paramecium.ASM19445v1.pep.all.fa |
| Cyanidioschyzon_merolae.ASM9120v1.pep.all.fa |
| Dictyostelium_discoideum.dicty_2.7.pep.all.fa |
| Emiliania_huxleyi.GCA_000372725.1.pep.all.fa |
| Fonticula_alba.Font_alba_ATCC_38817_V2.pep.all.fa |
| Homo_sapiens.GRCh38.pep.all.fa |
| Leishmania_major.ASM272v2.pep.all.fa |
| Monosiga_brevicollis_mx1.V1.0.pep.all.fa |
| Nannochloropsis_gaditana.NagaB31_1.0.pep.all.fa |
| Perkinsus_marinus_atcc_50983.JCVI_PMG_1.0.pep.all.fa |
| Plasmodiophora_brassicae.pbe3.h15.pep.all.fa |
| Reticulomyxa_filosa.Reti_assembly1.0.pep.all.fa |
| Rhizopus_delemar_ra_99_880.RO3.pep.all.fa |
| Rozella_allomycis_csf55.Rozella_k41_t100.pep.all.fa |
| Saccharomyces_cerevisiae.R64-1-1.pep.all.fa |
| Spironucleus_salmonicida.SSK3.0.pep.all.fa |
| Tetrahymena_thermophila.JCVI-TTA1-2.2.pep.all.fa |
| Thalassiosira_pseudonana.ASM14940v2.pep.all.fa |
| Thecamonas_trahens_atcc_50062.TheTra_May2010.pep.all.fa |
| Toxoplasma_gondii.ToxoDB-7.1.pep.all.fa |
| Trichoplax_adhaerens.ASM15027v1.pep.all.fa |
| Prokaryote Genomes |
| Acaryochloris_marina_mbic11017.ASM1810v1.pep.all.fa |
| Acetobacter_tropicalis.ASM75566v1.pep.all.fa |
| Acholeplasma_laidlawii_pg_8a.ASM1878v1.pep.all.fa |
| Acidianus_hospitalis_w1.ASM21321v1.pep.all.fa |
| Acidilobus_saccharovorans_345_15.ASM14491v1.pep.all.fa |
| Acidiphilium_cryptum_jf_5.ASM1672v1.pep.all.fa |
| Acidiplasma_aeolicum.ASM140294v1.pep.all.fa |
| Acidiplasma_cupricumulans.ASM140293v1.pep.all.fa |
| Acidithrix_ferrooxidans.ASM94929v1.pep.all.fa |
| Acidobacterium_capsulatum_atcc_51196.ASM2256v1.pep.all.fa |
| Acidovorax_temperans.ASM93558v1.pep.all.fa |
| Aciduliprofundum_boonei.T469.439481.9.PATRIC.fa |
| Aenigmarchaeota.archaeon.JGI0000106-F11.1130284.4.PATRIC.fa |
| Aeropyrum_pernix_k1.ASM1112v1.pep.all.fa |
| Agathobacter_rectalis_dsm_17629.ASM20993v1.pep.all.fa |
| Aigarchaeota_archaeon.JGI000010J15.1130285.3.PATRIC.fa |

| |
|---|
| Akkermansia_muciniphila_atcc_baa_835.ASM2022v1.pep.all.fa |
| Alcaligenes_faecalis.ASM77001v1.pep.all.fa |
| Algibacter_lectus.ASM76483v1.pep.all.fa |
| Algoriphagus_machipongonensis.ASM16627v1.pep.all.fa |
| Aminobacterium_colombiense.DSM12261.572547.3.PATRIC.fa |
| Anaerolinea_thermophila_uni_1.ASM19967v1.pep.all.fa |
| Aquifex_aeolicus.VF5.224324.8.PATRIC.fa |
| Archaeoglobus_fulgidus.2234.7.PATRIC.fa |
| Bathyarchaeota.1700835.3.PATRIC.fa |
| Borrelia_afzelii.HLJ01.1239934.3.PATRIC.fa |
| Bryobacter_aggregatus.MPL3.1340493.3.PATRIC.fa |
| Caldisericum_exile.AZM16c01.511051.3.PATRIC.fa |
| Calditerrivibrio_nitroreducens.DSM19672.768670.3.PATRIC.fa |
| Caldithrix_abyssi_dsm_13497.ASM24181v1.pep.all.fa |
| Campylobacter_jejuni.CJJ5070.pep.all.fa |
| Candidate_division_ws6_bacterium_gw2011_gwa2_37_6.ASM98950v1.pep.all.fa |
| Candidatus.Parvarchaeota.994838.4.PATRIC.fa |
| Candidatus_Aenigmarchaeota743730.4.PATRIC.fa |
| Candidatus_Caldiarchaeum_subterraneum.311458.9.PATRIC.fa |
| Candidatus_Koribacter_versatilis.Ellin345.204669.11.PATRIC.fa |
| Candidatus_atelocyanobacterium_thalassa_isolate_aloha.ASM2512v1.pep.all.fa |
| Candidatus_azambacteria_bacterium_gw2011_gwa2_45_90.ASM100212v1.pep.all.fa |
| Candidatus_beckwithbacteria_bacterium_gw2011_gwa2_47_25.ASM100188v1.pep.all.fa |
| Candidatus_campbellbacteria_bacterium_gw2011_gwd1_35_49.ASM99086v1.pep.all.fa |
| Candidatus_curtissbacteria_bacterium_gw2011_gwa1_40_16.ASM99599v1.pep.all.fa |
| Candidatus_daviesbacteria_bacterium_gw2011_gwc2_40_12.ASM99590v1.pep.all.fa |
| Candidatus_entotheonella_sp_tsy1.v3.pep.all.fa |
| Candidatus_korarchaeum_cryptofilum_opf8.ASM1960v1.pep.all.fa |
| Candidatus_magasanikbacteria_bacterium_gw2011_gwa2_45_39.ASM100165v1.pep.all.fa |
| Candidatus_nitrosoarchaeum_koreensis_my1.ASM22017v1.pep.all.fa |
| Candidatus_nitrososphaera_evergladensis_sr1.ASM73028v1.pep.all.fa |
| Candidatus_nomurabacteria_bacterium_gw2011_gwa1_46_11.ASM100221v1.pep.all.fa |
| Candidatus_omnitrophus_magneticus.ASM95409v1.pep.all.fa |
| Candidatus_peregrinibacteria_bacterium_gw2011_gwf2_43_17.ASM99931v1.pep.all.fa |
| Candidatus_saccharimonas_aalborgensis.ASM39243v1.pep.all.fa |
| Candidatus_uhrbacteria_bacterium_gw2011_gwa2_53_10.ASM100456v1.pep.all.fa |
| Candidatus_woesebacteria_bacterium_gw2011_gwa1_41_13b.ASM99654v1.pep.all.fa |
| Candidatus_wolfebacteria_bacterium_gw2011_gwb1_47_243.ASM100030v1.pep.all.fa |
| Candidatus_yanofskybacteria_bacterium_gw2011_gwc2_41_9.ASM99770v1.pep.all.fa |
| Chlamydophila_pneumoniae_ar39.ASM9108v1.pep.all.fa |
| Chroococcidiopsis_thermalis_pcc_7203.ASM31712v1.pep.all.fa |
| Chthonomonas_calidirosea.WRG1.2.pep.all.fa |
| Corynebacterium_argentoratense_dsm_44202.ASM59055v1.pep.all.fa |
| Croceibacter_atlanticus.HTCC2559.216432.7.PATRIC.fa |
| Cytophaga_hutchinsonii.ATCC33406.269798.16.PATRIC.fa |
| Deinococcus_geothermalis_dsm_11300.ASM19627v1.pep.all.fa |
| Desulfurispirillum_indicum_S5.653733.4.PATRIC.fa |
| Desulfurobacterium_thermolithotrophum.DSM11699.868864.3.PATRIC.fa |
| Dictyoglomus_thermophilum.H-6-12.309799.4.PATRIC.fa |
| Elusimicrobium_minutum.Pei191.445932.6.PATRIC.fa |

| |
|---|
| Fervidicoccus_fontis_kam940.ASM25842v1.pep.all.fa |
| Fervidobacterium_nodosum_rt17_b1.ASM1754v1.pep.all.fa |
| Gemmatimonas_aurantiaca_t_27.ASM1030v1.pep.all.fa |
| Gemmatirosa_kalamazoonesis.ASM52298v1.pep.all.fa |
| Geoglobus_ahangari.strain234.113653.22.PATRIC.fa |
| Hadesarchea.1776334.3.PATRIC.faa |
| Halalkalicoccus_jeotgali.B3.795797.18.PATRIC.fa |
| Halobacteriovorax_marinus_sj.ASM21091v2.pep.all.fa |
| Haloquadratum_walsbyi.C23.768065.8.PATRIC.fa |
| Heimdalarchaeote_AB_125.fa |
| Ilyobacter_polytropus.DSM2926.572544.3.PATRIC.fa |
| Lentisphaera_araneosa_htcc2155.ASM17075v1.pep.all.fa |
| Lokiarch.1538547.4.PATRIC.fa |
| Mariprofundus_ferrooxydans.ASM127361v1.pep.all.fa |
| Melioribacter_roseus_p3m_2.ASM27914v1.pep.all.fa |
| Methanobacterium_formicicum.DSM1535.pep.all.fa |
| Methanocaldococcus_jannaschii_dsm_2661.ASM9166v1.pep.all.fa |
| Methanopyrus_kandleri_av19.ASM718v1.pep.all.fa |
| Methanosarcina_siciliae.HI350.1434119.4.PATRIC.fa |
| Nanoarchaeota_archaeon.7A1577684.3.PATRIC.fa |
| Nanoarchaeum_equitans.Kin4-M228908.8.PATRIC.fa |
| Nitrosopumilus_maritimus.SCM1.436308.17.PATRIC.fa |
| Nostoc_azollae.551115.6.PATRIC.fa |
| Odinarchaeote_LCB_4.fa |
| Palaeococcus_ferrophilus.DSM13482.588319.7.PATRIC.fa |
| Planctopirus_limnophila_dsm_3776.ASM9210v1.pep.all.fa |
| Prosthecochloris_aestuarii_dsm_271.ASM2062v1.pep.all.fa |
| Rubrobacter_xylanophilus_dsm_9941.ASM1418v1.pep.all.fa |
| Thermanaerovibrio_acidaminovorans.DSM6589.525903.6.PATRIC.fa |
| Thermodesulfatator_indicus_dsm_15286.ASM21779v1.pep.all.fa |
| Thermodesulfovibrio_yellowstonii.DSM11347.289376.4.PATRIC.fa |
| Thermosphaera_aggregans_dsm_11486.ASM9218v1.pep.all.fa |
| Thorarchaeote_AB_25.fa |
| Ureaplasma_parvum_serovar_3.ASM82873v1.pep.all.fa |
| actinobacillus_muris.ASM103820v1.pep.all.fa |
| archaeon_GW2011_AR10.1579370.3.PATRIC.fa |
| clostridium_sordellii.ATCC9714_.pep.all.fa |
| haemophilus_ducreyi.ASM104335v1.pep.all.fa |
| halophilic_archaeon.DL31.756883.5.PATRIC.fa |

Table S1. List of Genomes used in LECA reconstruction.
