## Supplemental Table S2 for "Reconstructing and Analysing The Genome of The Last Eukaryote Common Ancestor to Better Understand the Transition from FECA to LECA"

## A.

| Pathway | Function | Protein family/type | Gene/s |
|---|---|---|---|
| **Autophagy animal (HS)** | Membrane trafficking | Rab associated proteins | PIK3R1, |
| | | Mitochondrial quality control factors | ATG2$^{at,sc}$/3-4$^{at,dd,sc}$,ATG8-9$^{at,dd,sc}$/10$^{at}$,ATG12$^{at,sc}$,ATG13$^{at}$, ATG18$^{at}$, ATG101, BECN$^{at,dd}$, SH3GLB1 |
| | | SNARE | STX17, VAMP8, SNAP29 |
| | Phosphoprotein phosphatases | Ligand-gated ion channel | ITPR1 |
| | | Protein Tyrosine phosphatases | MTMR14 |
| | DNA repair and recombination proteins | | HMGB1 |
| | Signal transduction | Peptidase | CTSD |
| **Apoptosis (HS)** | Signal transduction | Peptidase | CTSD |
| | Membrane trafficking | Phosphoprotein phosphatases | ITPR1-3 |
| | | Phosphoinositide-3-kinase regulatory complex | PIK3R1 |
| | | TNF receptor-associated factor | TRAF1 |
| | Ubiquitin system | Single Ring-finger type E3 | BIRC2, TRAF2 |
| | Transmembrane tubule forming | Perforin | PRF1 |
| **Cell cycle (HS)** | DNA replication proteins | DNA Replication Initiation Factors | ORC2-5 |
| | Ubiquitin system | Multi subunit Ring-finger type E3 | SKP1 |
| | Protein phosphatases and associated proteins | Protein Tyrosine phosphatases (PTPs) | CDC25A-C, CDC45 |
| | Transcription factors | Phosphoprotein phosphatases | Rb1, RBL1, RBL2 |
| | Transcription factors | Helix-turn-Helix | E2F1-5, DP1, DP2 |
| | Chromosome and associated proteins | SAC (spindle assembly checkpoint) factors | MAD2L2, MAD2 |
| | Chromosome and associated proteins | Sister chromatid cohesion proteins | STAG1_2,SCC1 |

| | | | |
|---|---|---|---|
| | Chromosome and associated proteins | Cyclins | CCNA, CCNB1/2,CCND1-3, CCNE, CCNH |
| | Chromosome and associated proteins | Cysteine Peptidases | ESP1 |
| | DNA repair and recombination proteins | Other check point factors | YWHAB/Q/Z, YWHAE, YWHAG/H |
| | Cell cycle | p53 signalling | SFN |
| **Cellular senescence (HS)** | Chromosome and associated proteins | Cyclins | CCNA, CCNB1-3, CCND2/3, CCNE |
| | DNA replication proteins | DNA Replication Termination Factors | NBN |
| | DNA repair and recombination proteins | Check point factors | RAD9A/B, HRAD1 |
| | Membrane trafficking | Rab associated proteins | PIK3R1/2/3 |
| | Messenger RNA biogenesis | mRNA surveillance and transport factors | ZFP36L |
| | Mitochondrial biogenesis | SLC25: Mitochondrial carrier | SLC25A4S |
| | Mitochondrial protein import machinery | Porin | VDAC1-3 |
| | Protein phosphatases and associated proteins | Protein Tyrosine phosphatases (PTPs) | CDC25A |
| | Phosphoprotein phosphatases | Ligand-gated ion channel | ITPR1-3 |
| | Transcription factors | Phosphoprotein phosphatases | Rb1, RBL1, RBL2 |
| | | Helix-turn-Helix | E2F1-5 |
| | | | LIN54 |
| | | | LIN9 |
| **Endocytosis (HS)** | Ubiquitin system | Single Ring-finger type E3 | CBL, CBLB, CBLC |
| | Cytoskeleton proteins | Actin-binding proteins | CIN85, ARPC1A[sc]/B[sc], ARPC2-5[at,dd,sc], CAPZA[at,dd,sc], CAPZB[at,dd,sc], AMPH[sc] |
| | Membrane Trafficking | GTPase | DNM[at], RBSN[sc], rab11fip125, VPS26[at,sc], VPS35[at,dd,sc], VPS45[at,dd,sc] |
| | | Sorting nexins | SNX1-2[at,dd,sc]/3[sc], SNX6, SNX12[sc], SNX32 |
| | | BAR family proteins | SH3GLB1, SH3GLB2, SH3GL, AMPH[sc] |

| | | | |
|---|---|---|---|
| | | AP-2 complex subunits | AP2M1[dd,sc], AP2S1[dd,sc], AP2A[at,dd,sc] |
| | | ESCRT complexes | TSG101[at,dd,sc], CHMP7 |
| | | WASH complex | CCDC53, SPG8 |
| | | | CLTC[dd,sc] |
| | | | SPG20[at] |
| | | | IST1[sc] |
| **Lysosome (HS)** | ATP synthesis | V-type ATPase | ATP6D/H |
| | Peptidases | Aspartic peptidases | NAPSA |
| | | Cysteine Peptidases | LGMN |
| | Hydrolases | Glycosylase | GLA,NAGA |
| | | Sterol esterase | LIPA |
| | | lysophospholipase | LYPLA3 |
| | | Phosphorus-containing anhydrides | ENTPD4 |
| | Tranferase | | GNPTAB |
| | Lectins | P-type lectin | GNPTG |
| | Membrane trafficking | AP-1/3 complex | AP1M, AP1S1-3, AP1G1, AP3M, AP3S, AP3D, AP4S1, AP4E1, AP4M1 |
| | | Others | GGA |
| | | Clathrin | CLTC |
| | Lysosome | Protein families: signalling and cellular | GM2A |
| | | | CTN5, BTS, LITAF |
| | | Transporters | SORT1 |
| **Meiosis – yeast (SC)** | DNA replication proteins | DNA Replication Initiation Factors | ORC2-5 |
| | DNA repair and recombination proteins | Other check point factors | MEC1 |
| | Chromosome and associated proteins | Sister chromatid cohesion proteins | STAG1/2 |
| | | SAC (spindle assembly checkpoint) factors | MAD1/2 |

| | | | |
|---|---|---|---|
| | | Cyclins | CLN2/3,CLB1-4 |
| | Transcription factors | Helix-turn-Helix | MATA2 |
| | Phosphoprotein phosphatases (PPPs) | Other centromeric chromatin formation proteins | PPP2R1/5 |
| | Cysteine Peptidases | Sister chromatid separation proteins | ESP1 |
| | Ubiquitin system | Multi subunit Ring-finger type E3 | APC2 |
| | | | HOP1 |
| **Necroptosis (HS)** | Ubiquitin system | Single Ring-finger type E3 | TRAF2/5,BIRC2 |
| | Ubiquitin-specific proteases (UBPs) | Cysteine Peptidases | CYLD, TNFAIP3 |
| | | toll-like | FAF1 |
| | | TRIF-related | TRIF |
| | Other mitochondrial DNA transcription and translation factors | SLC25: Mitochondrial carrier | SLC25A4S |
| | Clathrin-mediated endocytosis | Dynamin | DNM1L |
| | Membrane trafficking | ESCRT | VPS24, CHMP7 |
| | Mitochondrial protein import machinery | Porin | VDAC1-3 |
| **Oocyte meiosis (HS)** | Chromosome and associated proteins | SAC (spindle assembly checkpoint) factors | MAD2B, MAD2 |
| | | Cyclins | CCNB1/2, CCNE |
| | | Sister chromatid cohesion proteins | STAG3 |
| | Phosphoprotein phosphatases (PPPs) | Other centromeric chromatin formation proteins | PPP2R5 |
| | | Ligand-gated ion channel | ITPR1-3 |
| | Protein phosphatases and associated proteins | Protein Tyrosine phosphatases (PTPs) | CDC25C |
| | Ubiquitin system | Multi subunit Ring-finger type E3 | APC2, SKP1 |
| | Messenger RNA biogenesis | mRNA surveillance and transport factors | CPEB |

| | | | |
|---|---|---|---|
| | DNA repair and recombination proteins | Other check point factors | YWHAB/Q/Z, YWHAE, YWHAG/H |
| | Peptidases | Cysteine Peptidases | ESP1 |
| **Phagosome (HS)** | Membrane trafficking | SNARE | STX7/12/13/18[at], VAMP3 |
| | | NADPH oxidase complex | NCF1/4 |
| | Cytoskeleton proteins | Tubulin-binding proteins | DYNC1I/DYNC2H |
| | ATP synthesis | V-type ATPases | ATP6C[at]/D/G[at]/H, ATPeV1H |
| | Protein export | Secretion system | SEC61B[at] |
| | Chaperones and folding catalysts | Lectins | CALR[at], CANX[at] |
| **Peroxisome (HS)** | Ubiquitin system | Single Ring-finger type E3 | PEX3[dd,sc]/10-13[dd,sc] |
| | Oxidoreductases | Hydroxylase | ACOX2 |
| | Acyltransferases | Carnitine 0-octanoyltransferase | CROT |
| | Hydrolases | Acyl-CoA thioesterase | ACOT8 |
| | Peroxidases | Peroxiredoxin | PRDX5 |
| | Pore-forming | Peroxisomal membrane protein | PXMP2/4 |
| | | Mitochondrial inner membrane protein | MPV17[sc], MPV17L |
| **Regulation of actin cytoskeleton (HS)** | Chaperones and folding catalysts | Lectins | CALR, CANX |
| | ATP synthesis | V-type ATPases | ATP6C, ATP6D, ATP6G, ATP6H, ATPeV1H |
| | Chromosome and associated proteins | SAC (spindle assembly checkpoint) factors | DYNC1I, DYNC1H |
| | Protein phosphatases and associated proteins | Phosphoprotein phosphatases (PPPs) | APC |
| | Cytoskeleton proteins | Actin-binding proteins | ARPC1/B, ARPC2-5, CFL, NCKAP1, WASF2 |
| | Signal transduction | | CRK |
| | Membrane trafficking | Cytoskeleton proteins | PXN |
| | | Rho guanine nucleotide exchange factors | ARHGEF1 |
| | | Rab associated proteins | PIK3R1 |

## B.

| Pathway | Function | Protein family/type | Gene/s |
|---|---|---|---|
| **Base excision repair (HS)** | DNA replication proteins | DNA polymerase epsilon complex | POLE2[at,dd,sc]/4[sc], POLD2[at,sc]/3-4[at] |
| | Chromosome and associated proteins | Nucleosome assembly factors | HMGB1[at] |
| | | Heterochromatin formation proteins | MBD4 |
| **RNA transport (HS)** | Transfer RNA biogenesis | tRNA export factors | XPO1[at,dd,sc]/5[at,dd,sc] |
| | Spliceosome | SMN complex factors | GEMIN2[dd] |
| | | Other proteins involved in snRNP biogenesis | SNUPN |
| | Ribosome biogenesis | Export and cytoplasmic maturation factors | KPNB1[sc] |
| | Messenger RNA biogenesis | Transport factors | MAGOH[at,dd], SAP18[at], SRRM1[dd], NXF[sc], NXT1/2, NUP153/93[dd,sc]/54[sc]/50/155[at,dd,sc]/98[at,dd,sc]/160[at] /107[dd]/210[at,dd]/214, SENP2, POM121, SUMO[at,dd,sc], THOC5[at,dd]/7[at,dd], PAIP1 |
| | Translation factors | Initiation factors | EIF1[dd,sc], EIF3C[at,dd,sc]/J[dd]/E[at,dd], EIF4E[at,dd,sc]/G[at,dd,sc], EIF2B5[dd,sc]/3[dd,sc], FMR |
| | Messenger RNA biogenesis | Surveillance factors | UPF2[sc] |
| **DNA replication (HS)** | DNA replication proteins | DNA Replication Elongation Factors | RNASEH2B/C |
| | DNA replication proteins | DNA polymerase delta complex | POLD2[at,dd,sc]/3-4[at] |
| | | DNA polymerase epsilon complex | POLE2[at,sc]/4[sc] |
| | | DNA polymerase alpha / primase complex | POLA2[at,dd,sc], PRI2[at,dd,sc] |
| | | DNA Replication Elongation Factors | RPA2[at,sc]/4 |
| **Ribosome (HS)** | Ribosome | Large subunit | RP-L27[dd,sc]/28[dd]/29[dd,sc]/36e[dd,sc],RP-LP1[dd,sc]/2[dd,sc] |
| | | Small subunit | RP-S7e[dd,sc], RP-S10e[at,dd,sc] |
| **mRNA surveillance pathway (HS)** | Messenger RNA biogenesis | Transport factors | MAGOH[at,dd], SAP18[at], SRRM1[dd], NXF[sc], NXT1/2 |
| | Messenger RNA biogenesis | Surveillance factors | UPF2[sc], SMG5-6/7[at] |
| | | 5'processing factors | RNMT[sc] |
| | | 3'processing factors | PCF11[at,sc], CPSF1[at,dd,sc]/2/3[at,dd,sc]/4[dd,sc]/5[at]/6/7, FIP1L1, SYMPK[at,dd] |

| | | | |
|---|---|---|---|
| | Chromosome and associated proteins | Other centromeric chromatin formation proteins | PPP2R3[at]/5[at,dd,sc] |
| **Mismatch repair (HS)** | DNA replication proteins | DNA polymerase delta complex | POLD2[at,dd,sc]/3-4[at] |
| | | DNA Replication Elongation Factors | RFA2[at,sc], RPA4 |
| **RNA polymerase (HS)** | Transcription machinery | Pol II specific subunits | RPB4[at,dd,sc] |
| | | Pol I, II, III common subunits | RPB8[at,dd,sc] |
| | | Pol III specific subunits | RPC3[dd,sc]/4[sc]/6[dd,sc] |
| **Nucleotide excision repair (HS)** | DNA repair and recombination proteins | Basal transcription factors | TFIIH1/2[at]/3[dd,sc], TFIIH1, TTDA[dd,sc] |
| | Ubiquitin system | Multi subunit Ring-finger type E3 | DDB1[at,dd], CUL4[at,dd] |
| | DNA repair and recombination proteins | Other nucleotide excision repair factors | ERCC1[at,dd,sc], XPC[at,dd,sc] |
| | Cell cycle | Cyclins | CCNH[sc] |
| | DNA replication proteins | DNA polymerase delta complex | POLD2[at,dd,sc]/3-4[at] |
| | | DNA polymerase alpha / primase complex | POLA2, PRI2 |
| | | DNA polymerase epsilon complex | POLE2[at,sc]/4[sc] |
| | | DNA Replication Elongation Factors | RFA2[sc], RPA4[sc] |
| **Ribosome biogenesis in eukaryotes (HS)** | Transfer RNA biogenesis | tRNA export factors | XPO1[at,dd,sc] |
| | Ribosome biogenesis | Other 90S particles | NAT10[dd,sc] |
| | | GTPase | BMS1[dd,sc] |
| | | U3 small nucleolar RNA-associated proteins | MPP10[sc], UTP6[dd,sc], UTP22[dd,sc] |
| | | Kinase | CSNK2B[at,dd,sc] |
| | Messenger RNA biogenesis | Transport factors | NXF[sc], NXT1[sc]/2[sc] |
| **Homologous recombination (HS)** | DNA repair and recombination proteins | Other homologous recombination factors | MUS81[dd,sc], EME1 |
| | | BRCA1-A complex | FAM175A |
| | Ubiquitin system | Multi subunit Ring-finger type E3 | BRCA2 |
| | DNA replication proteins | DNA polymerase delta complex | POLD2[dd,sc]/3-4[sc] |

| | | | |
|---|---|---|---|
| | | Other telomere regulation proteins | NBN |
| | | DNA Replication Elongation Factors | RFA2[sc], RPA4 |
| **Basal transcription factors (HS)** | Transcription machinery | Basal transcription factors | TAF1[at,dd]/2[at,dd]/6[at,dd]/7[at]/9/9B[at]/10[at]/11[at,dd]/12[at]/13[at,dd] /14-15,TFIIA1[dd], TFIIE2[dd], TFIIH1[dd]/2[at,dd]/3-4[dd], TTDA[dd] |
| | Cell cycle | cyclin | CCNH |
| **Non-homologous end-joining (HS)** | DNA repair and recombination proteins | DNA-PK complex | XRCC5[at,dd,sc]/XRCC6[at,dd,sc] |
| | | DNA Ligase 4 complex | XRCC4 |
| **Spliceosome (HS)** | Spliceosome Complex A/B | U1 snRNP specific factors | SNRPA[at,sc], SNRPC[dd] |
| | | U1 related factors | PRPF40[at,sc] |
| | Spliceosome Complex A/B/C | U2 snRNP specific factors | SNRPB2[at,dd,sc], SF3A3[at,dd], SF3B1[dd,sc]/3[at,dd]/5[dd], PHF5A[dd,sc] |
| | | U2 related factors | RBM17[at,sc] |
| | Spliceosome Complex B/C | Prp19-CDC5 complex | CDC5L[at,dd,sc], CWC15[at,dd], PQBP1 |
| | | Prp19-related factors | SNW1[at,dd,sc], ISY1[dd,sc], BUD31[dd,sc] |
| | | U5 SnRNP specific factors | PRPF6[at,dd,sc]/8[at,dd,sc], TXNL4A[at,dd,sc] |
| | | U4/U6.U5 tri-SnRNP related factors | PRPF38B[at,dd], SNRNP27, SNU23sc |
| | Spliceosome Complex C | mRNA surveillance and transport factors | MAGOH[at,dd,sc] |
| | | Step II factors | SLU7[at,dd,sc], PRPF18[dd,sc] |
| | | Common spliceosomal components | HNRNPK, PCBP1, |
| | Ubiquitin system | Single Ring-finger type E3 | SART1[sc] |

## C.

| Pathway | Function | Protein family/type | Gene/s |
|---|---|---|---|
| **Mucin type O-glycan biosynthesis (HS)** | Glycan metabolism | O-glycan biosynthesis, mucin type core | GCNT1/4, B3GNT6, C1GALT2 |
| **Ubiquitin mediated proteolysis (HS)** | Ubiquitin system | Multi subunit Ring-finger type E3 | ELOC[sc], FBXO2, APC2[at,dd,sc], CUL1[at,dd,sc]/2/3[at,dd,sc]/4[at,dd]/5, SKP1[at,dd,sc], DDB1[at,dd] |
| | | Single Ring-finger type E3 | FANCL, BIRC2/3, RCHY1, SIAH1[at], CBLC, CBL, CBLB |
| | | UBL E3 ligases | PIAS1[sc]/2-4 |
| | | U-box type E3 | UBOX5, UBE4A/B[dd,sc], UBE2Q, |
| **N-Glycan biosynthesis (HS)** | Glycan metabolism | N-Glycan biosynthesis | DPM2[at]/3, FUT8 |
| | | N-glycan precursor biosynthesis | ALG2[sc]/11[at,sc]/12[sc] |
| | | N-glycosylation by oligosaccharyltransferase | OST1[sc]/2[at,sc], WBP1[at,sc] |
| **Mannose type O-glycan biosynthesis (HS)** | Glycosyltransferases | O-Glycan | POMGNT2, LARGE, B4GAT1 |
| | | Wide specificity | B3GALNT2 |
| **Other types of O-glycan biosynthesis (HS)** | Glycosyltransferases | O-Glycan | EOGT |
| **Proteasome (HS)** | Proteasome regulatory particles | non-ATPase subunits | PSMD1[dd,sc]/2[at,dd,sc]/3/4[at,dd,sc]/6[dd,sc]/8[at,dd,sc]/11[at,dd,sc]/12[at,dd,sc]/13[dd,sc], RPN13[dd], PSME4[dd] |
| **Protein processing in endoplasmic reticulum (HS)** | Ubiquitin system | Single Ring-finger type E3 | TRAF2, MARCH6, RNF5 |
| | Chaperones and folding catalysts | Lectins | CALR[at,dd,sc], CANX[at,dd,sc], PRKCSH[at,dd], |
| | | Others | HSPBP1[at] |
| | Protein export | Secretion system | SEC61B[at,dd] |
| | Glycan metabolism | N-glycosylation by oligosaccharyltransferase | OST1/2[at], WBP1[at] |
| | Protein processing | COPII complex | SEC23[dd,sc] |
| | | HRD1/SEL1 ERAD complex | UFD1[at,dd], NPLOC4[at,dd], DERL2[at,dd]/3[at,dd], HERPUD1 |
| | Glycosyltransferases | Quality control factors | HUGT[dd] |

| | | | |
|---|---|---|---|
| | Transcription factors | Basic leucine zipper (bZIP) | XBP1 |
| | Ubiquitin system | U-box type E3 | UBE4B[dd] |
| | | Ubiquitin-specific proteases (UBPs) | ATXN3 |
| | | Multi subunit Ring-finger type E3 | FBXO2/6, CUL1[at,dd], SKP1[at,dd] |
| | Protein processing | | UBX1at |
| | | Heat-shock proteins | CRYAA/B |
| | Mitochondrial biogenesis | Mitochondrial quality control factors | SIL1 |
| | Protein processing | Translocon-associated protein (TRAP) complex | SSR2[dd]/3-4 |
| **Protein Export (HS)** | Protein export | Secretion system | SRP68[dd,sc]/9[dd],SPCS1[dd,sc]/3[dd,sc]SEC61B[at,dd] |
| **SNARE Interactions (HS)** | Membrane trafficking | SNARE | BET1[at], YKT6[at,dd,sc], GOS1[at,sc], STX2[at,sc]/3[at,sc]/5[at,dd,sc]/6[dd]/7/8[dd]/11/16[at,dd,sc]/17/18[at,sc], VAMP1-3/4[sc]/5/7[at,dd]/8, SNAP23/29[at], SEC22[at,sc], VTI1[at,dd,sc] |
| **Sulfur relay system (hs)** | Ubiquitin system | Ubiquitin-like proteins (UBLs) | URM1[sc] |
| | tRNA modfication factors | Thiolation factors | CTU2[dd,sc] |
| **RNA degradation (hs)** | Messenger RNA biogenesis | mRNA degradation factors | C1D |
| | RNA processing | TRAMP complex | PAPD5/7 |
| | Messenger RNA biogenesis | mRNA degradation factors | CNOT1[at,dd,sc]/2[at,sc]/7[at,sc]/8[at,sc]/9[sc]/10, PARN[at,sc] |
| | | Other 3'-5' decay factors | DCPS[sc] |
| | | mRNA surveillance and transport factors | PATL1[sc] |
| **Various types of N-glycan biosynthesis (sce)** | Glycan metabolism | N-Glycan biosynthesis, high-mannose type | MNN10/11, HOC1 |
| | Glycosyltransferases | Glycan extension factors | OCH1 |
| | | N-glycan precursor biosynthesis | ALG2/11-13 |
| | Glycan metabolism | N-glycosylation by oligosaccharyltransferase | OST1/2, WBP1 |

**D.**

| Pathway | Function | Protein family/type | Gene/s |
|---|---|---|---|
| **Signaling pathways regulating pluripotency of stem cells (HS)** | Cytoskeleton proteins | Tubulin-binding proteins | APC |
| | Membrane trafficking | Rab GTPases and associated proteins | PIK3R1-3 |
| | Transcription factors | Helix-turn-helix | NANOG, POU5F, HESX1, PAX6, MEIS1, HOX1, LHX5, OTX1, DLX5, ISL1 |
| | | beta-Scaffold factors with minor groove contacts | SOX2 |
| | DNA replication proteins | DNA Replication Termination Factors | RIF1 |
| | | One cut domain, family member 1 | HNF6 |
| **Gap Junction (HS)** | Ligand-gated channels | Phosphoprotein phosphatases (PPPs) | ITPR1-3 |
| **p53 signaling pathway (HS)** | Protein families: signaling and cellular processes | - | SFN |
| | DNA replication proteins | CDK (cyclin dependent kinase) | CCNE |
| | DNA repair and recombination proteins | Other factors with a suspected DNA repair function | RRM2 |
| | Ubiquitin system | Single Ring-finger type E3 | RCHY1 |
| | Cell cycle | Cyclins | CCND1-3, CCNG1, CCNB1-2 |
| **Hedgehog signalling pathway (HS)** | Ubiquitin system | Multi subunit Ring-finger type E3 | CUL1/3 |
| | Cell cycle | Cyclins | CCND1/2 |
| **Phosphatidylinositol signaling system (HS)** | Ligand-gated channels | Phosphoprotein phosphatases (PPPs) | ITPR1-3 |
| | Membrane trafficking | Rab GTPases and associated proteins | PIK3R1-3 |
| | Membrane trafficking | Actin-binding proteins | PI4K2 |

| | | | |
|---|---|---|---|
| | Signal transduction | Hydrolases | INPP4 |
| | Signal transduction | Transferases | IPMKat |
| | Inositol phosphate metabolism, Ins(1,3,4,5)P4 => Ins(1,3,4)P3 => myo-inositol | Hydrolases | inositol-1,4,5-trisphosphate 5-phosphatase |
| | Inositol phosphate metabolism, PI=> PIP2 => Ins(1,4,5)P3 => Ins(1,3,4,5)P4 | Transferases | ITPK |
| | Protein phosphatases and associated proteins | Hydrolases | MTMR14 |
| | Signal transduction | Transferases | IP6K, PPIP5K |
| | Signal transduction | Hydrolases | SAC1[at,dd]/2 |
| **Adherens junction(hs)** | Transcription factors | beta-Scaffold factors with minor groove contacts | LEF1, TCF7, TCF7L2, TCF7L1 |
| | Ribosome biogenesis | Kinase | CSNK2B |
| | Cytoskeleton proteins | Actin-binding proteins | WASF2/3 |
| | Signal transduction | - | SORBS1 |
| **Notch signaling pathway(hs)** | Signal transduction | Notch signaling pathway | APH1A, NCOR2, SNW1 |
| | Peptidases | Aspartic Peptidases | PSEN1/2 |
| | Ubiquitin system | Single Ring-finger type E3 | DTX |
| **Focal adhesion(hs)** | Membrane trafficking | Rab GTPases and associated proteins | PIK3R1-3 |
| | - | - | ZYX, PXN |
| | Signal transduction | Hydrolases | RELN |
| | Ubiquitin system | Single Ring-finger type E3 | BIRC2/3 |
| | Cell cycle | cyclin | CCND1-3 |
| | Signal transduction | | SHC1-3, CRK |
| **Calcium signaling pathway(hs)** | Inositol phosphate metabolism, PI=> PIP2 => Ins(1,4,5)P3 => Ins(1,3,4,5)P4 | Transferases | ITPK |

| | | | |
|---|---|---|---|
| | Mitochondrial protein import machinery | Porin | VDAC1-3 |
| | Ligand-gated channels | Phosphoprotein phosphatases (PPPs) | ITPR1-3 |
| | Mitochondrial biogenesis | SLC25: Mitochondrial carrier | SLC25A4S |
| | | Mitochondrial calcium uniporter complex | MCU |
| **Tight junction(hs)** | Signal transduction | - | PRKAG, PRKAB |
| | Cell cycle | cyclin | CCND1 |
| | Transcription machinery | Transcription termination factor | SYMPK |
| | Cytoskeleton proteins | Actin-binding proteins | CTTN |
| | - | - | CRB3 |
| **Hippo signaling pathway(hs)** | Signal transduction | - | WWTR1 |
| | DNA repair and recombination proteins | Other check point factors | YWHAB/Q/Z/E/G/H |
| | Signal transduction | - | MOB1 |
| | Messenger RNA biogenesis | mRNA cycle factors | AJUBA |
| | Cytoskeleton proteins | Tubulin-binding proteins | APC |
| | Transcription factors | beta-Scaffold factors with minor groove contacts | LEF1, TCF7, TCF7L2, TCF7L1, SOX2 |
| | Cell cycle | Cyclins | CCND1-3 |
| | Membrane trafficking | Rab associated proteins | CALM |
| | Signal transduction | Hydrolases | SAC1 |
| **Apelin signaling pathway(hs)** | Signal transduction | - | PRKAG, PRKAB, UCP1 |
| | Mitochondrial biogenesis | Mitochondrial transcription factors | TFAM |
| | Cell cycle | cyclin | CCND1 |
| | Transcription factors | beta-Scaffold factors with minor groove contacts | MEF2A/B/C/D |
| | Membrane trafficking | Mitochondrial quality control | BECN, ATG8 |

| | | | |
|---|---|---|---|
| | | factors | |
| **Cell adhesion molecules (CAMs) (hs)** | Signaling molecules and interaction | Cell adhesion molecules (CAMs) | VTCN1 |
| **Wnt signaling pathway (HS)** | Signal transduction | - | PRICKLE |
| | Peptidases | Metalloendopeptidases | MMP7 |
| | Cell cycle | Cyclins | CCND1-3 |
| | Transcription factors | beta-Scaffold factors with minor groove contacts | LEF1, SOX17, TCF7, TCF6L1 |
| | Ubiquitin system | Multi subunit Ring-finger type E3 | CUL1, SKP1 |
| | Cytoskeleton proteins | Tubulin-binding proteins | APC |
| | Ubiquitin system | Single Ring-finger type E3 | SIAH1 |
| | Peptidases | Aspartic Peptidases | PSEN1 |
| | Messenger RNA biogenesis | Nuclear pore complex | SENP2 |
| | Ribosome biogenesis | Kinase | CSNK2B |
| | Signal transduction | Transferases | PORCN |
| **MAPK signaling pathway (hs)** | Signal transduction | - | LAMTOR3 |
| | DNA repair and recombination proteins | Other check point factors | CDC25B |
| | Transcription factors | beta-Scaffold factors with minor groove contacts | MEF2C, SRF |
| | Chaperones and folding catalysts | Heat shock proteins | HSPB1 |
| | Signal transduction | - | MAP3K7IP1, CRK |
| | Ubiquitin system | Single Ring-finger type E3 | TRAF2 |
| **MAPK signaling pathway – plant (AT)** | Transcription factors | Basic leucine zipper (bZIP) | VIP1 |
| | Signal transduction | - | TMEM22 |
| **MAPK signaling pathway – yeast (SC)** | Transcription factors | beta-Scaffold factors with minor groove contacts | RLM1, MCM1, SMP1 |
| | Glycosyltransferases | Structural polysaccharide | 1,3-beta-glucan synthase |

| | | | |
|---|---|---|---|
| | Signal transduction | - | BEM1, RGA1/2, ROM1/2 |
| | Cell cycle | Cyclins | CLN1/2, CLB1/2, MIH1 |
| | DNA repair and recombination proteins | Other check point factors | YWAE |
| **Ras signaling pathway (HS)** | Membrane trafficking | Rab GTPases and associated proteins | PIK3R1-3 |
| | Signal transduction | | SHC1-3 |

## E.

| Pathway | Function | Protein family/type | Gene/s |
|---|---|---|---|
| **Fatty acid elongation (HS)** | Fatty acid biosynthesis, elongation, endoplasmic reticulum | Hydrolyases | PHS1[at] |
| | Fatty acid biosynthesis, elongation, endoplasmic reticulum | Acyltransferases | ELOVL1/2[sc]/3[sc]/5-7 |
| **Glycerophospholipid metabolism (HS)** | Lipid biosynthesis proteins | Phospholipid acyltransferase | LPCAT3/4, LCLAT1, AGPAT3-4[at]/5, LPGAT1 |
| | Glycerophospholipid metabolism | Transferases | CEPT1, CHPT1[sc], CHK[at], CHAT, ETNK[at,dd,sc], LCAT[at] |
| | | Hydrolases | PLA2G16, LPIN[at,sc], NTE, LYPLA3 |
| | Triacylglycerol biosynthesis | Transferases | MBOAT1/2, PTDSS1 |
| **Primary bile acid biosynthesis (HS)** | Sterol biosynthesis | oxidoreductases | ACOX2, CH25H |
| | | Hydrolases | ACOT8 |
| **Fatty acid degradation (HS)** | - | oxidoreductases | ADH1[at]/4/6/7 |
| | - | Transferases | CPT2 |
| | Mitochondrial biogenesis | Transferases | CPT1B/C |
| **Steroid hormone biosynthesis (HS)** | - | oxidoreductases | SRD5A3 |
| | Glycosyltransferases | Hydrophobic molecule | UGT |
| | - | Transferases | SULT1E1 |
| **Biosynthesis of unsaturated fatty acids (HS)** | Lipid biosynthesis proteins | Lyases | PHS1 |
| | Lipid biosynthesis proteins | Transferases | ELOVL5/6 |
| **Ether lipid metabolism (HS)** | Glycerophospholipid metabolism | Transferases | CEPT1[sc], CHPT1[sc], LPCAT4 |
| | | Hydrolases | PLA2G16 |
| | Glycosyltransferases | Glycosphingolipid | CGT |
| **Sphingolipid metabolism (HS)** | Sphingosine biosynthesis | Transferases | CER5[dd], SGMS |
| | | Hydrolases | SMPD2, GLA |
| | Glycosyltransferases | Glycosphingolipid | CGT |

| | | | |
|---|---|---|---|
| **Linoleic acid metabolism (HS)** | Glycosyltransferases | Hydrolases | PLA2G16 |
| **Arachidonic acid metabolism (HS)** | Glycosyltransferases | Hydrolases | PLA2G16 |
| | | Lyases | LTC45 |
| | | Isomerases | PTGES3 |
| **Glycerolipid metabolism (HS)** | Acylglycerol degradation | Hydrolases | PNPLA2/3, lIPF |
| | Triacylglycerol biosynthesis | Transferases | TGL3[sc]/4[at,sc], MOGAT2[dd,sc]/3, DGAT1[at,dd]/2[at], AGPAT3-5 |
| | Triacylglycerol biosynthesis | Hydrolases | LPIN[at,sc] |
| | Lipid biosynthesis proteins | Transferases | MBOAT1/2, LCLAT1[at,sc] |
| | Glycosphingolipid biosynthesis - globo and isoglobo series | Hydrolases | GLA |
| **Inositol phosphate metabolism (HS)** | Inositol phosphate metabolism, Ins(1,3,4)P3 => phytate | Transferases | IPMK[at,sc], IPPK[at] |
| | Multiple inositol-polyphosphate phosphatase | Hydrolases | MINPP1[dd] |
| | Inositol phosphate metabolism, Ins(1,3,4,5)P4 => Ins(1,3,4)P3 => myo-inositol | Hydrolases | inositol-1,4,5-trisphosphate 5-phosphatase |
| | Inositol phosphate metabolism, PI=> PIP2 => Ins(1,4,5)P3 => Ins(1,3,4,5)P4 | Transferases | ITPK |
| | Signal transduction | Hydrolases | INPP4, SAC1[at,dd,sc]/2 |
| | Membrane trafficking | Actin-binding proteins | PI4K2[sc] |
| | Protein phosphatases and associated proteins | Hydrolases | MTMR14 |
| **Steroid biosynthesis (HS)** | Cholesterol biosynthesis | Isomerases | EBP[at] |
| | | oxidoreductases | MESO1[sc] |
| | | Hydrolases | LIPA[sc] |
| | | Transferases | SOAT[sc] |
| **α-Linolenic acid metabolism (AT)** | Triacylglycerol biosynthesis | Transferases | TGL4 |

**Table S2** Gene gains during the FECA to LECA transition sorted according to pathway categories; (A) Cellular Processes, (B) Genetic Information Processing, (C) Protein Processing, (D) Signalling, and (E) Lipid Metabolism.
